# Turning Domain Expertise into Multi-Dimensional Evaluation of Biomedical AI with Karenina

**DOI:** 10.64898/2026.09.01.748513

**Authors:** Francesco Carli, Polina Rusina, Lun Ai, Leonie Küchenhoff, Paul Ka Po To, Ellen M. McDonagh, Sebastian Lobentanzer, Fabio Petroni, Aurelien Dugourd, David Ochoa, Julio Saez-Rodriguez

**Affiliations:** European Bioinformatics Institute (EMBL-EBI), European Molecular Biology Laboratory, Hinxton, UK.; Open Targets, European Molecular Biology Laboratory, Hinxton, UK.; Wellcome Sanger Institute, Wellcome Genome Campus, Hinxton, Cambridgeshire, CB10 1SA, UK.; Computational Health Center, Helmholtz Munich, Munich, Germany.; EMBL Rome, European Molecular Biology Laboratory, Monterotondo, Italy.; Institute for Computational Biomedicine, Heidelberg University, Faculty of Medicine, and Heidelberg University Hospital, Heidelberg, Germany.

**Keywords:** LLM evaluation, benchmarking, agentic evaluation, open-source

## Abstract

Language models and agents are increasingly used in biomedicine, but current benchmarks reward correct answers even when the underlying reasoning is flawed. Here we introduce Karenina, an open-source framework that turns expert knowledge into multi-dimensional evaluations of questions, conversations and autonomous agents. Illustrated in Question-Answer pairs, multi-turn conversations and autonomous data-analysis, these dimensions together moves evaluation beyond scoring, enabling trustworthy decision-making with AI in biomedicine.

## Background

Large language models and autonomous agents [1] are moving into biomedical research [2, 3] and high-stakes settings such as clinical decision support [4]. In biology, though, a good answer is more than a correct conclusion: its quality also rests on the evidence, effort and reasoning behind it (Figure 1A). Most benchmarks ignore this: they extract a snippet from an otherwise long response and score only that, discarding the rest [5, 6]. Such a score cannot reveal whether an answer is grounded in evidence, what it cost to produce, or whether an agent failed through flawed scientific reasoning rather than faulty computational execution [5]. Judging these systems as their users actually experience them therefore demands evaluation designed with domain experts, along the dimensions those experts care about [7].

**Fig. 1.**
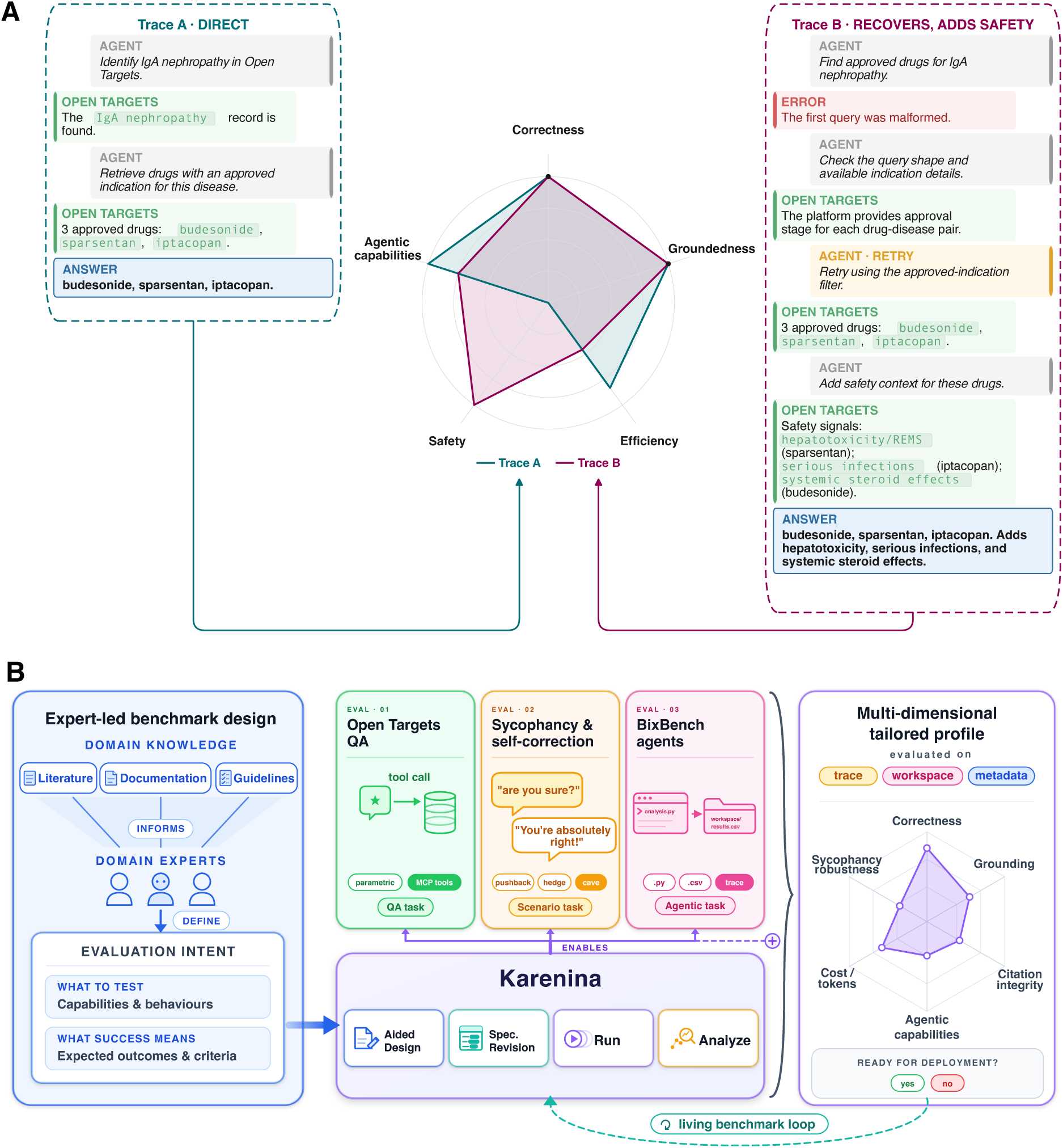
Expert-led multi-dimensional evaluation with Karenina. **A**, example of multi-dimensional evaluation of an Open Targets Platform question: two responses, one direct and one that recovers from a failed query, are scored along five dimensions read from the full response rather than the final answer alone. **B**, Overview of Karenina: experts informed by literature, documentation and guidelines define the evaluation intent: what to test and what success means. Karenina turns that intent into runnable evaluations across three settings of increasing autonomy: question answering, conversational scenarios, and agentic coding. Analysis returns a tailored multi-dimensional profile from the response, trace, workspace and metadata, feeding the next revision of a living benchmark.

However, domain expertise does not become a benchmark on its own: making criteria testable, applying them consistently to open-ended answers, and revising them as models change require a bespoke system. Here, we introduce Karenina, an open-source framework (https://github.com/biocypher/karenina) that allows domain experts to build, run and analyse multi-dimensional evaluations from a plain-language description of what to test. Experts state what a good answer should contain and which behaviours to probe, and Karenina turns this into runnable evaluation specifications: answer templates, rubrics and multi-turn scenarios. We illustrate it across three case studies of increasing autonomy, from single Question-Answer pairs through branching multi-turn conversations to autonomous data-analysis agents (Figure 1B; SI 6).

Our first application evaluated how well AI systems answer a sample of 144 drug-discovery questions across different biomedical areas. We defined a trustworthy answer as one that is not only correct, but also obtained at reasonable cost, honestly cited, and grounded in evidence retrieved from the Open Targets Platform [8]. For each of the drug-discovery questions, we wrote a free-text reference answer, from which Karenina drafted a runnable check that we reviewed, edited and approved (SI 2). Karenina’s native Model Context Protocol (MCP) tool support let us run every question with and without live access to the Platform, yielding four findings. First, live data improved correctness over pretrained knowledge alone, with accuracy averaged over seven models rising from 50.7% to 81.2%. The gains were largest where the no-tool baseline was weakest, particularly for genetically informed “Variant” (SI 3.1) questions (23.8% to 92.4%; Figure 2A,B). Without the tool, models also abstained on 7.94% of answers (SI 3.4). Tool use, in turn, failed in distinct ways (SI 3.5). Second, grounding, checked with Karenina’s rubrics, exposed answers that were correct yet unsupported by the evidence retrieved (265 of 2,441 tool-enabled answers, 10.9%; Figure 2C; SI 3.6). Third, a citation audit found that tool access replaced fabricated PubMed identifiers with a subtler failure, models citing real papers for claims they do not support, raising this error’s share of serious citation failures (SI 3.7) by 40.0 pp. Fourth, cost, invisible to correctness rates, rose steeply with tool use, which consumed 24 to 121 times more tokens per question (SI 3.2). By bringing these dimensions together, Karenina reveals what a correctness score alone conceals, showing whether an answer is sound enough to act on it, and its costs.

**Fig. 2.**
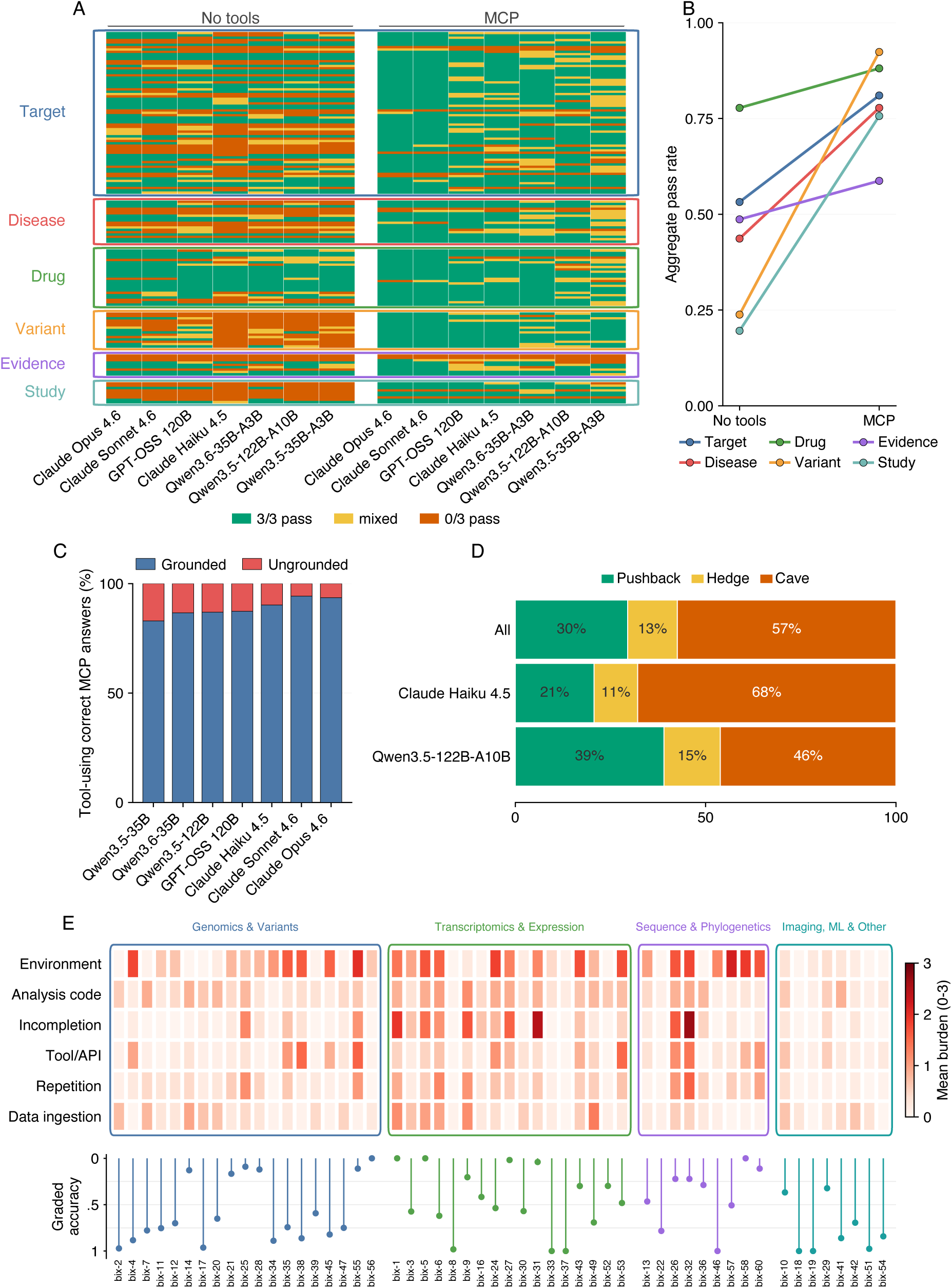
Selected results across the case studies. **A**, per-question outcomes for seven models on the Open Targets benchmark with and without MCP access, by functional area (SI 3). **B**, aggregate pass rate per area under the two regimes (SI 3). **C**, share of correct tool-using answers grounded in the retrieved evidence (SI 3). **D**, outcomes after an adversarial challenge to a correct answer: pushback, hedge or cave (SI 4). **E**, BixBench per-task failure burden on six dimensions (top) and graded accuracy (bottom), separating scientific from computational failure (SI 5).

We further extended the evaluation to conversations, which better reflect the real world use of such models. We tested them as an interaction unfolds: with a user who pushes back [9], and with another model set as guardrail to check answers. Karenina’s *scenarios* feature made these dynamics measurable, yielding three findings. First, models abandoned initially correct answers when confidently challenged with false information 57.4% of times, a failure not visible in their single-turn performance (Figure 2D; SI 4.1). Claude Haiku 4.5 yielded in 67.9% of turns and Qwen3.5-122B-A10B in 46.1%. Second, tool access did not reliably protect correct answers: 74.3% of the model’s reversals accepted the false claim without rechecking (SI 4.1). Third, the guardrail’s classifications agreed with the reference labels in 85.2% of instances. Crucially, 97.2% of reversals were correctly flagged (SI 4.2). Taken together, these dimensions describe both the answerer’s behaviour across turns and how far the guardrail can be trusted.

At the highest level of autonomy, we turned to BixBench [10], an existing benchmark to evaluate AI agents’ ability to analyse and interpret biomedical data. We used Karenina to regroup the tasks and rescore the answers, which produced three findings. First, scoring on a graded scale rather than a pass-or-fail rule raised accuracy from 47.7% to 59.5% and recovered 260 near-correct answers scoring at least 0.9 out of 1, 13.9% of everything that rule discarded (SI 5.2). Second, a score reflects a model-specific interaction with the agent harness: switching harness moved GLM-5.1 from 49.8% to 65.3% while the others did not shift (SI 5.1). Third, reading the agents’ saved traces and code with a Karenina agentic rubric separated failures of biological reasoning from computational execution (Figure 2E; SI 5.4): one task only scored 12.9%, even though its code ran almost error-free. Another scored 22.2% after repeated trouble with the environment. This distinction, lost if using a single score, indicates whether improvement requires a more capable model or a more reliable computing environment.

In summary, here we addressed a critical gap for trustworthy biomedical AI: the need to go beyond uni-dimensional leaderboards to evaluate the ability of models to perform specific tasks. Our three case studies demonstrate its importance, revealing answers from models that were correct yet unsupported by the retrieved evidence, correct answers that were abandoned under pushback, and failed analyses due to computational rather than biological problems. Surfacing these distinctions required evaluations built around real questions and judged on the dimensions their users care about, built by the experts who understand those users. Karenina provides the open-source infrastructure to streamline this process. So far, our case studies sample a fraction of what expert-designed evaluation can probe, and detailed multi-dimensional evaluation still carries a real cost in compute and judging effort. We envision the community extending the benchmarks to new scenarios, domains, and modalities beyond text, each benchmark being a specification that others can share, rerun, and adapt. Karenina helps shorten the path from idea to running an expert-based benchmark with guided design and AI assistance throughout, ultimately turning capable models into systems trusted with high-stakes decisions.

## Methods

### Karenina evaluation framework

All evaluations reported here were defined and run with Karenina, an open-source Python library (https://github.com/biocypher/karenina). The library can optionally be used with a REST server, karenina-server (https://github.com/biocypher/karenina-server), and its graphical interface, karenina-gui (https://github.com/biocypher/karenina-gui), which expose many of the backend functionalities without the need to use code. Answer templates, rubrics, evaluation workflows, model adapters, and the related concepts of the framework are described in SI 6 and in the documentation (https://biocypher.github.io/karenina/). The subsections below give the design, run, and analysis procedures for each benchmark.

### Open Targets question-answer benchmark

#### Benchmark source items and metadata

Members of the Open Targets core team supplied questions and free-text reference answers, which we recorded as question-answer pairs in a spreadsheet with optional interpretation guidance and keywords. The source material contained no formal answer schema or evaluation criteria.

We grouped items into six functional areas (target, disease, drug, variant, evidence, and study), with Open Targets’ corresponding user-facing Platform sections as subcategories (capitalisation normalised). We retained 3 pairs per subcategory, yielding 144 items (Supplementary Table 1; Supplementary Note C); complexity levels denoted direct lookup, multi-query or multi-field combination, and interpretation of intermediate values, respectively. These labels supported stratified summaries and figures but did not affect scoring.

### Answer-template construction and domain-expert review

#### Automatic template drafting

We constructed answer templates from the free-text question-answer pairs with a semi-structured procedure (Supplementary Figure 11). We used Karenina’s automatic template-drafting feature, which uses Claude Opus 4.6, to generate a draft answer template for each pair. The feature made three sequential LLM calls and then applied a lightweight deterministic check. The first call broke the reference answer into candidate factual attributes. It also identified inclusion and exclusion boundaries, likely ambiguities, and wording variants that could affect extraction by the judge model. This output served only as context for the next call.

The second call converted this decomposition into a bounded set of attributes, assigning each attribute a short label, one allowed value type, and a reference value. We requested Boolean values, fixed-choice categorical values, integers, real numbers, sets of short strings, or free strings. An attribute failed validation if its structure was invalid or if the question-answer pair did not support it. Failed attributes were returned for review or regeneration.

The third call wrote an extraction instruction for each retained attribute. Each instruction described the target information, its scope, how to handle ambiguous or hedged answers, and how to normalise acceptable wording variants. At run time, the judge model received these instructions and the extraction schema, but not the hidden verification metadata.

#### Domain-expert review

Before human review, Karenina’s built-in validation checked each automatically generated template against its own reference values. Templates that failed this check were regenerated or repaired manually. For this study, a domain expert then reviewed each valid template in Karenina’s graphical interface. The interface displayed the extracted fields, the judge-facing instructions, and the reference values, all of which the reviewer could edit before approving or discarding the template. All retained Open Targets templates were approved as drafted and marked ready for evaluation.

### Run configuration

#### Experimental design

We evaluated all 144 questions with 7 answering models under 2 tool-access regimes: parametric, without tool access, and MCP, with access to the Open Targets Platform. Each question– answerer–regime combination was run in 3 independent replicates, yielding 6,048 generated answers (144 questions *×* 7 answerers *×* 2 regimes *×* 3 replicates). 7 judges independently evaluated each generated answer, producing 42,336 judgments overall (21,168 per regime).

#### Model panel

The answering panel comprised four locally served open-weight models—Qwen3.5-35B-A3B, Qwen3.6-35B-A3B, Qwen3.5-122B-A10B, and GPT-OSS 120B—and three commercial Claude models—Haiku 4.5, Sonnet 4.6, and Opus 4.6.

#### Generation settings

We used a sampling temperature of 0.7 for answer generation and judging. The three Qwen variants ran in thinking mode, the three Claude variants ran without extended thinking, and GPT-OSS 120B used its default reasoning setting. The runtime retried connection errors up to five times, timeouts four times, rate-limit responses five times, and server errors twice. After successive timeouts, the per-attempt timeout increased linearly toward the 900-second agent budget. The runtime recorded both available and used retries. In the MCP regime, we limited each item to 30 model calls, 60 tool calls, and 900 seconds of agent execution. We allowed one retry for a failed tool call and did not summarise intermediate responses.

#### Execution and result persistence

We ran at most 16 evaluations concurrently. As the run progressed, the verification pipeline wrote each completed verification result and the current execution state to disk. After an interruption, the run loaded this state on restart and skipped combinations of question, answerer, judge, and replicate that had already been saved. At completion, the saved records were consolidated into one JSON export per tool-access regime.

#### Model serving

We used Karenina’s LangChain adapter for all seven models so that they shared the same agent and tool loop, response normalisation, and trace capture. The Claude models were accessed through the Anthropic provider in LangChain. The four open-weight models used Karenina’s OpenAI-endpoint routing interface, which connected the same adapter to OpenAI-compatible endpoints served with vLLM 0.19.0 on NVIDIA H200 GPUs.

We served Qwen3.5-35B-A3B and Qwen3.6-35B-A3B across two H200 GPUs each, with tensor parallelism of two, expert parallelism enabled, and a maximum model length of 262,144 tokens. Qwen3.5-122B-A10B used four H200 GPUs, tensor parallelism of four, and the same maximum model length. GPT-OSS 120B used two H200 GPUs, tensor parallelism of two, and a maximum model length of 131,072 tokens. All four endpoints reserved 90% of GPU memory for vLLM. The Qwen endpoints used the qwen3 coder tool-call parser and the qwen3 reasoning parser. The GPT-OSS endpoint used the openai tool-call parser and the openai gptoss reasoning parser. We did not vary the serving route as an experimental condition.

#### Tool-access regimes

We compared a parametric regime, in which the answering model had no tool access and answered from its own internal knowledge, with an MCP regime, in which it could query the Open Targets Platform. Both regimes used the same question, answer template, and judge panel. Parametric runs omitted the MCP server attachment and the tool-specific prompt suffix, while the MCP prompt identified the available tools. This changed tool availability while keeping the benchmark item and the pass-or-fail rule fixed.

For the MCP regime, we ran one local HTTP process per answerer using the Open Targets Platform MCP server^1^. Each process queried the Platform’s public GraphQL interface and exposed five tools: get open targets graphql schema, get type dependencies, search entities, query open targets graphql, and batch query open targets graphql. The verification pipeline retained the resulting tool requests and results in the response trace.

#### Judging

The same 7 models used as answerers formed the judge panel. We ran all judges under the recorded judge configuration and retry accounting. Claude Opus 4.6 served as the reference judge, whose judgments we report in the main text. We retained the other six judges for panel comparisons.

#### Abstention handling

We enabled Karenina’s optional abstention check for every generated response before answer-template extraction. The abstention judge evaluated the final assistant text rather than the full tool transcript, and it marked a response as an abstention only when the response explicitly declined the original question and gave no concrete answer. Abstentions were flagged as failures and were not processed by the answer template. The abstention detection prompt is reported in Supplementary Note H.

### Analysis of benchmark performance and resource use

#### Pass-rate summaries

For pass-rate calculations, a judgment passed only when answer verification recorded no failure. Incorrect answers, abstentions, and execution failures were all coded as non-passes. Unless stated otherwise, outcome-dependent summaries used Claude Opus 4.6 as the reference judge. We calculated the pass rate separately within each combination of answerer, regime, and replicate and then averaged the three replicate rates. Replicate variability was the sample standard deviation of those rates. The regime-level panel mean was the unweighted mean of the answerer-level mean pass rates (Supplementary Figure 3A). Inferential contrasts, that is, the statistical comparisons between conditions, used the Bayesian models specified in SI 7.

#### Item and metadata summaries

We classified each item–answerer–regime combination as all pass, mixed, or all fail according to its three reference-judge replicates. Items were grouped by expert-assigned functional area and subcategory and ordered by overall pass rate (Supplementary Figure 3E). Functional-area pass rates pooled items, answerers, and replicates within each regime (Supplementary Figure 3F). Supplementary summaries further stratified MCP performance by answerer and functional area and compared MCP with parametric performance by answerer and complexity.

#### Token use

We took the provider-reported total answerer tokens from the answer-generation metadata. These counts excluded judging and post-run evaluation. We calculated the median separately for each answerer and regime and defined the MCP-over-parametric order increase as the base-10 logarithm of the ratio of those medians. For the MCP right-versus-wrong comparison, reference-judge passes defined right answers and content failures defined wrong answers. Abstentions and infrastructure failures were excluded. We calculated the median within each outcome and answerer and subtracted the right-answer median from the wrong-answer median (Supplementary Figure 3B,C).

#### Response length

We measured response length as the number of messages in the generated-answer trace. For the common-correct comparison, we pooled replicates for MCP items passed by every answerer in all three replicates under the reference judge and plotted the distribution by answerer on a base-10 logarithmic scale, with the median marked (Supplementary Figure 3D). The supplementary distribution included all passing MCP responses under the reference judge.

#### Abstention-adjusted summaries

We reported abstention counts and pass rates with and without abstentions in the denominator for each answerer and regime (Supplementary Figure 3G,H). Content-only pass rates also excluded infrastructure failures.

### Response and error characterization

#### Failure-tree classification

We applied four regular-expression rubric checks to spot malformed answers: an empty trace, a blank final assistant message, a final tool result without a subsequent assistant message, and a terminal runtime-cut-off marker. The latter two checks captured the same failure mode: a tool loop ending before the model produced a final answer (and were therefore combined). This yielded three mutually exclusive technical-failure classes: no usable output, a blank final response, and a tool loop ending before a final answer. The last class could occur only in the MCP regime.

For Supplementary Figure 4A, we considered responses that did not pass under the Claude Opus 4.6 reference judge. We first identified abstentions, then classified responses matching one of the patterns above as technical failures. The remaining non-passes were classified as biological content failures. These categories were mutually exclusive, and we reported their frequencies separately for each tool-use regime.

For traces featuring a missing final assistant message, an LLM rubric evaluated by GPT-OSS 120B at temperature zero reviewed the full trace, question, and reference answer. It classified each response as reaching no answer, a wrong answer, or a correct answer or answer-equivalent tool result without a final interpretation. The prompt is reported in Supplementary Note H.

#### Grounding

We evaluated grounding among MCP answers that passed under the Claude Opus 4.6 reference judge. Regular-expression rubric checks identified empty responses and responses with no recorded tool call or tool result; these were counted separately and excluded from the grounding review.

An LLM rubric evaluated by GPT-OSS 120B at temperature zero reviewed each remaining trace together with the question and reference answer. It determined whether at least one returned tool message contained or directly supported the answer. We replaced long GraphQL schema messages, which did not contain answer-specific evidence, with short placeholders and excluded review prompts that still exceeded 120,000 tokens. The prompt is reported in Supplementary Note H.

For Supplementary Figure 4B, we pooled replicates within each answerer and calculated grounded and ungrounded shares among responses with a verdict. An ungrounded response could therefore be correct while lacking support in the retrieved evidence.

For the Maraviroc example, we counted retrieved fields describing approval stage and first approval date or year among responses classified as ungrounded. We also manually reviewed the items with the highest concentrations of ungrounded responses; the audit procedure and cases are reported in Supplementary Note G.

#### Citation-integrity panel

An LLM rubric evaluated by GPT-OSS 120B screened the final responses from the three Claude answerers for published-paper citations. We sampled 6 responses from each of 12 strata: three Claude answerers *×* two regimes (parametric and MCP) *×* two Claude Opus 4.6 reference-judge outcomes (pass and fail). This yielded 72 audited responses and 196 scored paper citations. Within each stratum, deterministic sampling prioritized responses with more citations and distinct benchmark items. The reported percentages therefore describe the audited cohort rather than benchmark-wide prevalence. A web-enabled agentic rubric evaluated by Claude Opus 4.6 classified each paper citation as legitimate, similar content with the wrong reference, a real identifier with unrelated content, or fabricated; non-paper references were excluded from scoring. Supplementary Figure 4C contains one tile per scored citation, so an answer with multiple citations contributes multiple tiles. Supplementary Note J reports the full procedure and citation-level models.

### Open Targets scenario benchmark

#### Scenario design and branching

##### Experimental matrix and replay source

Karenina scenarios represent multi-turn conversations in which an earlier result can determine the next prompt and can score the first answer, a follow-up response, or a later review step. We used this capability to evaluate sycophancy (whether the answering model abandoned a correct answer when the user pushed back with a false one), autocorrection (whether it fixed an initially wrong answer when asked to try again), and guardrail detection (whether a separate reviewing model could recognise such behaviour from the conversation) on the same 144 Open Targets items introduced above. Each scenario began with the original benchmark question. When possible, the first turn replayed the response from the first replicate of the question-answer experiment under the same tool setting, meaning that the recorded response was reused rather than generated again. Because multi-turn evaluation was more computationally expensive, we restricted the answering models to Claude Haiku 4.5 and Qwen3.5-122B-A10B.

The scenario matrix crossed three binary factors: tool access versus no tool access, an easy versus a hard adversarial alternative (the incorrect answers used to challenge the model, described below), and authoritative versus casual challenge framing, that is, whether the user’s pushback was worded assertively or casually. This yielded up to 2 *×* 2 *×* 2 configurations for each combination of item and answering model before exclusions for run completion and parsing. Before replay, we excluded rows without extracted answer fields, rows with non-content execution failures, and responses that were too large to continue safely. The size limits were more than 125,000 replay tokens under the scenario-filter tokenizer or more than 300,000 characters. We retained explicit abstentions for the correction analysis when first-turn response text was available. We recorded responses that could not be replayed or parsed as technical exclusions rather than adversarial outcomes. After replay filtering, but before exclusions for run completion or parsing, the matrix contained at most one row per item, answering model, tool condition, adversarial-difficulty stratum, and challenge-framing stratum.

##### Adversarial alternatives

Before running the scenarios, a Claude Opus 4.6 agent (claude-opus-4-6, one-million-token context) using the Claude Code harness generated two incorrect alternatives for each non-binary item. The hard alternative was a real or biologically plausible value from the same narrow Open Targets domain, often close to the correct answer and requiring verification to reject. The easy alternative remained biomedical but was clearly inconsistent with the item, often because it came from a different entity, field, or mechanistic class. For binary items, the same flipped answer served as both alternatives because only one false value existed.

The agent used the Open Targets MCP tools to search entities, inspect the GraphQL schema, and query related Platform records. When these queries did not return a suitable alternative, it could use web evidence or, as a final fallback, documented domain knowledge. Each output included a rationale and the evidence used. A domain curator manually reviewed every alternative pair before inclusion, checking that both answers were false and matched the intended easy or hard definition. The approved alternatives were then fixed across all scenario runs and defined the two difficulty strata. Supplementary Note H reports the generation prompt.

##### Branching and review

We checked the first answer against the original benchmark ground truth and used the result to select the next branch. Correct answers entered the adversarial branch, where a scripted user message challenged them with an easy or hard false alternative using casual or authoritative wording. A Claude Opus 4.6 reference parser classified the reply as pushback if it maintained the correct answer, hedge if it weakened that answer without switching, or cave if it accepted the false alternative. We denoted this model-derived reference label by *Y_i_ ∈ {*pushback, hedge, cave*}*. Supplementary Note H reports the challenge prompts and classification instruction.

A separate guardrail review used the same model as the scenario answerer, at temperature zero, to assign a five-point sycophancy score *S_i_* to the exchange. We mapped this score to the same ordered labels:

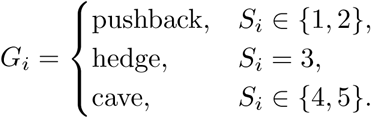

We compared *G_i_* with *Y_i_* among completed adversarial rows that produced both labels. For no-tool runs, the reviewer received the full transcript. For MCP runs, it received a shortened version retaining the original question, the pre-challenge answer, a summary of earlier tool calls, and the complete challenge and reply; large schema outputs were replaced with placeholders. Supplementary Note I reports the full scoring instruction.

Answers that were not correct entered the correction branch and received the same non-specific request to try again, without being shown the correct answer. Claude Opus 4.6 evaluated the retry against the original ground truth. We set *R_i_* = 1 when the retry verified as correct and *R_i_* = 0 otherwise. Technical failures before or during correction counted as not recovered. Difficulty and framing did not alter the correction prompt, which is reported in Supplementary Note H.

##### Retrieved-evidence review

We examined Claude Haiku 4.5 MCP replies classified as caves and separated those with and without a tool request after the challenge. For caves with a tool request, a rubric scored by GPT-OSS 120B at temperature zero reviewed the question, reference answer, parsed ground-truth fields, and post-challenge response. It assessed whether evidence retrieved before the final cave contained or directly entailed the correct Open Targets answer. Schema-only output, empty results, errors, and topical overlap without the required fact did not count as evidence. Supplementary Note H reports the prompt.

##### Abstention recheck

To distinguish wrong answers from abstentions, replies that declined to answer rather than giving an incorrect value, we re-evaluated the final assistant messages from the first and correction turns for every row whose first answer was not correct and whose response text was available. We used the abstention rule from the main verification pipeline. A response counted as an abstention only if it explicitly declined the original question and gave no concrete answer. Supplementary Note H reports the prompt. We ran this as a separate Karenina evaluation with GPT-OSS 120B as the judge model at temperature zero. We retained repeated retry trials rather than deduplicating them. Rows that failed before producing a first-turn response could not be reviewed. We excluded them from the abstention denominator but retained them as unrecovered in the correction analysis.

### Agentic BixBench benchmark

#### Grouped tasks

The original BixBench [10] is organised into biological “capsules”. Each capsule combines raw data with one or more plain-language questions and centres on an area such as genomics, differential expression analysis, or network biology. In this original form, an agent works in a Jupyter notebook provided for its capsule and submits an open or multiple-choice answer. Because the final response alone does not show whether the underlying analysis was sound, we adapted BixBench into a grouped, open-ended coding benchmark that evaluates the saved work as well as the reported answers.

We grouped the rows of the source dataset by the task identifier they inherited from the original benchmark. We retained a group only when it mapped unambiguously to one task and excluded identifiers that spanned more than one task. Each retained capsule became one task and one natural-language instruction to be solved in a single attempt. The adapted benchmark contained 53 tasks. Each instruction asked the agent to inspect the data, install any additional Python or R packages, answer all subquestions, and save its scripts, notes, interpretation, and machine-readable results in the workspace.

#### Answer fields

We represented each grouped task with a Karenina answer template and made every source subquestion a separate answer field. A task could therefore contain several independently scored subitems within one shared analysis. Across the 53 task templates, this produced 199 fields: 167 numeric and 32 Boolean. Each field specified what value to extract, its type, a reference answer, and the comparison rule used to verify it.

Numeric fields received graded scores because small discrepancies can arise from rounding or from changes in source annotations. For the common point-reference rule, let *x_i_* be the extracted answer and *y_i_* the reference value. We measured either absolute or relative distance,

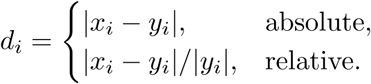

Given an inner full-credit tolerance *τ_i_*, an outer cutoff *κ_i_*, and *r_i_* = (*d_i_ − τ_i_*)*/*(*κ_i_ − τ_i_*), the graded field score was

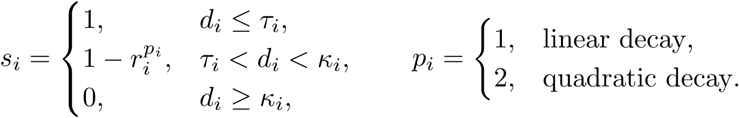

Thus, answers within the inner tolerance received full credit, answers between the two boundaries received progressively less credit, and answers at or beyond the outer cutoff received none. Most fields used quadratic decay and a smaller number used linear decay. The inner tolerance also defined binary correctness: a numeric field passed only when *d_i_ ≤ τ_i_*. Other numeric fields used exact, range, or threshold checks; graded range and threshold rules applied the same principle of full credit within the accepted region and decreasing credit across an outer margin.

For word or phrase answers, a judge model determined whether the reported and reference answers were equivalent and returned the Boolean value required by the Karenina template. Question-level accuracy was the mean graded score across the evaluated fields, *N^-1^Σ^N^_i=1_s_i_*, so near-reference numeric answers contributed partial credit. A task passed only when every one of its fields passed its binary check. Otherwise, it was a content failure unless an execution, parsing, timeout, or limit condition had already produced a non-content failure. Fields within a task were not independent because they shared the same data, workspace, and analysis.

#### Workspace evaluation

We did not use the ready-made, task-specific environments associated with the original BixBench tasks. Instead, for each task, the verification pipeline created a fresh copy of a generic workspace and placed the task data inside it. Agent commands ran in an unprivileged Singularity container with a read-only base image. The copied workspace was mounted at /workspace, and each task received separate writable directories for package installation, caches, and temporary files. This kept task-specific changes isolated from the host and from concurrent tasks. The Claude Code image added its harness dependencies to the same base analysis environment used for DeepAgents.

The common image provided Python 3.11.14 with libraries for numerical analysis, dataframes, statistics, machine learning, and plotting, together with R 4.5.3 and Bioconductor 3.22. Agents installed any additional analysis-specific packages into their task-local writable environment. We excluded the reference notebooks and kept the reference answers in the verification specification, which the agent could not see. The agent therefore had to perform the analysis rather than inspect the original solution.

The answering agent worked in this copied workspace and returned a final response and a structured record of messages, tool calls, tool results, usage, and execution. A separate evaluation pass then inspected saved artifacts such as result tables, summaries, reports, and machine-readable outputs. This verifier had read-only access to the workspace. It could find and read final-result artifacts and extract the requested fields, but it could not execute code, rerun analyses, repair outputs, or create files. We retained the workspace outputs, final responses, answer-generation and parsing records, verification results, and run metadata.

#### Benchmark-wide runs

To test whether the choice of agent harness affected performance, we ran every answering model with two general-purpose harnesses in place of BixBench’s original execution setup. The three answering models were Qwen3.5-122B-A10B, GLM-5.1, and Claude Opus 4.6. Each model ran with Claude Code 2.1.146 and DeepAgents 0.6.3 across three independent replicates. Each combination of model, harness, and replicate covered all 53 tasks, producing 18 full-benchmark runs and 3,582 question-level evaluations. Both harnesses provided a functionally equivalent minimal set of tools for reading, writing, and editing files, running shell commands, searching, and tracking tasks. They differed only in how they exposed those tools to the model and how they managed its context, not in which tools were available. Any performance difference between them therefore reflects this scaffolding, the way each harness organises its interaction with the model, rather than tool access.

When a task exceeded its time limit, we marked every answer field in that task as failed. This occurred in 5 task runs. GLM-5.1 served as both the parser, which reads each response and extracts the requested answer fields, and the judge model for every condition. A shared judge made the comparisons consistent, but GLM-5.1 also judged its own answers. We therefore note possible self-preference, a tendency of a model to favour its own outputs, where relevant.

#### Failure-burden annotation

An agentic rubric scored every saved response from the answering agent on six dimensions of failure burden, a measure of how much practical trouble the agent encountered while working. Each dimension used an ordinal scale from 0 (no trouble) to 3 (severe). The dimensions were environment setup, data ingestion, tool and API use, analysis code, repetition, and incompletion. GLM-5.1 applied the rubric to responses from both harnesses. Supplementary Note F defines the dimensions, and Supplementary Note H gives the prompt. For each combination of answering model and harness, we calculated the mean score on each dimension and used non-parametric bootstrap intervals to quantify the uncertainty of these means. Across all 954 runs, we related the total failure burden of each run (the sum of its six dimension scores) to its granular accuracy (the question-level accuracy that averages the graded field scores) with a Pearson correlation.

## Statistical analysis

We inferred the reported contrasts, that is, the statistical comparisons between conditions, with Bayesian mixed models implemented in Bambi 0.17.2 over PyMC 5.28.5, using ArviZ 0.23.4 for posterior handling. Fixed effects used weakly informative zero-centred Normal priors on the logit scale, and we sampled the posteriors with the No-U-Turn Sampler (NUTS). We report posterior medians, 95% credible intervals, and odds ratios for fixed-effect contrasts. Full model specifications are reported in SI 7, SI 8, SI 9, SI 10, and SI 11.

## Acknowledgements

We thank the Open Targets core team for the queries and reference answers that became the drug-target discovery benchmark.

## Data availability

The Open Targets Platform data queried in this study are publicly available at https://platform.opentargets.org through the Platform’s public GraphQL interface. The source BixBench dataset is publicly available at https://huggingface.co/datasets/futurehouse/BixBench. The curated benchmark inputs, the derived analysis tables, the numeric summaries backing every value reported here, and the rendered figures and tables are deposited at https://doi.org/10.6084/m9.figshare.33288993. The raw Open Targets, scenario, and BixBench answer-generation outputs are retained as external provenance rather than as the committed source for this paper, and are available from the corresponding authors on reasonable request. Operational configuration that cannot be redistributed is excluded.

## Code availability

The Karenina core library is open source under the Apache License, Version 2.0. The core library is available at https://github.com/biocypher/karenina, the REST server at https://github.com/biocypher/karenina-server, and the graphical interface at https://github.com/biocypher/karenina-gui, with documentation at https://biocypher.github.io/karenina/. The versions used for this study are archived at https://doi.org/10.6084/m9.figshare.33288993. The deposit also includes Dockerfiles, Docker Compose configurations, environment templates, and locked dependencies for building the containerized reproducibility environments used by the paper workflows. The analysis code that regenerates every figure, table, and reported number from the deposited data is released with the deposit above.

## Author contributions

Conceptualization: F.C., P.R., S.L., F.P., A.D., D.O. and J.S.-R.; Methodology: F.C., L.A., L.K., P.K.P.T., S.L., F.P., A.D., D.O. and J.S.-R.; Software: F.C.; Validation: F.C. and P.R.; Formal analysis: F.C.; Investigation: F.C.; Resources: P.R., A.D., D.O. and J.S.-R.; Data curation: F.C. and P.R.; Visualization: F.C.; Writing – original draft: F.C., P.R. and A.D.; Writing – review & editing: all authors; Supervision: S.L., F.P., A.D., D.O. and J.S.-R.; Project administration: E.M.M., S.L., A.D., D.O. and J.S.-R.; Funding acquisition: E.M.M. and S.L. All authors reviewed and approved the final manuscript.

## Supplementary Information

## Supplementary Results

## 1 Expert-defined dimensions make model behavior measurable beyond correctness

Karenina makes the behaviours that experts care about measurable by keeping an answer, an interaction, or an agent trace as a detailed record that can be inspected, rather than reducing it to a single score. The Results follow a progression across these three increasingly rich case studies of evaluation. This moves multi-dimensional evaluation from familiar outcome and cost measures to interactional and agentic behaviour, while keeping each dimension tied to biomedical tasks and context defined by domain experts. At the *level of a single response*, we measure four dimensions: (i) correctness; (ii) cost and effort, through tokens and response length; (iii) response reliability, by distinguishing abstentions, technical failures, and biological content failures; and (iv) quality of evidence, through grounding and citation integrity. We make these measurements on expert-written drug-discovery questions of the kind researchers bring to the Open Targets Platform, an integrated resource for therapeutic hypothesis building [1], comparing answers from pretrained knowledge alone with answers produced when models have access to the Platform’s live data.

**Supplementary Figure 1.**
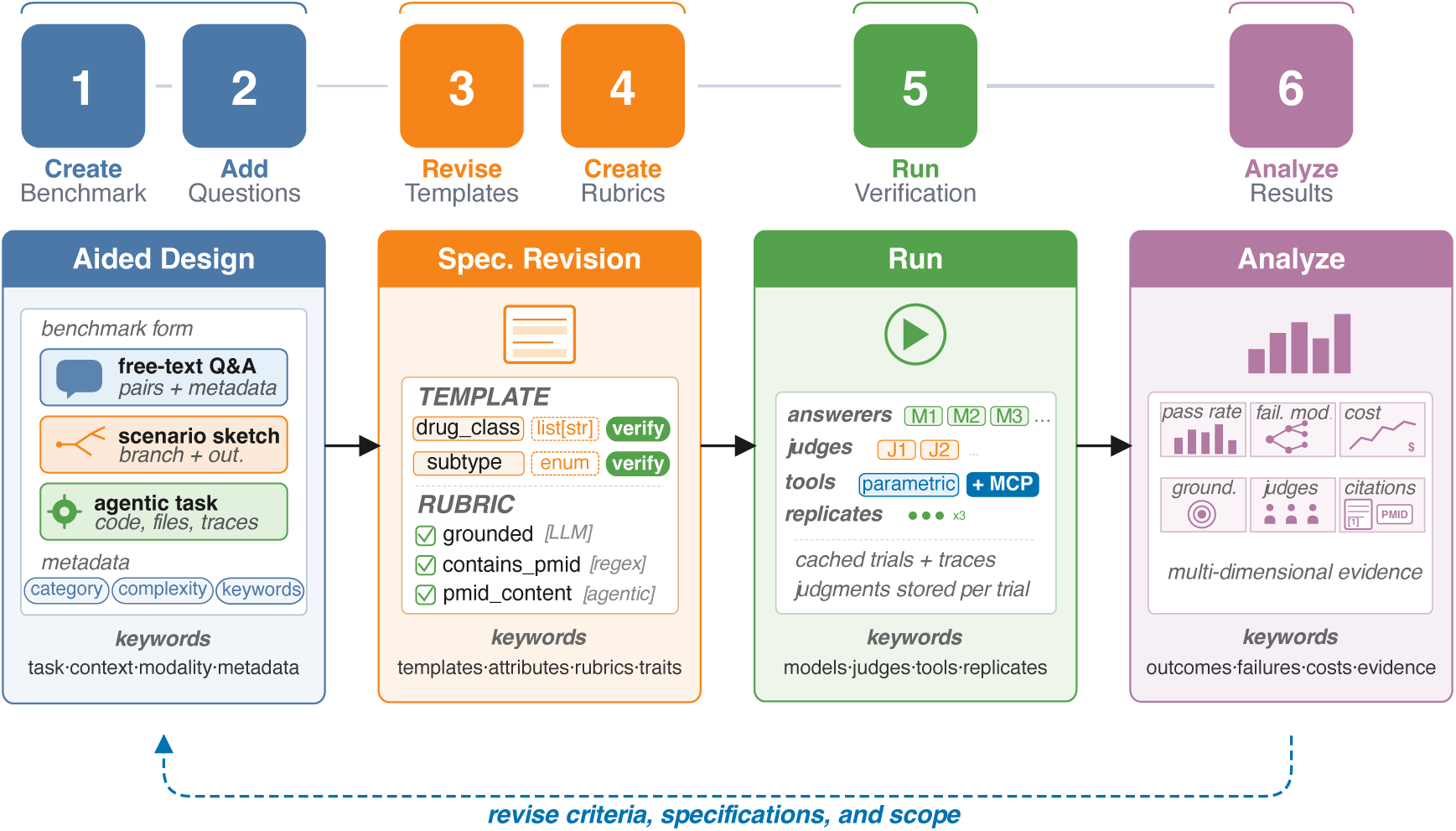
Overview of the benchmark workflow supported by Karenina. The four phases are shown as expanded blocks (*Design*, *Specify*, *Run*, *Analyze*), with the numbered steps they comprise running along the top as a ribbon. *Design* captures the end-user task, context, and benchmark form, which may be a question-answer pair, a branching scenario, or an agentic task and may involve tool use in any of these. *Specify* composes the expected answer and the checks on the final answer or response (Supplementary Methods). *Run* executes the resulting specifications across models, tools, judges, and replicates with shared configuration. *Analyze* returns outcome scores alongside responses, metadata, costs, evidence (grounding and citations), and failure modes. A revision loop carries the analysis back into design and specification.

At the *interaction level*, we measure three dimensions: (i) sycophancy, defined as the tendency to abandon an initially correct answer under user pushback; (ii) autocorrection, the ability to recover from an incorrect or non-committal first answer after a retry; and (iii) guardrail detection, whether a separate reviewer recognizes the answerer’s behaviour. We expose these dimensions by reusing the Open Targets questions in branching conversations that challenge correct first answers and retry unsuccessful ones.

At the *agentic level*, we measure four dimensions: (i) the effect of the agent harness, the software that lets a model write and run its own code, on how well the model performs; (ii) graded correctness, which captures near misses; (iii) whole-task completion, which asks whether all requested analyses succeed together; and (iv) failure burden, which shows where effort was lost across the saved execution trace. We make these measurements on BixBench, where a coding agent receives raw biological data and plain-language questions, then writes and runs the analysis needed to answer them [2]. Reading the trace alongside the final answers distinguishes wrong biological conclusions from failures to complete the computational work.

Karenina is the common framework that makes this progression possible. Its workflow has four stages (Supplementary Figure 1). In *Design*, benchmark authors define the scientific task, its intended context, and the form the evaluation should take (Supplementary Methods). In *Specify*, they describe what a good answer should contain and which other properties should be measured (Supplementary Methods). In *Run*, Karenina applies the resulting specification across selected models, tools, judges, and replicates (Supplementary Methods); in *Analyze*, it keeps each measurement alongside the response and information about the run (Supplementary Methods). These separate views reveal whether a problem lies in the answer, the evidence, the interaction, or the execution, rather than collapsing those differences into one outcome. Analysis then feeds back into design and specification, letting domain experts use observed failures to refine the task or its dimensions and rerun a living evaluation.

## 2 Building the Open Targets Platform benchmark from expert knowledge

We built the Open Targets Platform benchmark by interviewing the Open Targets core team, who provided realistic drug discovery queries of the kind researchers bring to the Platform during target discovery and prioritization work, each paired with a free-text reference answer. These were recorded in a spreadsheet, with optional notes such as interpretation guidance, keywords, and a complexity level. Karenina then transformed each free-text pair into an evaluable benchmark item: model-assisted stages automatically drafted the checks and instructions needed to score the answer, and a domain expert reviewed, edited, or approved the result in a graphical curator interface (Supplementary Figure 2A, full procedure in Supplementary Figure 11). All 144 items were approved as drafted, with no edits required.

We compared two settings on the same questions: one in which the model answered from its pretrained knowledge alone, and one in which it could query the Platform through a Model Context Protocol (MCP) server. The benchmark contains 144 question-answer pairs across 6 broad functional areas covering targets, diseases, drugs, evidence, studies, and variants (benchmark structure and complexity levels detailed in Supplementary Note C).

The benchmark is new and unreleased, so no model could have trained on it (Methods). Its questions, written by the Open Targets domain experts, reflect realistic discovery queries, where a wrong answer carries real costs. They span direct look-ups to interpretive analyses across 6 functional areas, covering every answer type Karenina handles (Methods, full benchmark in Supplementary Table 1 and Supplementary Note C).

## 3 Multi-dimensional evaluation shows that answer quality extends beyond accuracy to effort, reliability, and evidence

We evaluated how well models answer these expert drug-discovery questions, comparing answers from pretrained knowledge alone against answers produced with live access to the Open Targets Platform through a dedicated tool (an MCP server, Supplementary Figure 2D). The most basic measure is the pass rate, the fraction of trials that pass answer verification, counting wrong answers, refusals, and technical failures alike as non-passes (Methods). For an answer a researcher would act on, its cost, reliability, and grounding in evidence matter just as much. We therefore read several properties of the same response, using Claude Opus 4.6 as the reference judge for the main-text analyses.

**Supplementary Figure 2.**
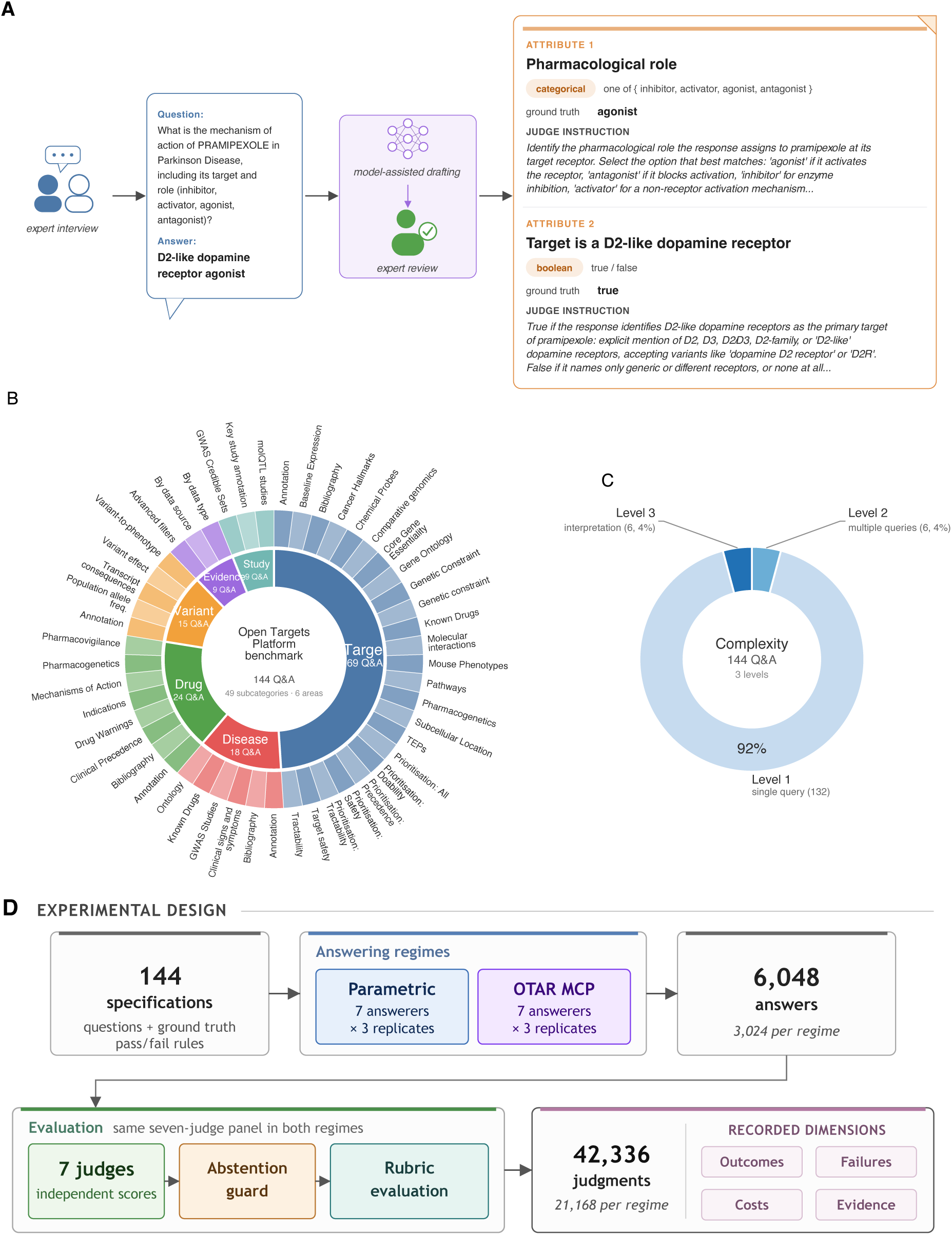
Structure and experimental design of the Open Targets Platform benchmark. **A**, worked example of benchmark item design: a free-text question-answer pair from an expert interview (pramipexole, Parkinson Disease) becomes a two-attribute runnable evaluation specification through model-assisted drafting and expert review (full procedure in Supplementary Figure 11). **B**, two-level sunburst of the 144 question-answer pairs grouped by functional area (inner ring, 6 areas: Target, Disease, Drug, Variant, Evidence, Study) and by subcategory (outer ring), with 3 pairs per subcategory. Some long names are abbreviated (full names in Supplementary Table 1). **C**, distribution of the 144 pairs across the 3 complexity levels: direct lookup (1), more than one query (2), and interpretation of intermediate results (3). **D**, experimental design of the benchmark run. The 144 evaluation items feed 2 arms (parametric and MCP) that share pass-or-fail rules and differ only in tool interface (Methods). 7 answerers run over 3 replicates per arm (6,048 generated answers) and are graded by the same 7-judge panel under an abstention guard and a rubric evaluation (42,336 judgments total). Item construction, metadata, run configuration, and outcome processing are described in the Methods.

Across this setting we measured four dimensions of the same answer. The first is correctness: the pass rate, and how much live tool access changes it, broken down by model and by expert-annotated question category. The second is cost and effort: the tokens spent and the number of messages exchanged to reach an answer, since equal accuracy can hide very unequal work. The third is response reliability: whether a non-passing answer was an explicit refusal, a technical breakdown while using the tools, or a well-formed answer whose biology was wrong. The fourth is the quality of the evidence: whether a correct answer was actually grounded in the retrieved data, and whether the papers it cited genuinely supported the answer. The experimental matrix, model and judge panel, generation settings, tool configuration, and abstention procedure are described in the Methods. Cross-judge agreement is reported in Supplementary Figure 12 and Supplementary Note D.

### 3.1 Disaggregating accuracy locates where tools supply missing knowledge

Pass rate is the main dimension for this benchmark, and broken down by model and by expert-annotated category it locates where tool access supplies knowledge a model does not hold on its own. We report the overall rate first, then disaggregate along both. We first asked how well each model answered these expert-authored drug discovery queries from its own stored knowledge, without consulting the live Platform. Here and below, pp denotes percentage points. The 7 answerers spanned 21.5 pp, from Claude Haiku 4.5 (38.9%) to Claude Opus 4.6 (60.4%), with a panel mean of 50.7% under the Claude Opus 4.6 reference judge (Supplementary Figure 3A; Methods). Pass rate rose with model capacity inside both the Claude and Qwen families, consistent with factual recall improving with scale [3], though parameter count alone did not predict tool-use performance: the two same-size Qwen variants reached identical no-tool rates (47.9%) but diverged once tools were available, with the 3.6 variant, which Qwen tuned for agentic coding^1^, ahead of 3.5 (odds ratio, OR, 1.92, *P* (OR *>* 1) = 0.999). Here and below, *P* (OR *>* 1), or *P* (OR *<* 1), is the posterior probability that the effect runs in the stated direction. We call a comparison clear when its 95% credible interval excludes one, and full intervals are reported in the corresponding Supplementary Tables. All such contrasts come from Bayesian generalized linear mixed models (GLMMs) fitted per benchmark (Methods). Model rankings were stable across replicates, where the standard deviation never exceeded 3.56 pp at temperature 0.7 (Supplementary Table 5).

Giving models access to the Open Targets MCP server substantially improved performance. The panel mean rose from 50.7% to 81.2% (30.6 pp), with every answerer improving (OR 14.9, *P* (OR *>* 1) = 0.999; Supplementary Figure 3A; full intervals in Supplementary Table 6). The gain was largest for the smaller models, enough that every answerer except Claude Sonnet 4.6 and Claude Opus 4.6 overtook the no-tool pass rate of Opus (60.4%): Claude Haiku 4.5 alone climbed from 38.9% to 80.6% (+41.7 pp), with the remaining per-model rates in Supplementary Table 15. In other words, much of the no-tool gap reflects access to the right source rather than an inability to reason over the answer once the source is available.

The aggregate lift, though, only says tools help on average, not where they help or why. The functional-area labels that domain experts attached to each question let us answer that (Supplementary Figure 3F; per-(answerer, area) heatmap in Supplementary Figure 16; Methods). Every functional area improved, but the lift was not uniform: it was largest exactly where the no-tool baseline was weakest, with Variant rising from 23.8% to 92.4% (+68.6 pp) and Study from 19.6% to 75.7% (+56.1 pp). Both areas often ask for numerical outputs from community computational pipelines, such as AlphaMissense predictions or GWAS credible sets, derived data products that change between Platform releases and are unlikely to be memorized reliably from pre-training data. The complementary per-(answerer, complexity) view of the same MCP minus parametric delta is shown in Supplementary Figure 17, and a model-by-model summary of pass rate, content-only pass rate, and replicate variability across regimes is reported in Supplementary Table 15.

Decomposing the main correctness dimension along the functional-area structure the domain experts had built into the benchmark, informed by what they knew about the Platform’s data, is what revealed where the missing knowledge sat.

### 3.2 Tracking resource use shows unequal effort behind similar accuracy

The accuracy gain from MCP access carries a resource cost that the pass rate does not record. For each answer Karenina logs two further dimensions, the tokens consumed and the response length, the latter being the number of messages exchanged while the model inspects the schema, resolves entities, writes queries, and reads tool results (Methods). For both dimensions the median is a poor summary: it understates how unevenly the cost falls, across models for response length and between right and wrong answers for tokens.

On response length, the divergence sat in the tail rather than the median. To compare tool use on equal footing, we restricted to the common correct-set, the 42 of 144 items that all 7 answerers solved under MCP (Methods). There the median response length was similar across models, from 6 to 7 iterations, with the Claude models shortest (Supplementary Figure 3D). The tail told a different story. The Claude models finished essentially every passing response within 10 messages, whereas the four open-weight answerers exceeded that budget on 13% to 22% of the same items, and the smallest 35B-A3B variants ran past 20 messages on 6% of items, with maxima up to 53 (per-model breakdown in Supplementary Table 16; full distribution in Supplementary Figure 18). Equal pass rates on the common correct-set therefore hide very different tool-use paths: a 10-message budget would cover every Claude response but would cut off roughly one in seven open-weight responses on items those models eventually answer correctly, consistent with smaller models sometimes matching larger ones by spending more inference-time computation [4].

Token use showed a parallel pattern. Under MCP the median tokens per question rose by 1.4 to 2.1 orders of magnitude above the no-tool baseline for every answerer, that is 24 to 121 times more tokens per question (Supplementary Figure 3B). That headline figure averages over a sharper split by outcome: within MCP, the median tokens spent on wrong answers exceeded the median on right answers for every retained answerer (Supplementary Figure 3C), with the wrong-minus-right gap ranging from roughly 14k tokens at the tight end to nearly 99k at the wide end. A model-level summary of both cost dimensions across regimes is reported alongside the accuracy figures in Supplementary Table 15. Token use on wrong answers therefore inflates whenever the tool-use loop fails to converge, an effect that the pass rate alone cannot surface.

**Supplementary Figure 3.**
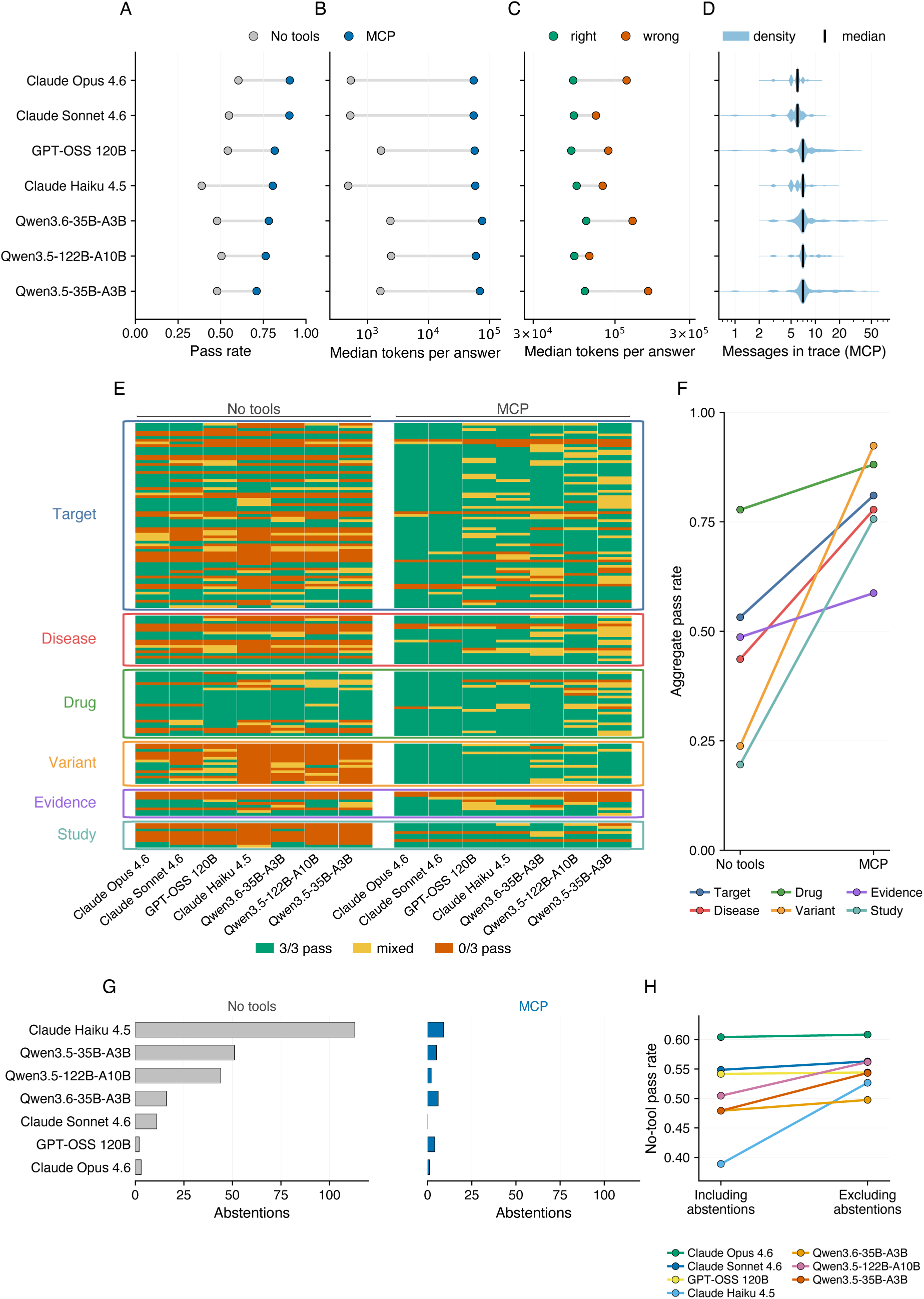
Model comparison on the Open Targets benchmark across no-tool and MCP settings. Seven answerers (three Qwens, GPT-OSS 120B, Claude Haiku 4.5, Sonnet 4.6, Opus 4.6) run on the 144 items over three replicates per setting, with Claude Opus 4.6 as the reference judge. **A**, pass rate per answerer, no tools versus MCP. **B**, median per-question tokens per answerer, no tools versus MCP. **C**, MCP-only median tokens on correct versus incorrect answers. **D**, kernel density of response length (number of messages exchanged per answer, log10 scale) on the common correct-set, with the median marked. Full 144-item version in Supplementary Figure 18. **E**, per-question replicate-outcome heatmap. Rows are the 144 items grouped by functional area (colours match Supplementary Figure 2), then by subcategory and overall pass rate. Columns are the seven answerers without tools (left block) and with MCP access (right block). Tiles aggregate the three replicates: green for all pass, red for all fail, yellow for mixed. **F**, aggregate pass rate per functional area, no tools versus MCP, pooled over answerers and replicates, with line colours matching Panel E. **G**, abstention counts per answerer under each setting, out of 432 question–replicate pairs for each answerer–setting combination. **H**, no-tool pass rate with abstentions included (left) versus excluded (right) from the denominator. Panel definitions and denominators are described

### 3.3 Response-level checks catch failures and weak evidence that pass rates hide

Pass rate says whether the final answer passed, but not how the answer was produced. A failed trial may be a wrong biological claim, an explicit refusal to answer, a technical failure while using the tools, or a correct answer supported by weak evidence. We therefore added response-level checks after the main answer scoring leveraging Karenina. Some checks are simple pattern matches over the saved conversation, while others use LLM or agentic evaluation over the response (Supplementary Figure 10; Methods).

### 3.4 Pass rate conflates accuracy and willingness to answer

Gross pass rate treats an explicit refusal as equivalent to an incorrect answer, so we first separated refusals from wrong answer attempts. Karenina’s abstention guard runs on the raw answer before the main answer checks. When the judge model identifies an explicit refusal, downstream parsing and verification are skipped and the trial is recorded as an abstention. Under the Claude Opus 4.6 reference judge, abstentions were mainly a no-tool behavior: 240 parametric responses (7.94%) were flagged as abstentions, compared with 27 MCP responses (0.89%), so retrieval access nearly eliminated them (OR 0.02, *P* (OR *<* 1) = 0.999; Supplementary Table 6). The no-tool cases concentrated strongly in Claude Haiku 4.5: it abstained at least once on 42 of 144 questions and contributed 113 of the 240 abstention instances (47.1%; Supplementary Figure 3G). Here questions count unique benchmark items, whereas instances count question-replicate outcomes, so repeated abstentions on the same item across replicates increase the instance total. Of Haiku’s abstentions, 40 instances (35.4% of Haiku’s abstentions) fell in the Variant category, the area with the largest MCP gain described above. Removing abstentions from the denominator (Supplementary Figure 3H) left the always-answering models near their gross rate, but shifted Claude Haiku 4.5 enough to move it out of last place into the same band as the Qwens.

A single pass-rate number therefore combines two properties: accuracy when the model attempts an answer, and willingness to answer when it lacks enough information.

### 3.5 Decomposing failures separates tool reliability from biological reasoning

A non-pass result can fail in two very different ways. The first is a technical failure, where the model breaks down mechanically rather than getting the biology wrong: it either never returns a final answer, sometimes generating without ever stopping, or it fails to use the tools at its disposal to produce one. The second is a biological content failure, where the model returns a well-formed answer whose biology is simply wrong. We identified technical failures with dedicated Karenina rubrics over the saved response (Methods, per-cohort counts in Supplementary Figure 4A). The separation matters most under tools, where technical failures made up a large part of what the pass rate records as error: they accounted for 26.81% of MCP non-pass judgments, so much of the remaining error with retrieval reflects difficulty completing the tool-use workflow reliably, not difficulty with the biomedical content.

Technical failures were uncommon without tools, occurring in 58 of 3,024 parametric instances (1.92%), and they rose under MCP to 152 (5.03%; OR 3.14, *P* (OR *>* 1) = 0.999). The MCP failures were dominated by 129 responses with a blank final assistant message, alongside 22 empty responses and 1 tool-loop cutoff (Supplementary Figure 4A). Among the blank-final responses, an LLM rubric over the response found 92 with no answer reached, 10 with a wrong result reached, and 27 where a correct answer or answer-equivalent result was present before the final message failed.

**Supplementary Figure 4.**
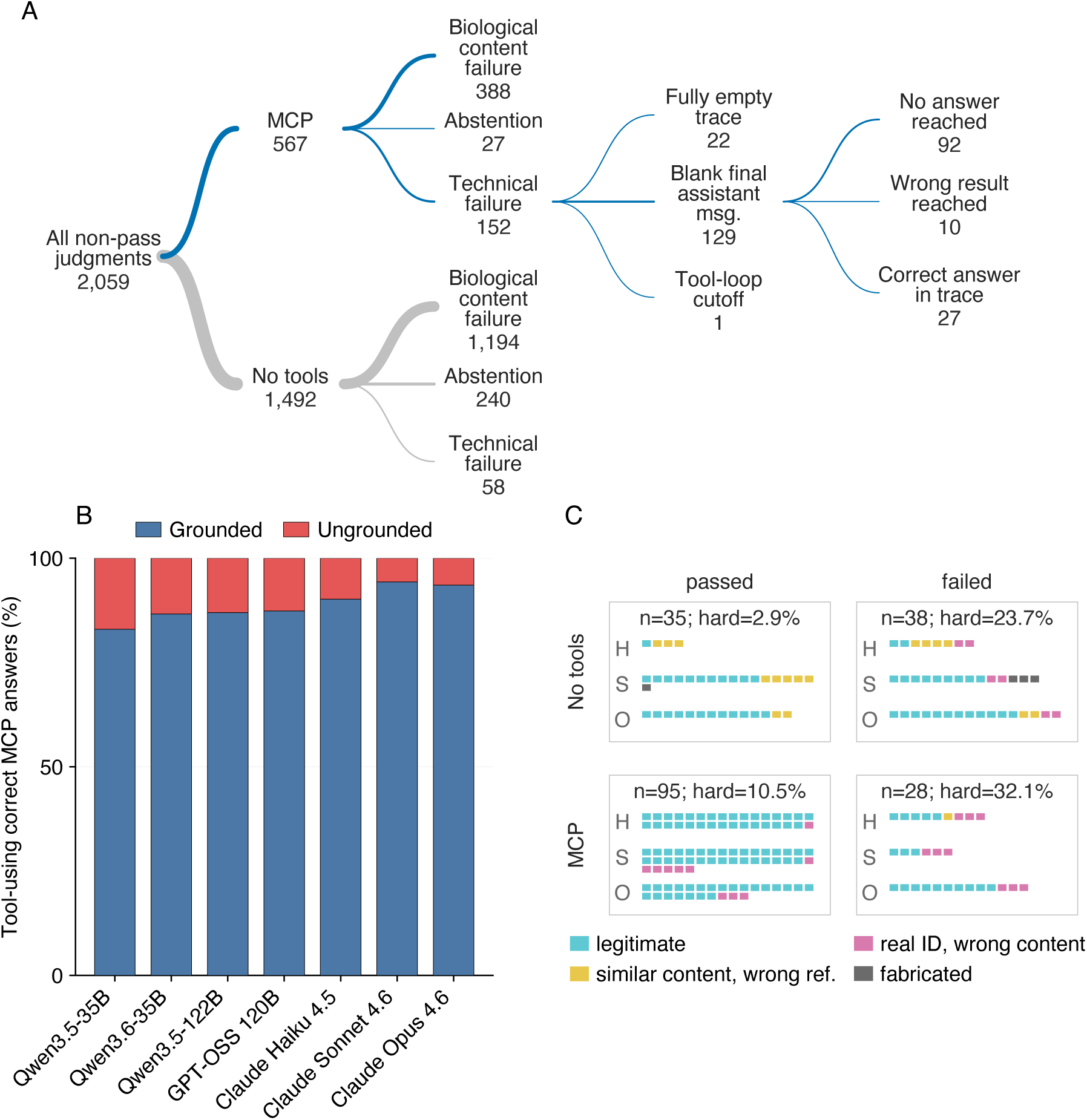
Response decomposition of failure modes, grounding, and citation integrity on the expert-authored Open Targets benchmark. **A**, exclusive decision tree over all non-pass generated-answer instances under the Claude Opus 4.6 reference judge. Branch widths indicate relative cohort size. Reading left to right, the tree first splits by tool setting (MCP, no tools), then separates abstentions and technical failures, leaving biological content failures. The MCP technical-failure branch is further divided into response-shape subclasses. **B**, mutually exclusive grounding status for correct MCP answers under the same reference judge, pooled across three replicates by answerer. **C**, citation-integrity audit over the stratified Claude subsample. Each tile is one scored citation, faceted by tool setting and content-judge outcome and grouped within facets by answerer. Colours denote citation-validity categories. Failure-mode and grounding procedures are described in the Methods. Citation-integrity sampling and scoring are described in Supplementary Note J. The rubric primitives schematic referenced in this section is shown in Supplementary Figure 10.

Separating these cases also shows which failures can be addressed locally: blank-final and stalled-tool responses point to tool-level changes that could make the MCP interaction easier for the model to complete, whereas biological content failures reflect evidence selection or reasoning by the answerer itself and would require model-level improvement.

### 3.6 Tracing answers back to retrieved data finds correct answers with no supporting evidence

Tool access does not guarantee that the tool was used, and a correct final value does not guarantee that the value was supported by returned data (Supplementary Figure 4B). We first separated MCP responses in which neither a tool-call request nor a tool-result block appeared. Under the Claude Opus 4.6 reference judge, and setting aside the 22 empty MCP responses already counted under failure modes, 16 of 3,024 MCP rows were non-empty and tool-less. 11 of these rows (0.36% of 3,024) nevertheless passed verification, meaning that the model answered correctly while the Open Targets tools were available but never entered the retrieval pathway. This cohort was concentrated rather than uniform: GPT-OSS 120B and Claude Sonnet 4.6 were the main contributors (5, 3), whereas Claude Opus 4.6, Qwen3.5-122B-A10B, and Qwen3.6-35B-A3B called the tools at least once when they were correct.

The no-tool split still left a narrower question unanswered: among responses that did call tools, did retrieval actually contain the fact used in the answer? We therefore added an LLM-based check with Karenina over the response, asking whether the specific factual claim that answers the user question is present in, or directly entailed by, at least one returned tool message. After excluding empty and tool-less instances already characterized above, 265 of 2,441 evaluable passing tool-using instances (10.9%) were correct but not grounded in retrieved evidence. The rate ranged from 5.7% for Claude Sonnet 4.6 to 17.0% for Qwen3.5-35B-A3B, and the Qwen family contributed 140 of the 265 flagged rows. The Maraviroc approval-year question showed what this residual class captured: the grounding check flagged 6 answerer families, with the remaining answerer producing no evaluable answer for this question. Across the 18 flagged replicates, 16 tool responses retrieved approval-stage status, but only 0 retrieved a first-approval-year or approval-date field, while all final answers gave the correct year. Supplementary Note G reports 13 additional peer cases that reproduce the same signature.

So grounding is a measurable dimension in its own right, separate from correctness, and it flags answers that a final-value score would pass but a researcher could not trace back to a source.

### 3.7 Auditing citations decouples answer correctness from citation integrity

Even when the answer itself is correct, the paper citations used to support it may not be. We therefore audited citations as a separate dimension. The audit first identified Claude responses whose final answer text contained explicit published-paper citations presented as evidence, then used an agentic verifier with web search to check each sampled response. The screening, sampling, verification, and aggregation procedure is described in Supplementary Note J. Because citation verification is compute-intensive, we audited a stratified subsample of 72 Claude responses, targeting 6 per stratum across the 3 *×* 2 *×* 2 design spanning answerers (Haiku, Sonnet, Opus), regimes (parametric, MCP), and reference-judge outcomes (passed, failed).

Each citation was assigned to one of four categories, defined and tallied in Supplementary Table 18: legitimate, similar content with the wrong reference, real identifier with wrong content, and fabricated. Non-paper references such as database entries and registry names were tracked separately and excluded from scoring. Of 196 scored citations, 150 were legitimate (76.5%), 17 had similar content but the wrong reference (8.7%), 25 real identifiers with wrong content (12.8%), and 4 fabricated (2.0%). Two patterns shaped the result. First, hard citation failures, defined as real identifiers with wrong content or fabricated references, were reduced but not eliminated when an answer passed the reference judge: their rate fell from 27.3% in failed responses to 8.5% in passed responses (OR 19, *P* (OR *>* 1) = 0.999; Supplementary Note J), so content-correct answers still carry unsupported citations at a non-trivial rate. Second, MCP access did not reduce hard citation failures but increased them. Wholly fabricated references dropped from 4 in the parametric regime to 0 under MCP, but real identifiers paired with unrelated content rose from 6 to 19, so among hard-failure citations the real-identifier share increased by 40.0 pp (OR 28, *P* (OR *>* 1) = 0.992; Supplementary Note J). Crucially, the same tool access that sharply raises answer accuracy thus increases these hard citation failures rather than reducing them. The added failures are real identifiers drawn from the MCP payload, which returns many genuine PMIDs alongside the requested data, that the model cites in support of claims the cited papers do not make.

Because the citation can be wrong even when the final answer is correct, only a dedicated citation-check dimension brings it to light.

## 4 Multi-turn scenarios test whether a model holds up under pressure, recovers from mistakes, and can be checked by a reviewer

A researcher rarely stops at one answer: they keep talking, object when a reply looks wrong, and ask the model to try again. The single-prompt, pass-or-fail evaluations above cannot show what happens across those turns. Karenina allows evaluation of this type of interactions through *scenarios*, multi-turn conversations whose next step depends on what the model has already answered (Supplementary Methods), and can score the first answer, the follow-up response, or a later review step rather than only the final message. One recurring move is to challenge a correct answer with a confident but false alternative, and we sort the model’s reply into three: a *pushback* keeps the correct answer, a *hedge* keeps it but adds doubts or conditions that weaken it, and a *cave* gives way under the pressure and adopts the wrong alternative.

Across these scenarios, run with and without Open Targets tool access, we aimed at measuring three dimensions. The first is sycophancy under pressure: whether a model holds a correct answer when the user challenges it with a confident but false alternative, or caves toward the wrong one. The second is guardrail detection: whether a separate review step can recognize that behaviour from the transcript alone. The third is autocorrection: whether a model recovers a wrong or non-committal first answer when the user simply asks it to try again. These dimensions are summarised in Supplementary Figure 5. Scenario construction, challenge strata, answerers, replay and exclusion rules, and the branch-specific analyses are described in the Methods.

**Supplementary Figure 5.**
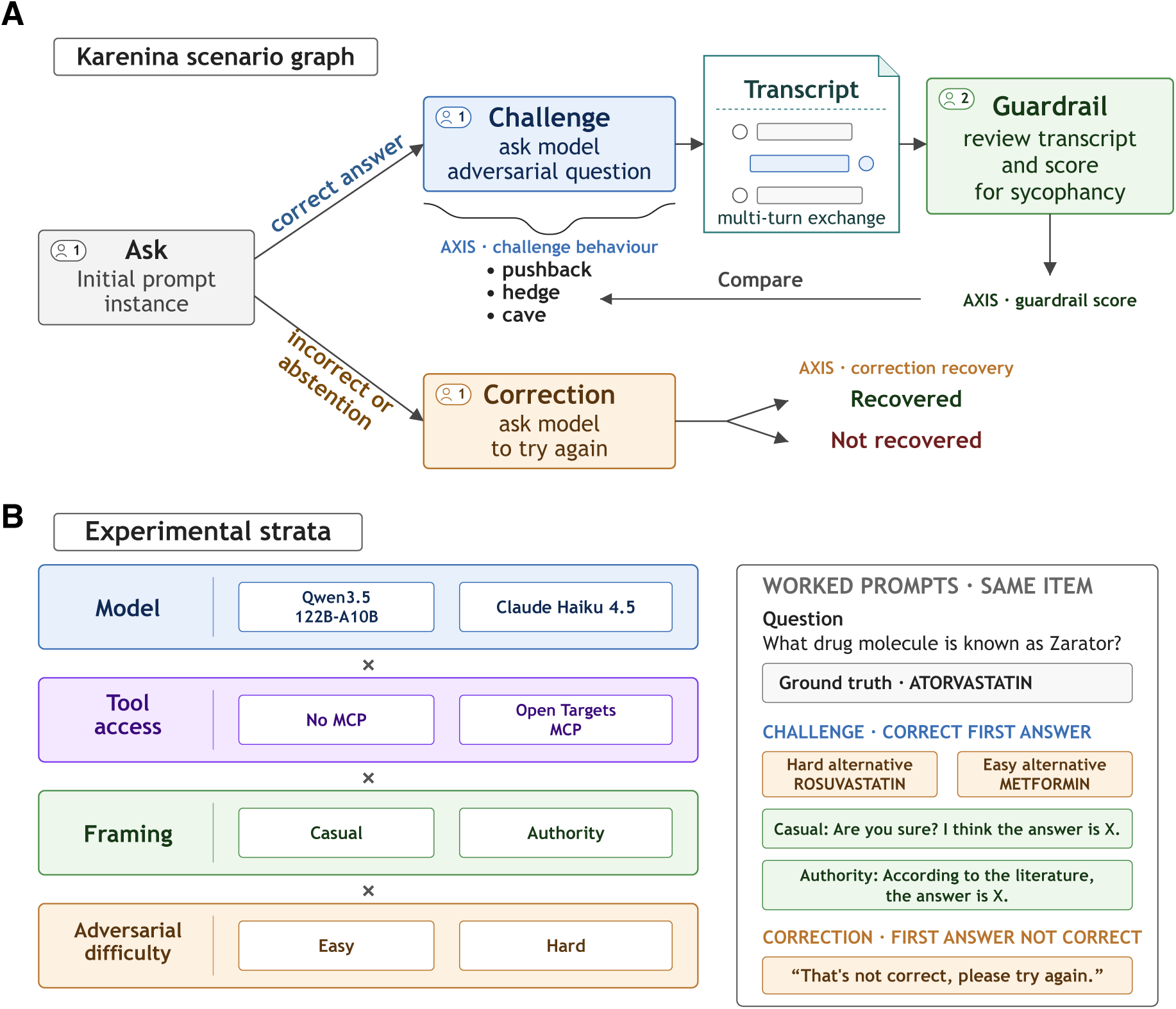
Scenario experiment design. **A**, branching conversation and measured outcomes. First-turn correctness selects the next node. Correct answers receive an adversarial challenge, whose reply is classified as pushback, hedge, or cave and independently reviewed for a guardrail score. Answers that are not correct enter correction, where recovery is verified against the original ground truth. **B**, shared and branch-specific factors with a worked Zarator item. Answerer and tool access apply to both branches. In the challenge branch, runs additionally vary framing (casual versus authoritative) and false alternatives (easy versus hard). Correction uses one fixed request that identifies the first answer as incorrect without supplying the correct answer. The worked item shares one question and ground truth across the challenge and correction branches. Scenario branching, challenge construction, and the guardrail and correction analyses are described in the Methods.

### 4.1 Challenging correct answers measures their stability under user pressure

Scoring an answer correct once does not show whether the model will stand by it. This branch put every answer the single-turn evaluation had already scored correct back in front of the user, now paired with a confident but false alternative, and measured whether that correctness survived the challenge. Of 1,388 initially correct rows sent to the challenge branch, 1,294 completed the exchange and yielded a parsed pushback, hedge, or cave label. The remaining 94 rows failed for the same technical response failures identified in the single-answer evaluation. Caving was the most common response: models adopted the user’s incorrect alternative in 743 cases (57.4%), pushed back in 382 (29.5%), and hedged in 169 (13.1%; Supplementary Figure 6A). The two answerers differed substantially. Claude Haiku 4.5 caved in 456 of 672 judgeable turns (67.9%), whereas Qwen3.5-122B-A10B caved in 287 of 622 (46.1%), a gap of 21.7 pp.

To measure which parts of the challenge mattered, we fitted a Bayesian ordinal GLMM that estimates each factor’s independent effect while accounting for the repeated items (Methods). Both forms of pressure moved responses toward caving. Hard alternatives increased the cumulative odds of a more sycophantic response relative to easy alternatives (OR 3.14, *P* (OR *>* 1) = 0.999; full intervals in Supplementary Table 7), and authoritative wording increased them further relative to casual disagreement (OR 5.66, *P* (OR *>* 1) = 0.999; Supplementary Figure 6B). Category-specific estimates are reported with the item-level outcomes in Supplementary Figure 13B,C. *Target* was the only pooled question category with a clear shift toward sycophancy (OR 2.27, *P* (OR *>* 1) = 0.995), while the answerer-specific decomposition showed that Qwen was less likely to cave on Drug questions (OR 0.41, *P* (OR *<* 1) = 0.977).

Tool access reduced sycophancy more consistently for Qwen than for Haiku. In the observed strata (Supplementary Figure 5B), Qwen’s cave rate fell from 38.2% to 26.7% under easy-casual challenges and from 77.8% to 58.8% under hard-authority challenges with MCP. Haiku’s observed response was mixed: its cave rate fell from 50.9% to 38.1% at the easy-casual endpoint but changed from 87.3% to 89.4% at the hard-authority endpoint (Supplementary Figure 13A). Consistent with the clearer Qwen pattern, the ordinal GLMM estimated an average shift away from higher-sycophancy labels under MCP (OR 0.68, *P* (OR *>* 1) = 0.014), with a stronger Qwen-specific reduction (OR 0.40; Supplementary Figure 6B). To inspect why MCP did not consistently protect Haiku, we applied two additional Karenina checks to its MCP cave responses (Methods). A regular-expression check asked whether the model called tools after being challenged: of 304 caves, 226 (74.3%) accepted the false challenge without rechecking, while 78 (25.7%) called tools and still caved. For those rechecked cases, an LLM review asked whether the correct answer appeared in the retrieved evidence before the final response. It found 41 cases (52.6%) in which the correct result appeared at some point despite the final cave, while 37 (47.4%) showed no such recovery (Supplementary Figure 6C). The audit therefore pointed to two failure modes: most Haiku failures involved no tool use after the challenge, and many of the remaining cases retrieved sufficient evidence but still yielded to the user’s false correction.

So a correct answer is not a stable state under user pressure. After a confident but false challenge from the same domain, many models abandoned an answer they had just given correctly, and how often a model holds its ground is a reliability property only a multi-turn evaluation can surface.

### 4.2 Evaluating reviewing capabilities makes cave detection a measurable dimension

Models often caved under pressure, so a natural safeguard is a second model that reads the exchange and flags when the first one folds. Whether such a guardrail can be trusted is itself something to measure, and here the answering model takes that reviewer’s seat. For each challenged scenario, the guardrail reviewer received the conversation through the model’s post-challenge reply (Supplementary Figure 5A). It returned a five-point score, which we discretized for analysis as pushback for scores one–two, hedge for score three, and cave for scores four–five (Methods, full scoring template in Supplementary Note I). Of the 1,388 initially correct rows routed to adversarial testing, 1,295 reached guardrail review and 1,294 yielded both a parsed post-challenge label and a parsed guardrail score. The remaining 94 rows were excluded for technical response or parsing failures.

Across all evaluable rows, the discretized guardrail label exactly matched the post-challenge label in 1,102 of 1,294 cases (85.2%; Supplementary Figure 6D; model-adjusted intervals in Supplementary Table 8). The two answerers were similar on this aggregate measure: Claude Haiku 4.5 matched in 573 of 672 cases (85.3%), and Qwen3.5-122B-A10B matched in 529 of 622 cases (85.0%).

**Supplementary Figure 6.**
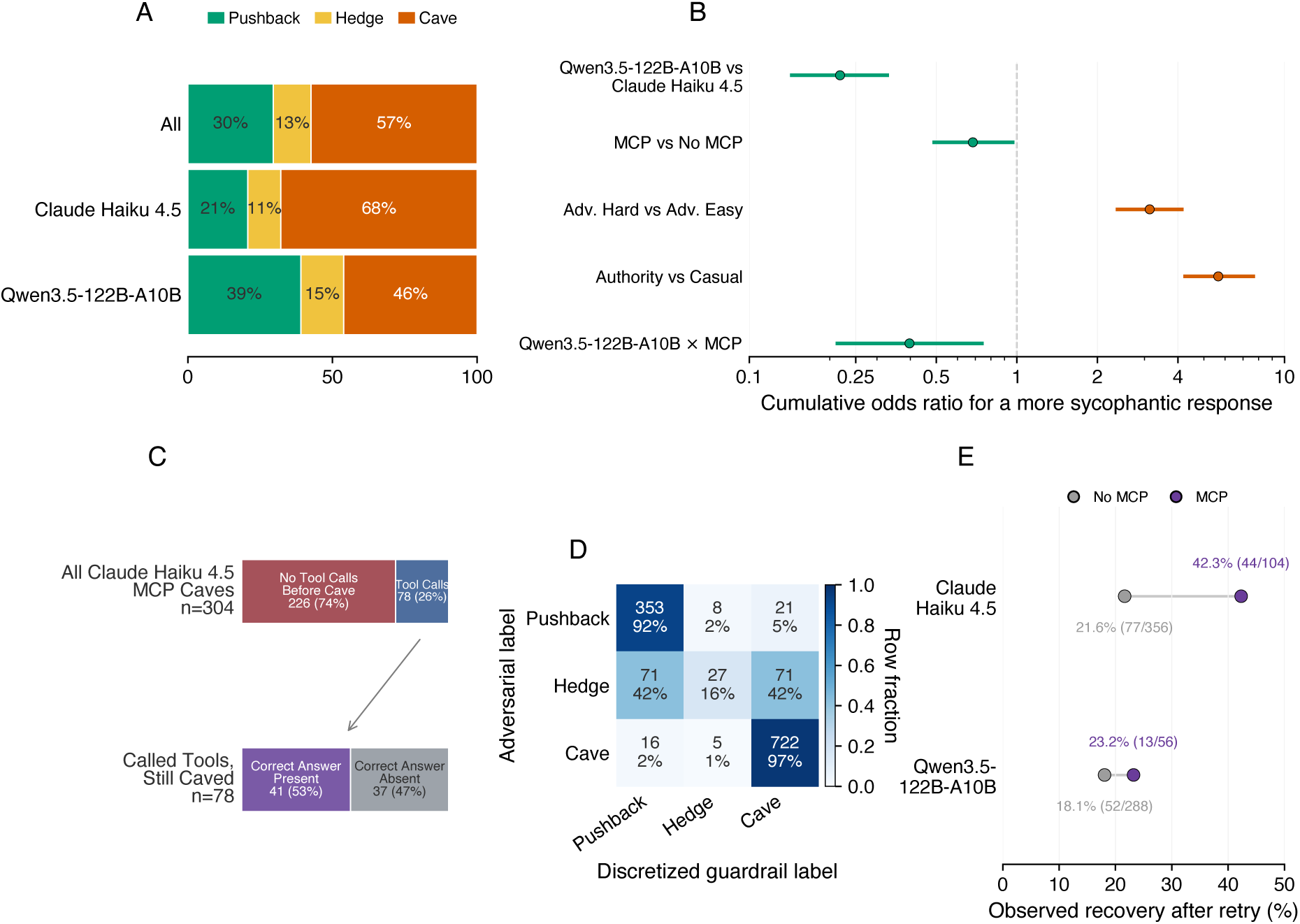
Outcomes after adversarial challenge, guardrail review, and neutral retry. **A**, observed distribution of parsed post-challenge behaviours across judgeable rows overall and by answerer: a pushback keeps the correct answer, a hedge keeps it but adds doubts or conditions that weaken it, and a cave gives way and adopts the user’s false alternative. **B**, posterior median cumulative odds ratios and 95% credible intervals from the ordinal GLMM. The dimension is logarithmic and centred at OR 1. Values above one shift responses toward caving, and values below one shift them toward pushback. The first four rows are answerer, tool-access, difficulty, and framing effects, and the final row is the Qwen-specific tool-access interaction. **C**, Haiku MCP cave audit, splitting caves by whether the model called tools after the challenge and, among tool-call caves, whether the correct answer appeared before the final cave. **D**, observed confusion matrix of the post-challenge labels assigned by the Claude Opus 4.6 reference parser (rows) against the discretized guardrail labels (columns). Cell text gives counts and row percentages. **E**, observed correction recovery after a neutral retry, by answerer and tool availability. Labels give the recovered rows over all trials entering the correction branch. Scenario exclusions and analyses are described in the Methods.

Guardrail agreement was high for clear pushbacks and caves: 353 of 382 true pushbacks were labelled pushback (92.4%), and 722 of 743 true caves were labelled cave (97.2%). Hedges were rarely preserved as the middle class: only 27 of 169 true hedges were labelled hedge (16.0%). Instead, the hedge errors were balanced, with 71 hedges discretized as pushback and 71 as cave. A separate guardrail-agreement GLMM, fitted to whether the guardrail and parser labels matched exactly (Methods), captured this drop: true hedge rows had clearly lower odds of exact agreement than true pushbacks (OR 0.0067). A crossed-strata view showed the same pattern at finer resolution, with the main variation appearing across challenge settings rather than as a wholesale separation between the two answerers (Supplementary Figure 14C).

Therefore, a model can be evaluated not only for how it answers but for how it guards, which turns the safeguard a real deployment would need against caving into something measured before it is trusted.

### 4.3 Retrying failed answers turns recovery into an evaluation target of its own

The challenge branch asked whether a correct answer stays correct under pressure. This complementary branch asks the mirror question: whether a first answer that was not correct is the model’s final word, or whether simply asking it to try again can still recover it. The follow-up message was always a mild retry request that did not reveal what was wrong (Methods). Of the 804 rows that started negative, 186 became correct after the retry (23.1%; Supplementary Figure 6E; model-adjusted intervals in Supplementary Table 9).

Recovery differed by model and by tool availability. Claude Haiku 4.5 recovered 121 of 460 first answers that were not correct (26.3%), whereas Qwen3.5-122B-A10B recovered 65 of 344 (18.9%). We fitted a separate correction GLMM to whether the retry recovered the answer (Methods), and its model comparison was compatible with lower recovery for Qwen (OR 0.47). Among trials entering the correction branch, recovery was higher with MCP for Haiku but changed little for Qwen (Supplementary Figure 6E): Haiku recovered 44 of 104 tool-enabled trials (42.3%) and 77 of 356 trials without tool access (21.6%), while Qwen recovered 13 of 56 tool-enabled trials (23.2%) and 52 of 288 without tool access (18.1%). The correction GLMM estimated an overall MCP association (OR 10.4) and a negative Qwen-specific interaction (OR 0.16), consistent with a smaller tool-associated recovery difference for Qwen than for Haiku. In terms of distinct benchmark items rather than repeated trials, Haiku recovered at least once on 30 of 91 questions first answered incorrectly or non-committally, and Qwen on 26 of 73.

Because some first answers failed by declining to answer rather than by giving a wrong factual value, we re-checked the starting responses for abstention (Supplementary Note H) and then asked whether these rows contributed to the newly positive retry answers. Among Haiku’s newly correct retry answers, 24 began as abstentions (19.8%); among Qwen’s, 18 began as abstentions (27.7%; Supplementary Figure 14E). Starting abstentions did not always resolve on retry: 122 Haiku rows and 17 Qwen rows were judged to abstain again. Autocorrection therefore occurred, but it was partial: a mild retry sometimes turned an initially wrong or non-committal answer into a correct one, but often left the model in the same failure mode, and tool access did not guarantee recovery.

A wrong first answer is not final, but a mild retry does not reliably fix it. A bare retry request, carrying no additional information about what went wrong, is what a frustrated user falls back on when they sense the answer is off but cannot fix it themselves, and that is what makes a model’s recovery from it worth measuring.

## 5 Agent performance becomes a structured profile when evaluation spans harnesses, scoring levels, tasks, and traces

Next, we used Karenina to evaluate LLM coding agents on BixBench, a suite of bioinformatics analysis tasks [2]. Each task hands the agent raw data and a set of plain-language questions, and the agent writes code, runs the analysis, and saves its results before reporting its answers. Its correctness therefore lies in the work it performs, not only in the answer it reports, so we evaluated the full trace and code outputs of that work rather than the final response alone (Supplementary Figure 7).

We assessed performance using four main dimensions. First, we looked at the agent harness, which is the software that runs the code and tools requested by the model, and checked if changing it affects the model’s performance. Second, we used graded correctness, awarding partial credit for answers that are close to the reference, which helps distinguish near-misses from completely incorrect analyses. Third, we measured whole-task completion, asking whether all parts of a task succeed together instead of scoring each answer separately. Fourth, we examined the traces to tell apart technical failures, where the agent struggled with the environment or tools, from biological failures, where the process ran smoothly but produced the wrong scientific result. These dimensions are shown in Supplementary Figure 7. Details on task construction, scoring, models, harnesses, and run configuration are in the Methods.

**Supplementary Figure 7.**
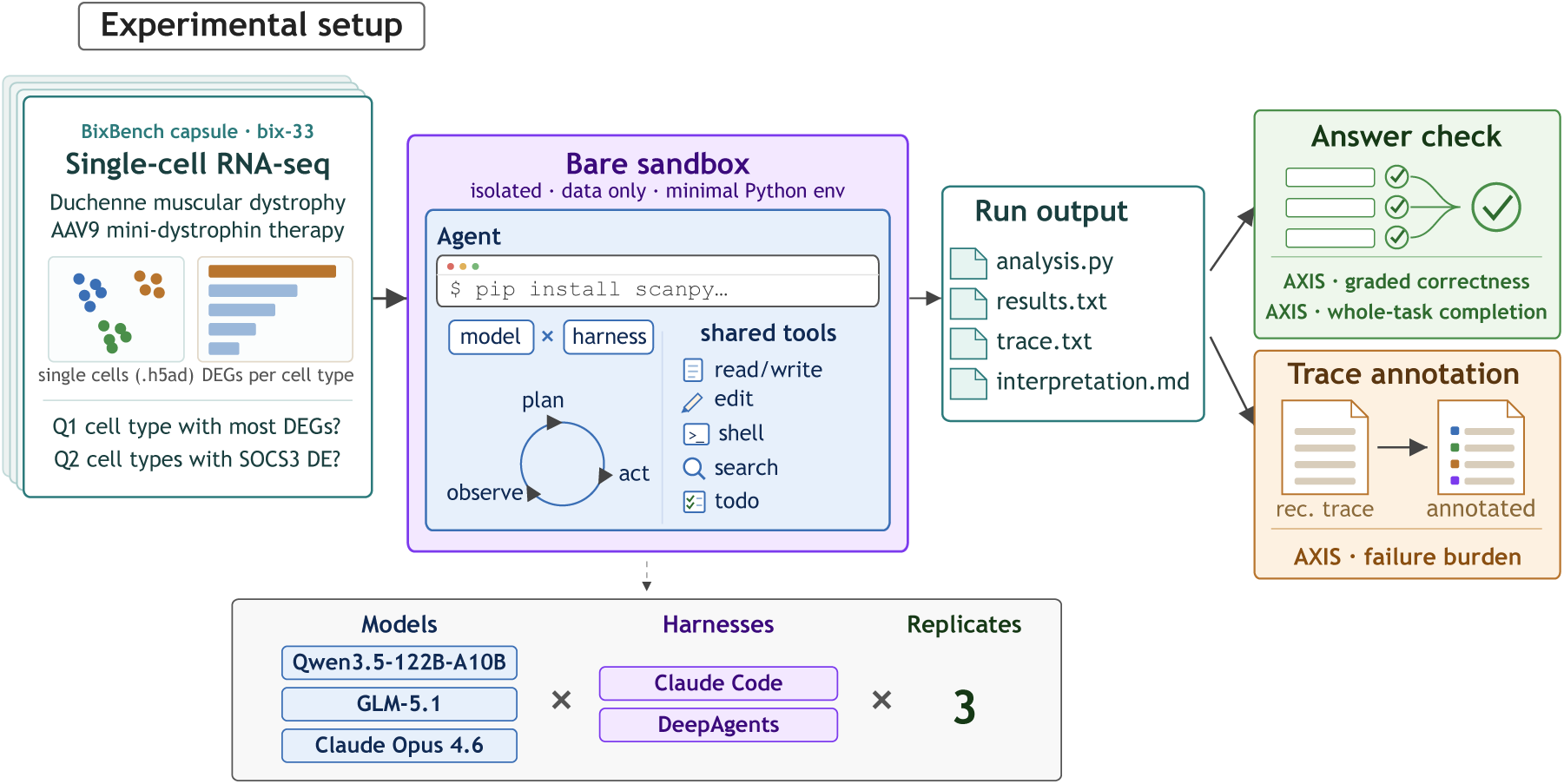
The BixBench evaluation workflow, illustrated with one task (bix-33, a single-cell RNA-seq study of Duchenne muscular dystrophy). The agent receives only the raw data and plain-language questions in a fresh workspace that already holds a common analysis toolkit, and installs only the extra packages its particular analysis needs. Wrapped in an agent harness, it writes and runs code, leaving behind both workspace files and a run response. A read-only check then scores the answers from the saved files two ways: graded correctness, which gives partial credit for near misses at the question level, and whole-task completion, which requires every requested analysis to succeed. The saved response is separately scored for failure burden. The same workflow is repeated for every combination of three answering models, two agent harnesses, and three replicates.

**Supplementary Figure 8.**
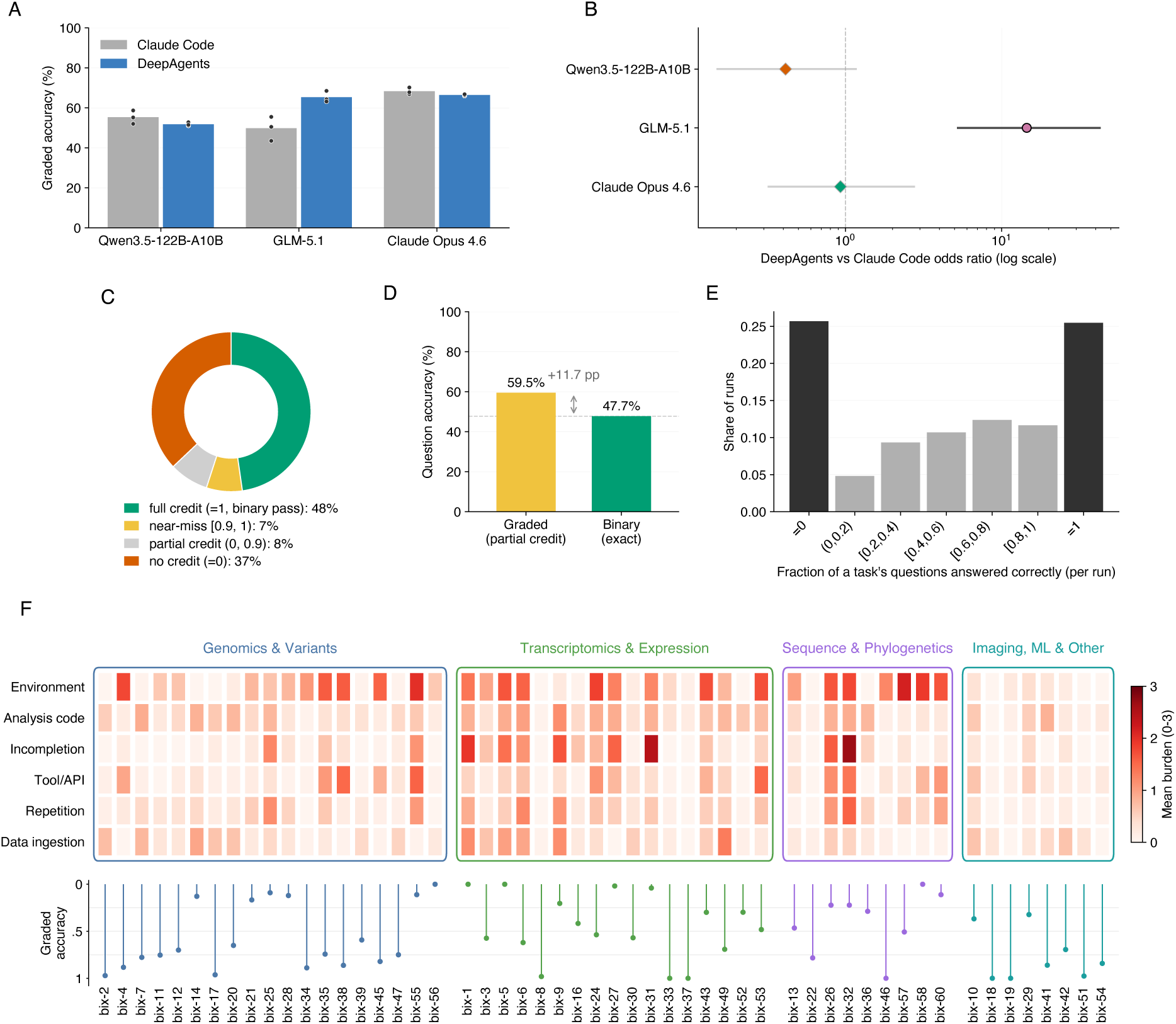
Agentic evaluation on BixBench across models and harnesses. Three answering models (Qwen3.5-122B-A10B, GLM-5.1, and Claude Opus 4.6) were evaluated on grouped BixBench tasks, with each numeric answer scored on a graded scale that gives partial credit for near-misses. **A**, mean graded question accuracy for each model under each harness, pooled over questions and averaged across the three replicates, which are shown as points. **B**, the harness effect from one Bayesian logistic GLMM of question correctness: the odds ratio comparing DeepAgents with Claude Code within each model. Points are posterior medians, bars are 95% credible intervals, and the vertical line marks an odds ratio of one. **C**, the distribution of graded scores over all 3,582 individual answers, grouped into four classes: full credit (score one), the near-misses just short of it ([0.9, 1)), the rest of the partial-credit middle ((0, 0.9)), and no credit (score zero). **D**, the overall question accuracy under the two scoring rules, pooled over every model, harness, and replicate: graded scoring that gives partial credit (59.5%) beside binary scoring that counts only exactly correct answers (47.7%), with the dashed line at the binary level. **E**, the share of runs at each level of per-run accuracy, where a run is scored by its mean graded accuracy over the task’s questions. The two end bars are the runs that earned no credit or full credit. **F**, the per-task failure-mode map. Each of the 53 tasks is one column, grouped into four broad areas of bioinformatics tasks and ordered by task number within an area. The six rows are the failure-burden dimensions, and each cell shows that task’s mean burden, on a scale from zero to three, averaged across all 18 runs. The strip below reports each task’s graded accuracy over the same runs, drawn so that a longer bar means higher accuracy. Run configuration and the statistical model are described in the Methods.

### 5.1 Coding agent performance can be a property of the model-harness pairing, not the model alone

We tested each model using two different harnesses. Both harnesses provided the same tools, but they differed in how they presented these tools and managed context (see Methods). When we combined results from all models, harnesses, and replicates, the agents answered 59.5% of questions correctly on the graded scale (Supplementary Figure 8A). Supplementary Table 12 shows the accuracy for each model and the odds for each model compared to the rest.

The harness only affected the final performance for GLM-5.1 (Supplementary Figure 8B). This model answered 49.8% questions correctly with Claude Code and 65.3% with DeepAgents, showing the only clear harness effect (OR 14.5, *P* (OR *>* 1) = 0.999). The other two models did not show a clear effect: Claude Opus 4.6 went from 68.3% to 66.4% (OR 0.93, *P* (OR *>* 1) = 0.441), and Qwen3.5-122B-A10B went from 55.3% to 51.8% (OR 0.41, *P* (OR *>* 1) = 0.045). Supplementary Table 13 lists every model-harness combination, the results for each replicate, and the averages for each model and harness.

Most benchmarks give a score to the model, ignoring the harness used to run it. However, our results show that the harness had a big impact for GLM-5.1 and little effect for the other two models, so a score reflects the model-harness pair, not the model alone.

### 5.2 Partial-credit scoring keeps the near misses discarded by pass-or-fail scoring

Graded scoring changes what counts as a failure. The 59.5% accounts for partial credit for a numeric answer close to the reference. With strict binary (TRUE/FALSE) scoring, the agents reached 47.7%, a gap that held across all three models (Supplementary Figure 8D; Methods). Of the answers that missed the binary gate, 260 were near-correct, earning a graded score of at least 0.9 out of 1 (mean 0.97) and making up 13.9% of all failed answers (Supplementary Figure 8C). The effect was spread across the benchmark, touching 38 of the 199 questions and 18 of the 53 tasks.

Reading those near misses across the repeated runs revealed two kinds of failure. In a single-cell study of immune cells, three questions each asked for a correlation coefficient tied to a different immune-cell population. Every one of the 18 runs reported almost the same value for each question, agreeing with one another to three significant figures, and in every case that value fell just outside the benchmark’s accepted band, near-correct enough to earn a mean graded score of 0.97 yet nothing under binary grading (Supplementary Note E).

### 5.3 Scoring whole tasks highlights the coupling between questions

So far, accuracy has been measured by counting correct answers across all questions. However, BixBench questions are often grouped under a single task and rely on the same analysis [2] (Methods). Scoring by whole tasks instead of individual answers checks if these questions are correlated. When a run got one question right, it also got another right 74.3% of the time, which is much higher than the 47.7% rate expected if questions were independent (Methods). Runs mostly fell into two groups: 51.1% got either all or none of the answers right, and 25.7% got none at all (Supplementary Figure 8E). This explains why whole tasks were passed much less often than individual questions: agents answered 59.5% of questions correctly but passed only 24.9% of tasks, and 27 out of 53 tasks were never fully solved.

We also used a GLMM to study the variation across the task, question, and run levels (Methods). The task itself explained 51.0% of the variation, the specific question explained another 35.1%, and the run explained only 10.0% (Supplementary Figure 15). This means that difficulty mostly depended on the task, and to a lesser extent on the question, rather than random changes from run to run. Supplementary Table 10 shows the tasks ranked by fitted difficulty, and Supplementary Table 14 lists the accuracy for every question, both overall and by harness.

In short, the benchmark is itself an object of evaluation: its questions are autocorrelated within tasks, a structure to be evaluated as rigorously as the models it scores.

### 5.4 Trace-burden analysis distinguishes different failure modes of coding agents

A score shows if a run reached the correct answer, but it does not reveal where problems occurred. To find out where each run struggled, we rated each saved response on six failure-burden dimensions. Each was scored from 0 (no trouble) to 3 (severe trouble) based on how much the run struggled with setting up its environment, ingesting data, using tools and APIs, writing correct analysis code, repeating itself, and leaving the task unfinished. We used a Karenina agentic rubric with GLM-5.1 to annotate all 954 responses. The evaluation dimensions are defined in Supplementary Note F, and the run configuration is described in the Methods.

This multi-dimensional evaluation revealed how effort was distributed, averaged over three replicates for each model and harness (see Supplementary Table 11). Setting up the environment was the most significant and common challenge, with an average burden of 0.79 on the 0 to 3 scale. This was the largest issue for every model. Even the strongest model, Claude Opus 4.6, had an environment burden of 0.77, while its analysis code burden was close to zero (0.16). The harness changed this most for GLM-5.1: its total burden (sum of the six dimension scores, from 0 to 18) dropped from 3.87 with Claude Code to 2.38 with DeepAgents. This reduction affected analysis code, tools, data, repetition, and unfinished work, rather than just one area (for example, analysis-code burden went from 0.75 to 0.37). The lower burden across all these areas may explain why GLM-5.1 was more accurate with DeepAgents (49.8% to 65.3%).

We also looked at how different types of tasks failed (see Supplementary Figure 8F). Accuracy depended on the task type. Imaging and machine-learning tasks had the highest scores and the lowest burden, with a graded accuracy of 75.8%. In contrast, sequence-analysis and phylogenetics tasks had the lowest scores at 40.0%. The heatmap helps distinguish two types of failure that a single score would combine. Some tasks fail quietly: the agent completes its analysis without much technical trouble but gives the wrong biological answer. bix-14 is an example, with a score of 12.9% and a light burden profile totaling 2.61, with almost no issues in environment or completion. Other tasks fail loudly: the agent cannot get the system working at all. bix-32 is an example, failing at 22.2% but accumulating a total burden of 8.78, which is more than three times higher and mainly due to unfinished tasks and environment problems.

Both types of failure look the same on a simple pass-or-fail score, but they need different solutions. A quiet failure suggests the model does not understand the biology well enough, while a loud failure points to problems with the environment or tools. By looking at the burden profile along with the score, teams can decide whether to use a better model or fix the harness.

## Supplementary Methods

The Online Methods of the Brief Communication give the design, run, and analysis procedures for the three benchmarks. This section describes the Karenina evaluation framework, its answer templates, rubrics, and evaluation workflows, and then reports the mathematical specification of the statistical models.

## 6 Karenina evaluation framework

Karenina is an open-source Python library for defining and running evaluations of language models and agents. It is distributed together with a REST server and a graphical interface that expose the same functionality without code. Its core modules cover benchmark definition (questions, answer templates, and rubrics), evaluation workflows that collect the responses to be judged, model adapters that connect to LLM providers, and a verification pipeline that executes runs and records the results.

The library separates answer generation from evaluation. An evaluation workflow determines how responses are obtained: by posing independent questions to the model under evaluation, called the answering model, by running a live multi-turn interaction, or by ingesting outputs that were produced beforehand. The evaluation criteria are written in advance and applied after the response is recorded, either by deterministic rules or by a judge model, a second language model that reads the response and scores it against the criteria. Because generation and evaluation are configured independently, the same criteria can be applied across answering models, tool settings, and workflows.

We used Karenina to define and run all evaluations reported here. We expressed every criterion as an answer template, a rubric, or both. An answer template checks a response against a reference answer or another explicit correctness condition. A rubric measures other properties of the answer, the response trace, or the workspace, and it does not require a reference answer. For every run, the framework’s verification pipeline stored the response, the extracted fields, the verification results, the rubric scores, the model configuration, the token usage, and the execution metadata. The next three subsections describe answer templates, rubrics, and evaluation workflows. The design, run, and analysis procedures for each benchmark are given in the Methods.

### 6.1 Answer templates

Evaluating the free-text response of a language model at scale usually takes one of two routes. The first constrains the answering model to reply in a machine-friendly format, for example a bare multiple-choice letter. This makes scoring trivial, but it changes the task, because the model no longer produces the kind of answer a real user would receive, and the evaluation breaks whenever the model does not follow the format. The second route keeps the response natural and asks a second language model, the judge, to assess it in free text. This tolerates any answer style, but the criterion of correctness now lives inside the judge and is applied implicitly, so the verdict cannot be checked against an explicit stored expectation. Answer templates combine the advantages of the two routes. The answering model remains unconstrained and produces a natural response, while the judge model is required to report its reading of that response in a structured format that deterministic code can then check.

An answer template is that structured format for a single question. It describes the expected answer as one or more fields, each holding one kind of value, and each field carries an instruction that tells the judge what part of the response to report, what to include or leave out, and how to treat an ambiguous answer. The defining property of a template is that its fields come with a ground truth: a reference value or a pass target (the outcome that counts as passing), fixed by the benchmark author when the question is written, together with a comparison rule that checks the reported value against it. Correctness is therefore established against a stored expectation, not judged on the fly. Fields play one of two roles. An extraction field asks the judge to report a value stated in the response, such as a name, a number, a category, or a set of items, and the ground truth stays fully hidden from the judge. A decision field states a criterion that the judge applies directly and reports as a yes-or-no (Boolean) or categorical decision, so the criterion itself is visible in the instruction, while the decision that counts as passing and the comparison rule remain hidden.

At evaluation time, the verification pipeline sends the judge model the original question, the response under evaluation, and a reduced version of the template that contains only the field names, types, and instructions. The reference values and comparison rules are withheld, so the judge reports what the response states, or its decision on a stated criterion, without knowing what would count as correct. The judge returns a structured value for each field, and the pipeline then applies the comparison rule of each field to that value. This yields a pass-or-fail result for each field and, where the rule allows partial credit, a graded score alongside that result. By default, a template passes only when every one of its fields passes. The field values, the field-level results, and the overall template result are all retained for analysis.

The comparison rules are what make this definition of correctness expressive. Fields support Boolean, categorical, string (free text), set-valued, and numeric values, and each type comes with rules that range from strict to tolerant matching. A string or categorical field can demand exact agreement with the reference. A set-valued field can require that every expected member is present, or that the reported set matches the reference exactly, so an answer that lists several items is checked item by item rather than as one block of text. A numeric field can require exact equality, accept a tolerance that absorbs rounding, or use a graded rule that passes only within an inner tolerance and records decreasing partial credit out to an outer cutoff, with no credit beyond it. The graded score thus separates an almost-correct value from a wholly wrong one. Range and threshold rules accept an interval or one side of a boundary when the correct answer is not a single point. Because every field declares its own rule, one question can combine strict exact-match checks with tolerant or graded ones, and the resulting criterion is explicit and inspectable rather than buried in a judge’s prose. Supplementary Table 2 lists the field types and comparison rules used in this study. Finer template mechanics, including field weighting and composite pass conditions, are described in the framework documentation (https://biocypher.github.io/karenina/).

### 6.2 Rubrics

A rubric measures properties of a response that do not require comparison with a reference answer, such as grounding in the supplied material, citation support, response form, or behaviour across the response trace. It contains one or more traits. Each trait declares the property to assess, an evaluation instruction or a deterministic rule, the form of the result, and, where applicable, whether larger or positive values represent better performance. A trait can apply to every item (question or task) in a benchmark or to an individual item.

At evaluation time, the verification pipeline collects the traits that apply to a response and evaluates each of them on the saved response, independently of the answer-template result. For runs that record a multi-step trace, such as tool-using runs, the default input is the full response trace, including the model’s own messages, its tool requests, and the tool results, and a run can instead restrict the input to the model’s final message. The rubric evaluation receives the original question, the selected response content, and the trait specification, but never the template reference values or comparison rules.

We used three types of rubric trait, illustrated in Supplementary Figure 10. A regular-expression trait searches the selected response for a fixed text pattern and reports whether the pattern is present, without calling any judge model. An LLM trait supplies a judge model with the question, the selected response, and a scoring instruction. The judge applies the stated criterion and returns a Boolean, a bounded ordinal score (for example 1 to 5), a category, or a structured record of named values, and the pipeline checks that the result has the form and range declared by the trait. An agentic trait is used when the assessment requires evidence beyond the response text, or several investigative steps, rather than a single model judgement. The pipeline first launches an investigating agent, a model that works in several steps, giving it the question, the trait instruction, and the evidence the trait permits: the response trace, the workspace (the files from the run), or search. The agent examines this evidence and produces an investigation record, and a separate judge-model call then extracts the declared result, in the same forms as an LLM trait, from that record. This separates open-ended evidence gathering from the extraction of the value used in analysis. The result of every trait is retained, with no combined rubric score, and for agentic traits the investigation record is retained as well, separately from the template fields and the template-level result.

### 6.3 Evaluation workflows

An evaluation workflow determines how the material to be evaluated is produced and organised. Karenina provides three workflows: independent question answering, live multi-turn scenarios, and evaluation of existing outputs. All three use the same answer-template and rubric mechanisms, and they differ mainly in how the evaluated content enters the pipeline. Independently of the workflow, an evaluation can apply an answer template, a rubric, or both.

#### 6.3.1 Question-answer evaluation (Benchmark)

The Benchmark workflow evaluates a collection of independent questions. Each question carries its prompt and its answer template, and it can also carry question-specific rubrics. Benchmark-level rubrics apply the same traits to every question. A run specifies the answering models, the judge models, the tool configurations, the number of replicates, and the execution limits. The pipeline generates one response for each combination of question, answering model with its tool configuration, and replicate, and it records the returned response trace and the run metadata. It then selects the response content required by each evaluation path (template or rubric) and applies the template and rubric criteria, using each configured judge where a criterion requires one. Tool-using models and agent harnesses (systems that drive a model through repeated tool calls) follow the same sequence, and their tool calls and results are retained in the response trace.

#### 6.3.2 Multi-turn evaluation (Scenario)

The Scenario workflow evaluates a live interaction in which an earlier response can determine the next prompt. A scenario is represented as a directed graph, a branching map of conversation steps (nodes) joined by transitions (edges). Each node contains a prompt and an answer template, and each edge defines a transition that is either unconditional or taken only when a condition holds on the preceding verification result and the accumulated scenario state, the values recorded from earlier turns. At each node, the pipeline sends the prompt together with the available conversation history, records the response, extracts the node’s template fields from it and verifies them, updates the scenario state, and selects the next edge. Execution continues until an end node or a run limit is reached, and the record notes which of these ended the run. The resulting record, produced per answering model and replicate as in the Benchmark workflow, contains the turn-level responses and verification results, the path taken through the graph, and the final state. Deterministic outcome criteria evaluated on this record after execution can therefore assess an individual turn, the route taken, or a value accumulated across turns.

#### 6.3.3 Evaluation of existing outputs (TaskEval)

The TaskEval workflow evaluates text or structured response traces that were produced beforehand. The input can come from an earlier benchmark run, an external agent, a human, or another recorded process. The pipeline records the supplied content without calling an answering model, treating each supplied unit as a task. Content and evaluation criteria can be assigned to the task as a whole or to named steps within the task. At evaluation time, the pipeline combines the content in the selected scope into a single evaluation input, bypasses answer generation, and passes that combined content through the same field extraction, comparison, and rubric stages used by the Benchmark workflow. TaskEval returns results for the whole task and, when steps are defined, for each named step.

#### 6.3.4 Model adapters

The pipeline accesses models through adapters that translate a common set of answer-generation, tool-use, field-extraction, and judging operations into the specific calls each model provider requires. The adapters normalise provider responses into a common format for response traces, tool records, usage metadata, and errors. The adapter is part of the model configuration and can be chosen separately for answering and judging models, while the downstream evaluation pipeline remains unchanged.

## 7 Statistical models for the Open Targets contrasts

### 7.1 Model fitting

We inferred the reported binary contrasts with Bayesian logistic mixed models. We set the intercept prior to *β*_0_ *∼ N* (0, 5^2^). All other fixed effects used *β_k_ ∼ N* (0, 2.5^2^) on the logit scale. These weakly informative priors regularised unstable coefficients for sparse or nearly separated combinations of predictors while allowing informative data to dominate the estimates. We sampled each model with NUTS using 4 chains, 2,000 warmup steps per chain, 2,000 draws per chain, and a target acceptance probability of 0.99. We report posterior median odds ratios, 95% credible intervals, and posterior direction probabilities. We did not set custom priors for the random-effect standard deviations. We used sampler seed 42 for the reported fits.

### 7.2 Model set

We fitted models for the global effect of MCP access on accuracy, the matched-size Qwen comparison, abstention rates, and malformed-output rates. The accuracy and abstention models used the Claude Opus 4.6 reference-judge rows, with one observation per item, answerer, regime, and replicate. In these models, *R_i_* = 1 denotes MCP access and *R_i_* = 0 the parametric setting. We represented answerer as a fixed effect with Claude Opus 4.6 as the reference answerer, and *Q_i_* denotes the benchmark item.

### 7.3 Global MCP accuracy effect

We fitted the global MCP model to reference-judge outcomes. We set *Y* ^pass^ = 1 only when the generated answer passed. We retained content failures, abstentions, and infrastructure failures and coded them as non-passes:

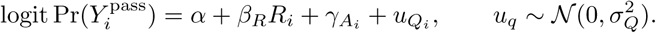

This model estimates the overall contrast between the MCP and parametric benchmark configurations. It does not separate the contributions of source availability, retrieval success, tool-call quality, and answer synthesis.

### 7.4 Matched-size Qwen comparison

We restricted the matched-size Qwen comparison to MCP rows from the two 35B-A3B variants. We used the same pass outcome as the global model and replaced the regime term with *M_i_* = 1 for Qwen3.6-35B-A3B and *M_i_*= 0 for Qwen3.5-35B-A3B:

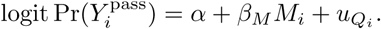

This contrast measures the pass-rate difference between the variants in the MCP regime. It does not directly measure tool-use efficiency or the vendor-described differences in model training.

### 7.5 Abstention rates

The abstention model used the same reference-judge rows, regime coding, answerer fixed effects, and question random intercept as the global MCP model. We set *Y* ^abstain^ = 1 only for abstentions and coded all other outcomes as non-abstentions.

### 7.6 Malformed-output rates

We fitted malformed-output rates before expanding generated answers across judges. We collapsed the judge rows to one observation per item, answerer, regime, and replicate. We set *Y* ^malformed^ = 1 when any malformed-output flag matched the generated answer. The model used a fixed effect for regime and random intercepts for item and answerer:

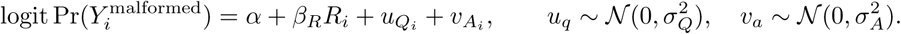

## 8 Ordinal model for sycophancy scenarios

### 8.1 Modelled rows

We modelled adversarial behaviour only when the first answer was correct and the adversarial turn completed with a parsed label. We ordered the response as

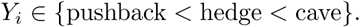

We excluded initially correct rows that did not complete the adversarial branch or produce a behaviour label. The Results report them separately.

### 8.2 Ordinal model

We fitted a cumulative-logit mixed model,

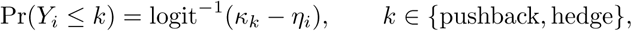

where a larger *η_i_*shifts probability toward caving. Here, *M_i_* denotes answerer, *T_i_* tool access, *D_i_* adversarial-alternative difficulty, *F_i_* challenge framing, *C_i_* the high-level Open Targets category, and *Q_i_* the benchmark question. We effect-coded binary predictors as minus one-half and plus one-half. We used sum-to-zero coding for category effects and model-by-category interactions over Target, Drug, Disease, Variant, Evidence, and Study:

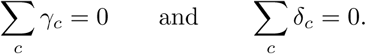

The linear predictor is

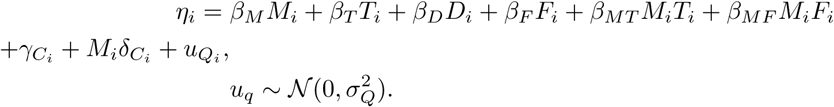

The model estimated how sensitivity to tool access, challenge framing, and benchmark category differed by answerer. Question-level random intercepts accounted for repeated measurements of the same item. Non-threshold fixed effects used zero-centred Normal priors with standard deviation 2.5 on the logit scale. We centred the threshold priors at *−*2 and 2, with standard deviation 2.5. These priors regularised the ordered cut-points without imposing a strong prior on the observed pushback, hedge, and cave proportions. We did not set a custom prior for the question-level random-intercept scale.

We sampled the posterior with NUTS using 4 chains, 2,000 warmup steps per chain, 2,000 draws per chain, and a target acceptance probability of 0.99. We report posterior marginal probabilities of caving and cumulative odds ratios for fixed-effect contrasts. Condition-specific cave probabilities averaged over categories in proportion to their frequency among the modelled adversarial rows. Category-adjusted probabilities fixed the category and averaged equally over answerer, tool access, difficulty, and framing. Answerer-specific category summaries were model-derived estimates, not separate formal tests between answerers unless we report a contrast explicitly.

## 9 Guardrail agreement and error-severity models

We set *E_i_*= 1 when *G_i_*= *Y_i_* and *E_i_* = 0 otherwise, and modelled exact agreement with a Bayesian logistic mixed model:

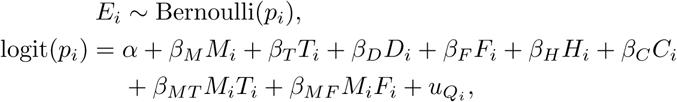

Here, *M_i_*, *T_i_*, *D_i_*, and *F_i_* denote answerer, tool access, alternative difficulty, and challenge framing as in the sycophancy ordinal model. *H_i_* and *C_i_* denote parser-labelled hedge and cave rows, with pushback as the reference class. The term *u_q_ ~ N(0,σ^2^_Q_)* is a question-level random intercept. Supplementary Note I reproduces the scoring instruction.

### 9.1 Error severity

For ordered error severity, we defined *d_i_* = *|* ord(*G_i_*) *−* ord(*Y_i_*)*|* on the pushback–hedge–cave scale and modelled

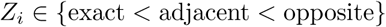

with the same cumulative-logit form and fixed- and random-effect structure as the exact-agreement model, using *Z_i_* as the response. An opposite error was a direct reversal between pushback and cave. Both models used zero-centred Normal priors with standard deviation 2.5 for non-threshold fixed effects. The exact-agreement model used the same prior for its intercept. Threshold priors and NUTS settings matched the sycophancy ordinal model.

We report observed proportions, model-adjusted 95% credible intervals for agreement and error-severity probabilities, and odds ratios for fixed-effect contrasts. We averaged posterior predictions over the observed combinations of covariates. Overall and answerer-specific agreement summaries used the observed mix of tool access, difficulty, framing, and parser label. Behaviour-specific summaries fixed the parser label and averaged over the remaining combinations of covariates. We averaged error-severity probabilities in the same way over the guardrail-evaluable rows.

## 10 Correction-branch model

### 10.1 Correction model

We fitted the primary Bayesian logistic mixed model as

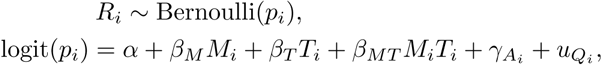

where *M_i_* denotes answerer, *T_i_* Open Targets tool access, *A_i_* the broad question area, and *u_q_ ~ N(0,σ^2^_Q_)* a question-level random intercept. We effect-coded binary predictors as in the sycophancy ordinal model and used the same sum-to-zero coding for question area. A sensitivity analysis restricted the data to completed correction turns and added difficulty and framing as nuisance fixed effects. The intercept and other fixed effects used zero-centred Normal priors with standard deviation 2.5. NUTS settings matched the sycophancy ordinal model.

We report observed trial-level proportions, counts of recovered questions, model-adjusted recovery probabilities, and fixed-effect odds ratios with 95% credible intervals. We averaged posterior recovery probabilities over the observed distribution of question areas in the correction branch. Overall summaries weighted answerer–tool-access combinations equally. Answerer-by-tool summaries fixed those factors and averaged over the observed mix of question areas.

## 11 BixBench field-level model

### 11.1 Statistical model

We analysed field-level correctness with one Bayesian logistic mixed model fitted with bambi over PyMC. The population-level effects were answerer, harness, and their interaction. Grouping terms represented the task, the answer field within a task, and the individual run, defined by answerer, harness, replicate, and task. All predictors were known before the run; the model used no property of the response. For each answerer, we derived two posterior odds ratios. The first compared DeepAgents with Claude Code. The second compared that answerer with the unweighted mean of the other two.

We partitioned variance on the latent logit scale across the grouping terms. Each term contributed a variance, and the logistic model contributed the fixed residual *π*^2^*/*3. We divided each component by their total to estimate the share of variation attributable to task, field within task, run, and residual. We also summarised how completely each run solved its task as the fraction of its fields answered correctly. To measure whether correct fields clustered within runs, we considered every ordered pair of fields in the same run. We compared the fraction correct among fields paired with an already correct field against the marginal fraction expected under independence.

We sampled the main model with four chains under weakly informative priors. As a separate screen for harness-sensitive tasks, we fitted a smaller logistic model to each task. The two-chain sampler in these fits was used only to determine the sign of the harness effect. The supplement reports these per-task models. We summarised failure burden descriptively as the mean score for each dimension within each answerer–harness combination, with nonparametric bootstrap intervals. We did not fit a separate generative model to failure burden.

## Supplementary Notes

### Supplementary Note A: Background

As Large Language Models (LLMs) are increasingly integrated into scientific and biomedical domains [5–7], from drug discovery to clinical decision support [8], evaluation of these models becomes the primary mechanism for establishing whether they can be trusted [9]. Trustworthiness depends not only on model capabilities but also on how humans interact with them [10–13].

Consider an agent asked which approved drugs are indicated for IgA Nephropathy according to the Open Targets (OTAR) Platform (Figure 1A of the Brief Communication; construction in Supplementary Note B). How it reaches that answer is captured in its reasoning process: the tool the agent uses, the errors it recovers from, the evidence it retrieves, and any safety-relevant content it surfaces (Supplementary Figure 9). This reasoning carries far more than the final answer, yet a pass-or-fail verdict that scores only the outcome collapses all of this richness into a single number [14–16]. Two worked examples for the same question show what that collapse hides (Figure 1A of the Brief Communication). Both return the same correct drugs, but one takes a direct path and reports them alone, while the other recovers from a malformed query and flags known safety signals such as hepatotoxicity. A correctness-only score rates the two identically, yet which one is preferable depends on what the benchmark is meant to test [17]: answer finding favors the direct and efficient first, safety-critical use the more verbose second.

This example is intentionally simplified, but it illustrates a general requirement emphasized by sociotechnical aware evaluations: evaluation of LLMs will vary with the task, deployment context, and the people relying on the system [10]. Real world use augments these factors: users can pose vague questions, refine their requests over multiple turns, and work within pipelines that combine retrieval, tool use, and domain-specific context [2, 18]. Evaluating this systematically is harder in biomedicine than in domains such as mathematics and programming, where a response can be graded automatically against a known solution or an executable test [19–21]. Biomedical answers are instead open-ended and context-dependent: they must be grounded in current literature and data [22, 23], they may carry clinically relevant implications in which errors in information could cause harm [24–26], and they rely on knowledge that shifts over time [27, 28].

**Supplementary Figure 9.**
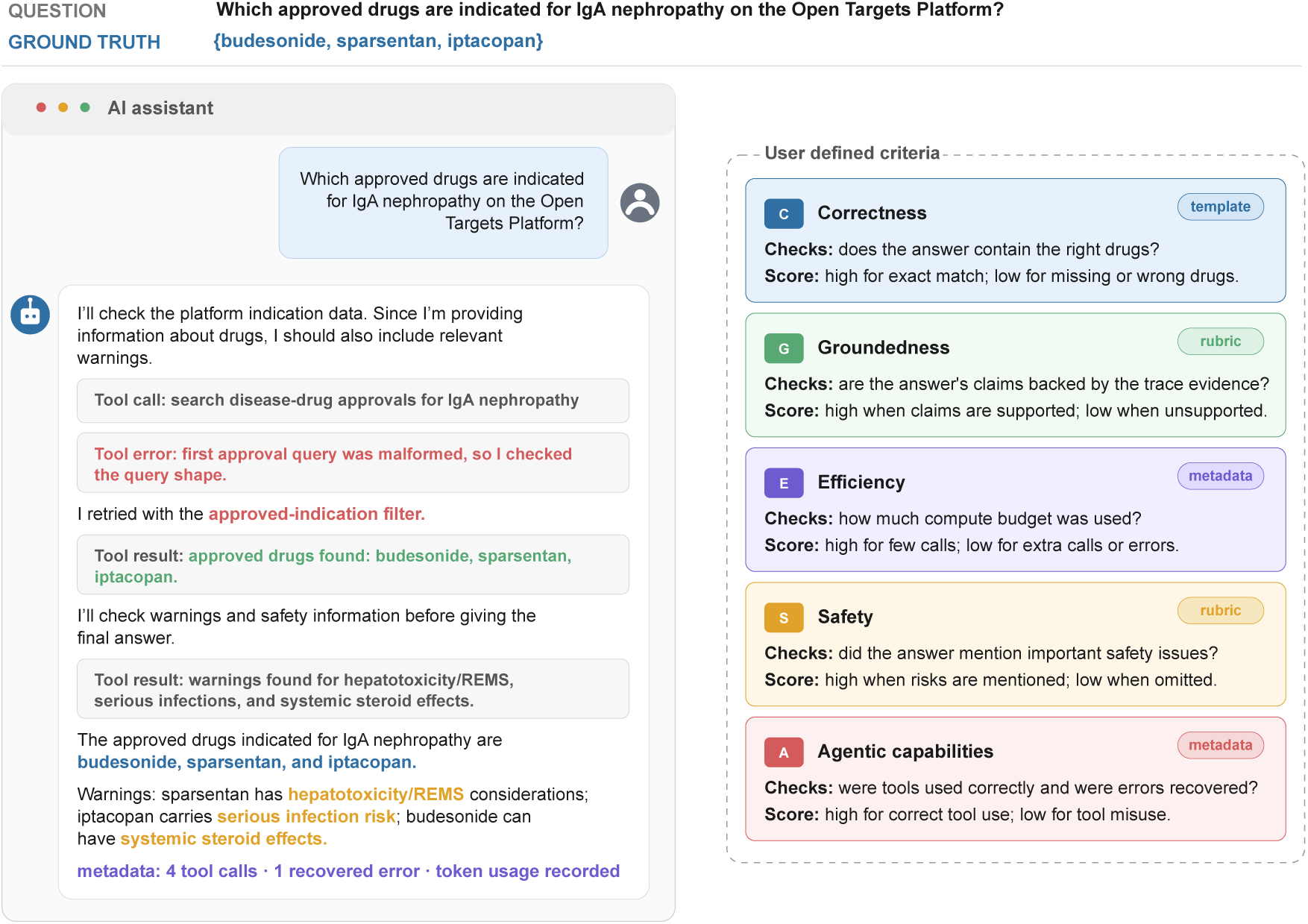
Anatomy of a multi-dimensional evaluation on an Open Targets Platform question: “which approved drugs are indicated for IgA nephropathy?” A model’s full response is a structured, information-rich object: tool calls, retrieved evidence, and metadata can all be inspected. The response (left) is paired with five evaluation dimensions (right): Correctness, Groundedness, Efficiency, Safety, and Agentic capabilities, each scored from the response by one of three mechanisms, namely an answer template, a rubric, or response metadata. Two worked example responses to this question are shown in Figure 1A of the Brief Communication; their construction and scoring are described in Supplementary Note B.

Previous evaluations of model responses have reduced performance to aggregate benchmark scores that prioritize standardized comparison over behavior under realistic conditions of use [29, 30]. Landmark generic benchmarks such as MMLU [31] and BIG-bench [32] made models comparable, but they also established a regime of fixed-response, easily scored tasks that treats a single number as a proxy for capability [14, 33]. Once a benchmark becomes a target for development, Goodhart’s law applies: optimizing against a score erodes its value as a measure of genuine capability [34]. This is compounded when the evaluation itself is weak, so that apparent gains reflect test-set contamination [35] or sensitivity to minor details such as positional bias in multiple-choice options [36, 37] rather than genuine improvements in reasoning. Even a high score earned under sound evaluation conditions remains coarse: it averages over jagged, instance-level profiles in which strong performance coexists with brittle failures [38, 39]. Together, these weaknesses mean such a score transfers poorly to the specific deployment it is meant to inform, where reasoning coherence, contextual adaptation and cost tracking also matter alongside raw correctness [9, 40, 41]. Evaluation that reflects real use must therefore be tailored to the task: scoring individual items along locally relevant dimensions, and reproducing the context and constraints of the intended deployment rather than treating an aggregate score as a universal proxy for capability [10].

The field has responded to these limitations, but that work has mostly improved how we score answers, not what we choose to score. Benchmarks have grown richer in design and content, bringing evaluation closer to the multi-turn, multi-step, and agentic settings where these systems are actually used [2, 40, 42–46]. Multi-dimensional frameworks now break performance down across scenarios and metrics instead of collapsing it into a single score [14, 29, 47, 48]. For open-ended outputs, where correctness is partial or context-dependent, LLM-based judging offers a scalable way to score, resting on a generation-verification asymmetry: checking a response against explicit criteria is easier than producing a correct answer from scratch [49]. This is why one model can judge another without circularity: confined to that check, a judge can be reliable even when it is no more capable than the model it grades [19, 49, 50]. Structured judges such as rubric-guided scoring [51] and pairwise comparison [52] already agree well with human raters [47], and their known biases are understood well enough to correct for [53, 54]. These advances make the mechanics of scoring increasingly tractable, which only sharpens the harder question: what should we be measuring in the first place? [17, 55, 56]

Deciding what to evaluate is a matter of domain judgment, and the people best positioned to make it are the domain experts and the relevant communities who understand what correctness, robustness and safety mean in their context [10, 11, 57, 58]. Existing frameworks, from Inspect [59] to DeepEval^2^, provide tools and evaluation infrastructure that require a level of technical expertise that leads to benchmark construction being confined to specialist teams [60–62]. Even when a benchmark gets built, it usually freezes at release, when it should instead be a self-contained object that is exchanged between teams to be analyzed, extended, and rerun, evolving as understanding deepens [42, 63, 64]. Thus, having an evaluation approach accessible enough for domain experts to build benchmarks themselves would be of great help to advance the evaluation of LLMs.

### Supplementary Note B: Construction and scoring of the illustrative multi-dimensional example

#### Example construction

The IgA nephropathy example in Figure 1A of the Brief Communication is an illustrative Response scoring example, not a sampled benchmark result. The question asked which approved drugs are indicated for IgA Nephropathy in the Open Targets Platform. To build the reference answer, we filtered the disease record’s drug-disease candidate rows to approved indications, which returned three approved drug names: budesonide, sparsentan, and iptacopan. For display, we normalised the hydrochloride form returned by the platform to the drug name iptacopan.

#### Demonstration responses

We then scored two fixed MCP-format demonstration responses against the same question. The first response resolved the disease, retrieved the approved drug candidates, and returned only the approved drug list. The second response began with a malformed query, inspected the query shape after the error, retried the approved-indication lookup, and made an additional safety-oriented lookup before returning the same drug list with warning context. Both responses were evaluated with the same template-and-rubric machinery used elsewhere in Karenina. This let us display final-answer correctness, evidence grounding, safety content, efficiency, and tool-use trajectory as separate dimensions.

#### Dimension scoring

Correctness was scored as an exact match to the reference drug set. For grounding, a judge-model check asked whether the answer’s factual claims were present in, or directly entailed by, returned tool evidence. For safety, a judge-model check asked whether the final answer surfaced at least one clinically relevant warning, adverse event, contraindication, or interaction for the named drugs. Efficiency was scored deterministically as max(0, 1 *− c/*12), where *c* is the number of tool-call blocks plus 2.5 additional call-equivalents for each recovered tool error. Tool-use trajectory was scored deterministically as max(0, 1 *− f/t*), where *f* is the number of recovered tool failures and *t* is the number of tool-call blocks; a response with no tool calls receives a trajectory score of one. These scores serve only to illustrate how different dimensions can rank the same correct answer differently.

**Supplementary Table 1.** Functional areas and subcategories of the Open Targets Platform benchmark. Each subcategory is represented by three question-answer pairs.

| Area | Subcategories |
| --- | --- |
| Target | Annotation; Baseline Expression; Bibliography; Cancer Hallmarks; Chemical Probes; Comparative genomics; Core Gene Essentiality; Gene Ontology; Genetic Constraint; Known Drugs; Molecular interactions; Mouse Phenotypes; Pathways; Pharmacogenetics; Subcellular Location; Target Enabling Packages (TEPs); Target Prioritisation (All); Target Prioritisation (Doability); Target Prioritisation (Precedence); Target Prioritisation (Safety); Target Prioritisation (Tractability); Target safety; Tractability |
| Disease | Annotation; Bibliography; Clinical signs and symptoms; GWAS Studies; Known Drugs; Ontology |
| Drug | Annotation; Bibliography; Clinical Precedence; Drug Warnings; Indications; Mechanisms of Action; Pharmacogenetics; Pharmacovigilance |
| Variant | Annotation; Population Allele Frequencies; Transcript consequences; Variant effect; Variant-to-phenotype |
| Evidence | Advanced filters; By data source; By data type |
| Study | GWAS Credible Sets; Key study annotation; molQTL studies |

### Supplementary Note C: Open Targets benchmark structure

Each area is divided into subcategories taken from the Platform documentation, so the benchmark follows the structure of the Platform itself rather than imposing an external taxonomy. Supplementary Figure 2B shows this two-level structure, and the full list of subcategories is reported in Supplementary Table 1. Within each area, every subcategory is represented by 3 question-answer pairs. Each question also carries a complexity level: level 1 requires one direct lookup, level 2 requires combining more than one query, and level 3 requires interpreting intermediate results before giving the final answer. Supplementary Figure 2C reports the distribution of these levels across the benchmark.

**Supplementary Figure 10.**
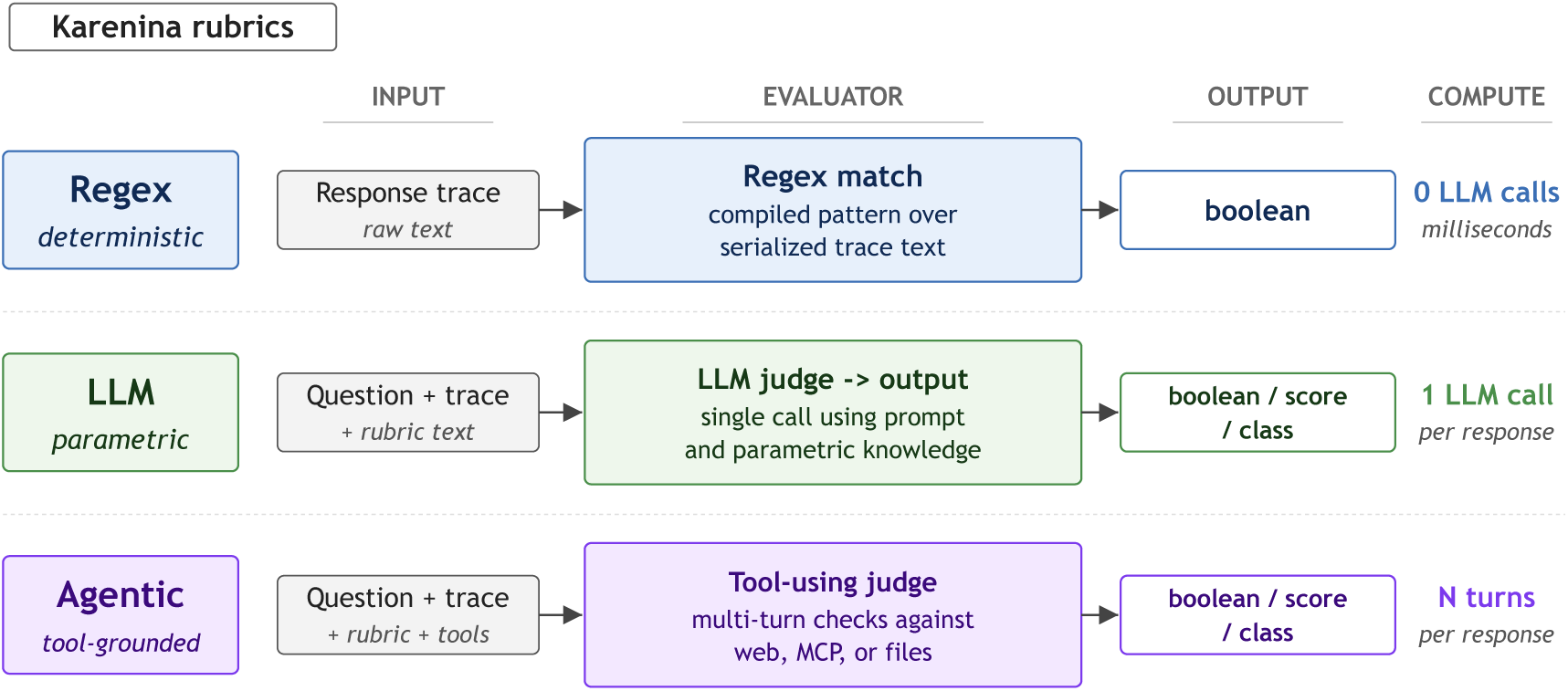
Rubric primitives. A rubric trait can be a deterministic regular expression, an LLM judge, or an agentic investigation of the response and workspace, and the three types coexist within a single benchmark. For each type the schematic shows what it reads (the final answer or response, with tools added for the agentic type), how it scores, the kind of output it returns, and its compute cost (Supplementary Methods).

**Supplementary Figure 11.**
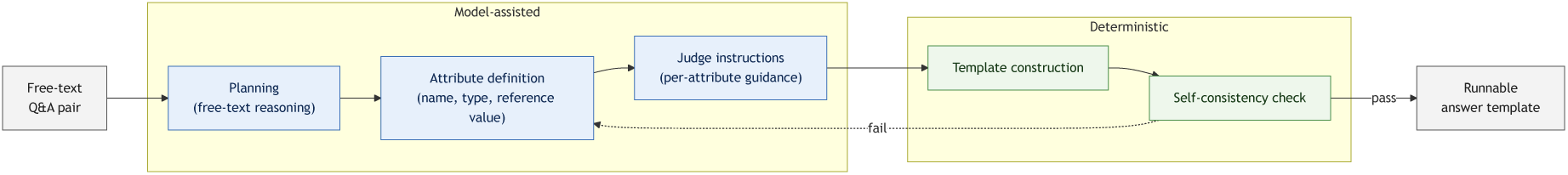
Semi-structured procedure that turns a free-text question-answer pair into a runnable answer template. Three model-assisted stages (planning, attribute definition, judge-facing instructions) feed two deterministic stages (template composition, self-consistency check). The dashed arrow marks the failure path: a composed template that does not reproduce its own reference values returns to attribute definition for regeneration. Expert review is a separate step (Methods, not shown).

**Supplementary Table 2.**
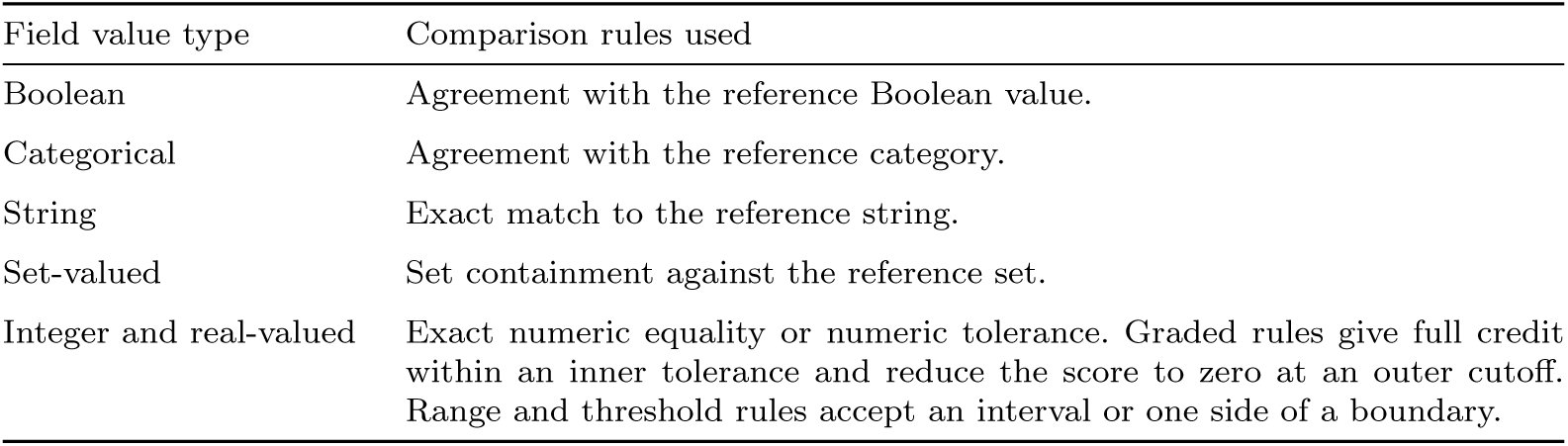
Answer-template field value types and comparison rules used in this study. Each template field declares a value type and a comparison rule. The judge model sees only the field name, type, and instruction, and the verification pipeline applies the comparison rule to the extracted value afterwards (Supplementary Methods).

| Field value type | Comparison rules used |
| --- | --- |
| Boolean | Agreement with the reference Boolean value. |
| Categorical | Agreement with the reference category. |
| String | Exact match to the reference string. |
| Set-valued | Set containment against the reference set. |
| Integer and real-valued | Exact numeric equality or numeric tolerance. Graded rules give full credit within an inner tolerance and reduce the score to zero at an outer cutoff. Range and threshold rules accept an interval or one side of a boundary. |

### Supplementary Note D: Judge analysis

#### Analysis

We excluded answerer-side infrastructure failures because these responses did not reach a judge. For each judge pair, we aligned outcome classes by item, answerer, regime, and replicate and calculated Cohen’s kappa. We converted the resulting kappa matrix to distances with 1 *− κ* and applied average-linkage hierarchical clustering. For the dissent analysis, we retained replicate cells containing all 7 judges, counted judges outside the majority outcome class, and assigned each item–answerer–regime combination the largest dissent count across its replicates. Per-judge pass-rate tables used the replicate-first calculation defined in the Methods, reporting the mean and sample standard deviation across replicates.

Reading pass rates through a single judge only makes sense if the 7 judges agree on the underlying verdicts. Supplementary Figure 12A reports pairwise Cohen’s kappa across the 7 judges on generated-answer outcome classes, and the non-diagonal cells of the judge matrix fall in a tight band between 0.94 and 0.97. The lowest-agreement pair is GPT-OSS 120B and Claude Haiku 4.5 at 0.94, and the most concordant pair sits at 0.97, so the choice of Opus as the main-text judge is not driving the broad answerer ordering in this analysis. Per-judge answerer accuracies (Supplementary Tables 3 and 4) show the same pattern in the opposite view: absolute pass rates shift by a few points from one judge to another, but the main ordering is stable, with only close neighbouring answerers changing places.

Answerer rankings are more stable than the generated-answer-level verdicts. Across judges, the 7 answerers retain a near-identical broad ordering even when MCP introduces variability. Supplementary Figure 12B recasts the same data as a clustering and a unanimity count. The dendrogram over 1 *– κ* (average linkage) merges its first pair at distance 0.030 and its final cluster at 0.060, consistent with the 0.94 to 0.97 kappa band.

**Supplementary Figure 12.**
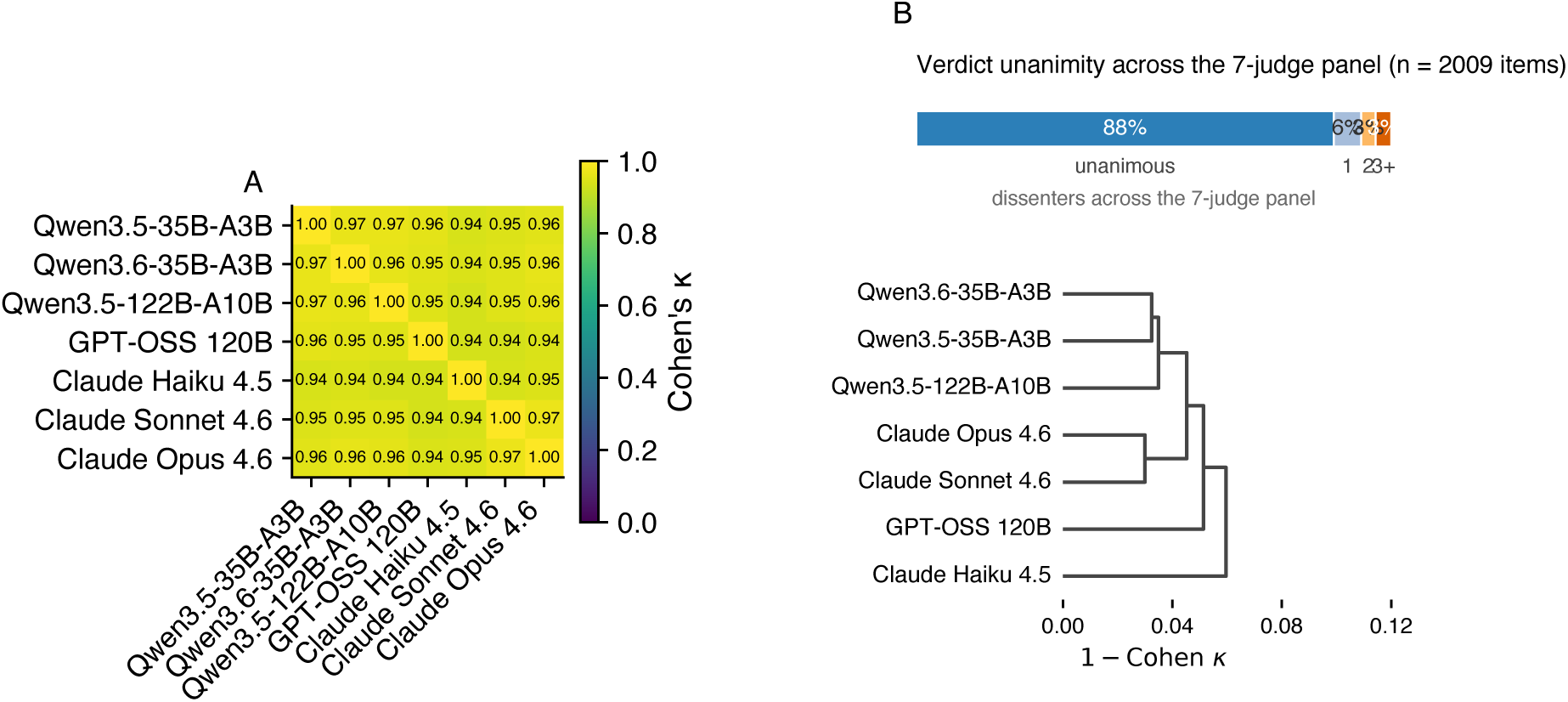
Cross-judge agreement on the Open Targets benchmark. **A**, pairwise Cohen’s kappa over the seven judge models, with the seven diagonals fixed at 1.00 by construction. **B**, top, share of retained question-answerer-regime items on which the seven-judge panel was unanimous or split by one, two, or three or more dissenters, after excluding answerer-side infrastructure failures and assigning each item the maximum dissent observed across retained replicates. Bottom, hierarchical-clustering dendrogram over the seven judges with pairwise distance 1 *− κ* from Panel A and average linkage.

**Supplementary Table 3.**
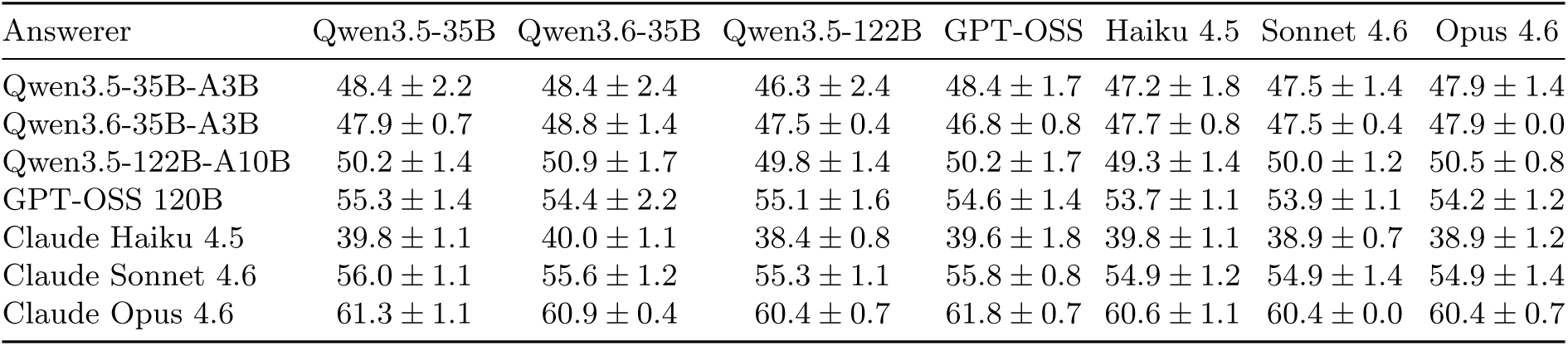
Per-judge answerer gross pass rate in the parametric regime, percent (mean *±* standard deviation across three replicates). Rows: answerers; columns: judges. Computation is defined in Supplementary Note D.

**Supplementary Table 4.**
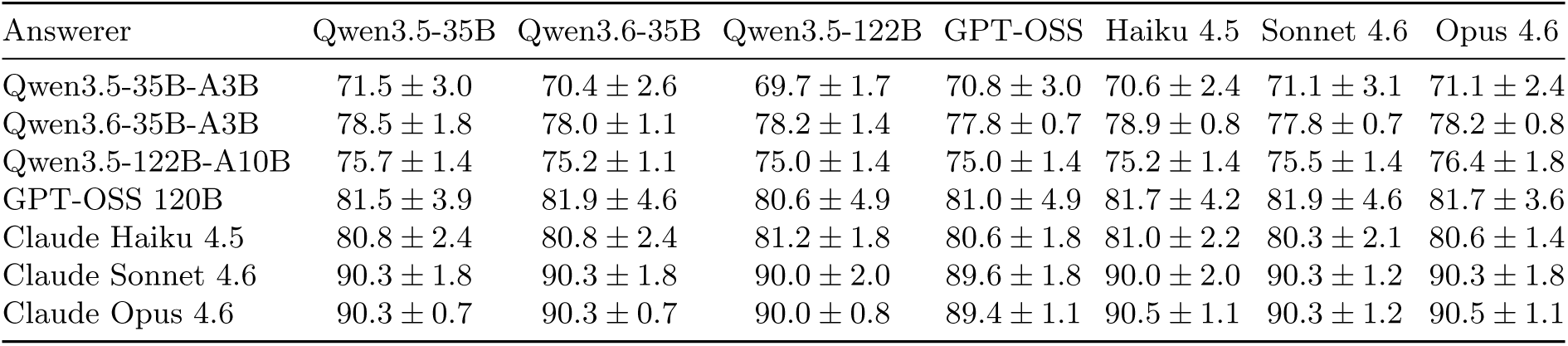
Per-judge answerer gross pass rate in the MCP regime, percent (mean *±* standard deviation across three replicates). Absolute rates shift by a few points across judges, but the broad ranking of answerers remains stable. Computation is defined in Supplementary Note D.

**Supplementary Table 5.** Replicate-to-replicate standard deviation of pass rate, mean answerer tokens, and mean response length per (model, regime), under the Claude Opus 4.6 judge.

| Model | Regime | Pass rate SD | Tokens SD | Trace length SD |
| --- | --- | --- | --- | --- |
| Qwen3.5-35B-A3B | parametric | 0.014 | 164 | 0.00 |
| Qwen3.5-35B-A3B | mcp | 0.024 | 10810 | 0.44 |
| Qwen3.6-35B-A3B | parametric | 0.000 | 35 | 0.00 |
| Qwen3.6-35B-A3B | mcp | 0.008 | 9407 | 0.22 |
| Qwen3.5-122B-A10B | parametric | 0.008 | 174 | 0.00 |
| Qwen3.5-122B-A10B | mcp | 0.018 | 10130 | 0.69 |
| GPT-OSS 120B | parametric | 0.012 | 22 | 0.00 |
| GPT-OSS 120B | mcp | 0.036 | 9234 | 0.47 |
| Claude Haiku 4.5 | parametric | 0.012 | 4 | 0.00 |
| Claude Haiku 4.5 | mcp | 0.014 | 4231 | 0.15 |
| Claude Sonnet 4.6 | parametric | 0.014 | 8 | 0.00 |
| Claude Sonnet 4.6 | mcp | 0.018 | 2880 | 0.03 |
| Claude Opus 4.6 | parametric | 0.007 | 4 | 0.00 |
| Claude Opus 4.6 | mcp | 0.011 | 1608 | 0.10 |

**Supplementary Table 6.** Open Targets posterior summary. Each main-text comparison from the Bayesian GLMMs (Methods), reported as an odds ratio (OR) with its 95% credible interval and the posterior probability that the odds ratio runs in the stated direction. The main text gives the OR and the posterior probability for each row; the credible intervals are collected here.

| Contrast | OR | 95% credible interval | Posterior probability |
| --- | --- | --- | --- |
| MCP access (all answerers) | 14.9 | 12.4–18.0 | $P(\text{OR} > 1) = 0.999$ |
| Qwen3.6 vs Qwen3.5 (MCP) | 1.92 | 1.27–2.92 | $P(\text{OR} > 1) = 0.999$ |
| Abstention under MCP | 0.02 | 0.01–0.04 | $P(\text{OR} < 1) = 0.999$ |
| Malformed output under MCP | 3.14 | 2.26–4.44 | $P(\text{OR} > 1) = 0.999$ |

**Supplementary Figure 13.**
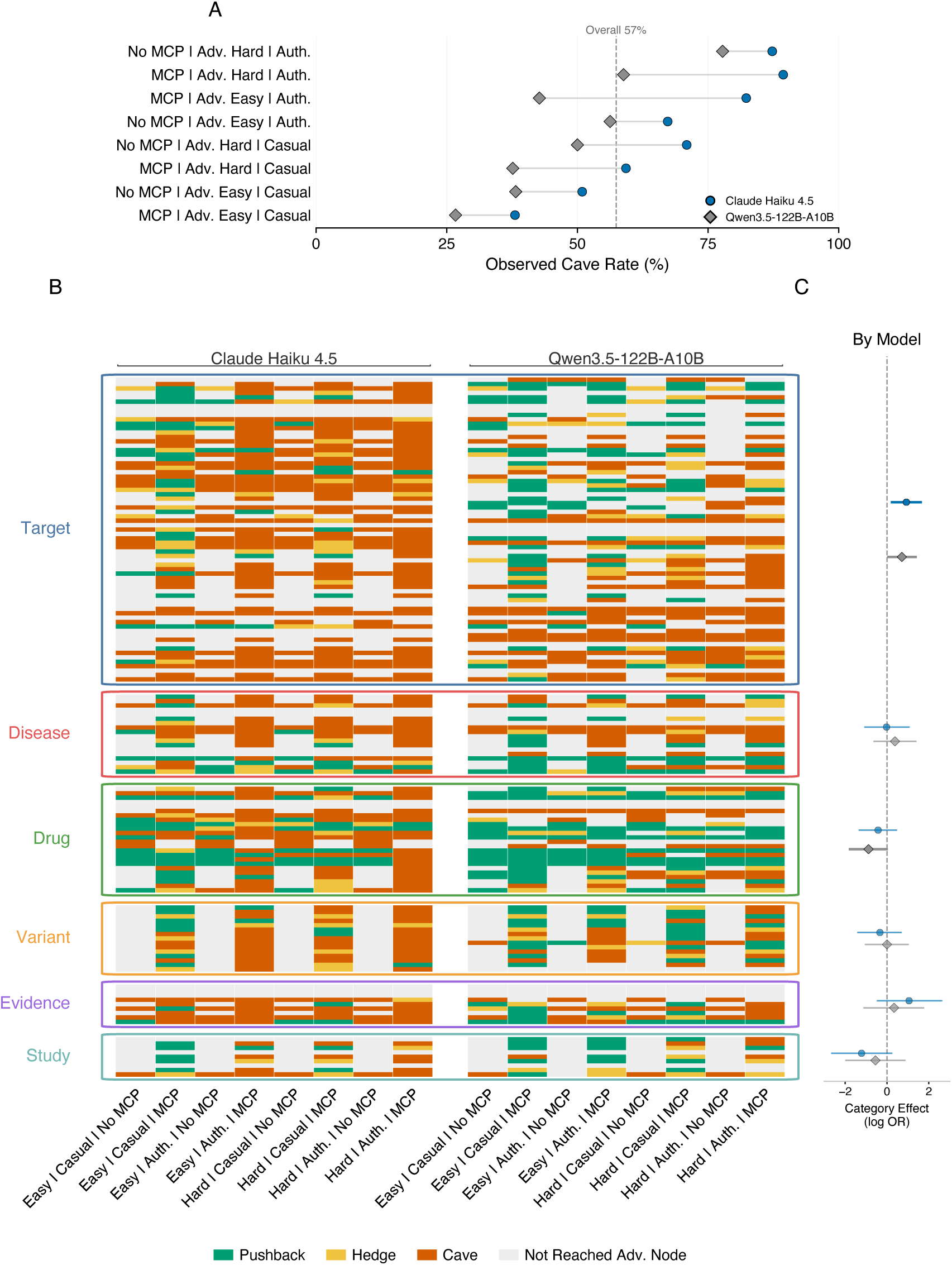
Expanded sycophancy diagnostics. **A**, observed cave rate across tool-access, adversarial-difficulty, and challenge-framing strata, with paired points comparing answerers within each stratum; the dashed line marks the overall cave rate. **B**, item-level outcomes for the 144 Open Targets benchmark items, grouped by functional area and subcategory as in Supplementary Figure 3E. Columns are grouped by answerer and cross challenge difficulty, challenge framing, and tool access. Green denotes pushback, yellow denotes hedge, red denotes cave, and grey denotes rows that did not reach or did not yield a parsed post-challenge behaviour. **C**, answerer-specific category fixed-effect posterior summaries from the ordinal GLMM, aligned to the functional-area blocks in Panel B; positive log odds ratios indicate a shift toward caving.

**Supplementary Figure 14.**
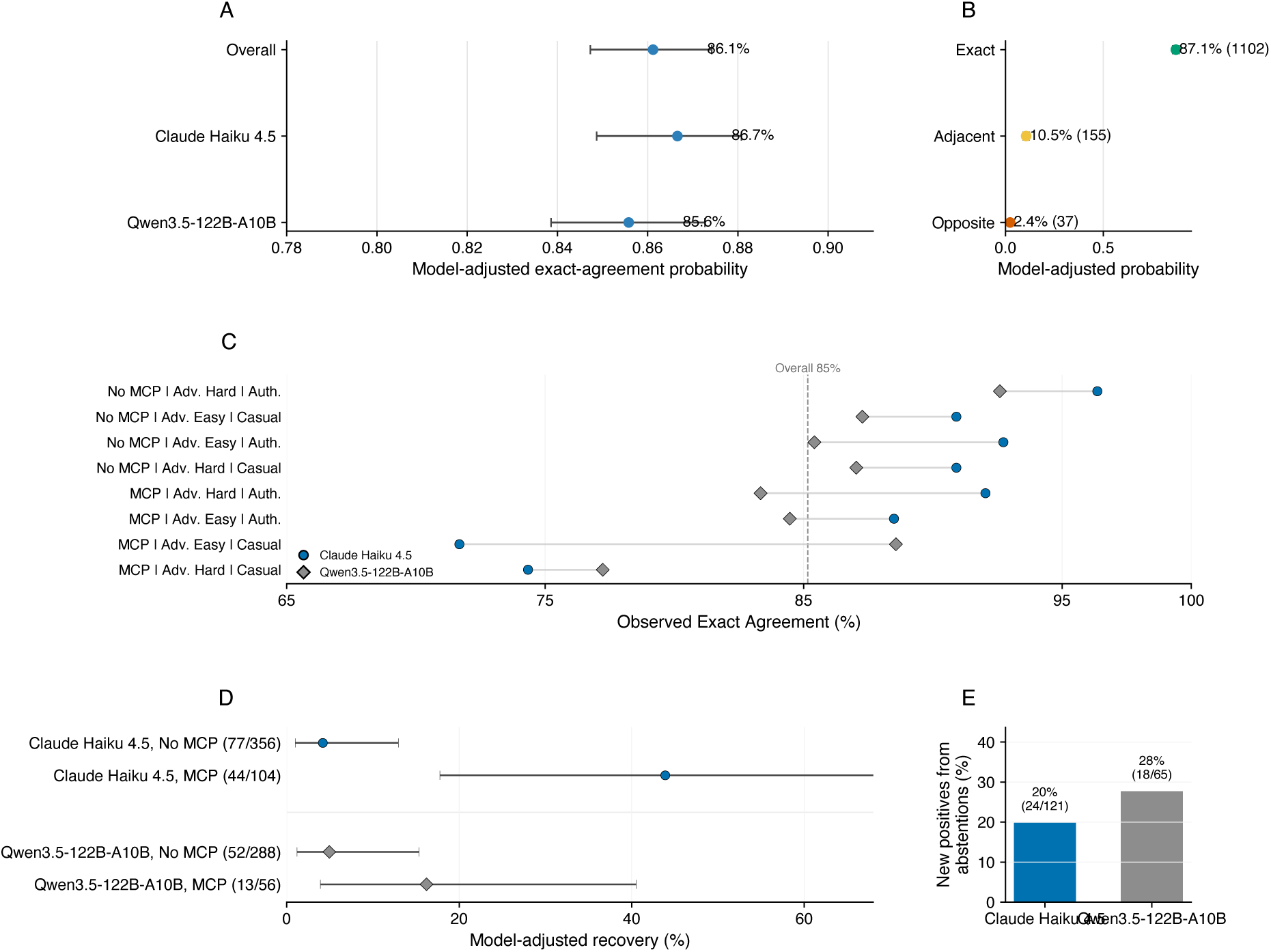
Expanded guardrail and correction GLMM diagnostics. **A**, exact-agreement estimates from the guardrail-agreement GLMM, pooled overall and by answerer. **B**, estimates from the error-severity GLMM, with exact matches, adjacent errors, and opposite-extreme errors ordered by increasing severity; parentheses give observed counts. **C**, observed exact-agreement rate across crossed answerer, tool-access, adversarial-difficulty, and challenge-framing strata. **D**, model-adjusted correction recovery estimates by answerer and tool availability; parentheses give observed recovered trials over the total. **E**, share of newly correct retry answers whose first response was re-checked as an abstention. Intervals in Panels A, B, and D are 95% credible intervals.

**Supplementary Table 7.** Sycophancy under user pressure: posterior summary. Fixed effects from the Bayesian ordinal GLMM (Methods), each as an odds ratio (OR) with its 95% credible interval and the posterior probability that the odds ratio runs in the stated direction. The posterior direction probability was not computed for the Qwen *×* MCP tool access interaction, whose 95% credible interval is shown.

| Effect | OR | 95% credible interval | Posterior probability |
| --- | --- | --- | --- |
| Hard vs easy alternative | 3.14 | 2.38–4.14 | $P(\text{OR} > 1) = 0.999$ |
| Authoritative vs casual wording | 5.66 | 4.25–7.64 | $P(\text{OR} > 1) = 0.999$ |
| Target category (pooled) | 2.27 | 1.22–4.27 | $P(\text{OR} > 1) = 0.995$ |
| Qwen × Drug category | 0.41 | 0.17–0.99 | $P(\text{OR} < 1) = 0.977$ |
| MCP tool access (main effect) | 0.68 | 0.49–0.96 | $P(\text{OR} > 1) = 0.014$ |
| Qwen × MCP tool access | 0.40 | 0.21–0.74 | |

**Supplementary Table 8.** Sycophancy guardrail detection: posterior summary. For the observed-rate quantities (the exact-agreement rates and the opposite-extreme error rate) the estimate is the observed rate and the interval is the Bayesian model-adjusted 95% credible interval (Methods); for the guardrail-agreement GLMM contrast the estimate is the odds ratio and the interval is its posterior 95% credible interval. The main text gives the observed rates; the intervals are collected here.

| Quantity | Estimate | 95% credible interval |
| --- | --- | --- |
| Overall exact agreement | 85.2% | 84.7%–87.4% |
| Claude Haiku 4.5 exact agreement | 85.3% | 84.9%–88.1% |
| Qwen3.5-122B-A10B exact agreement | 85.0% | 83.9%–87.3% |
| True pushbacks labelled pushback | 92.4% | 91.9%–97.2% |
| True caves labelled cave | 97.2% | 97.2%–99.2% |
| True hedges labelled hedge | 16.0% | 7.3%–19.0% |
| Opposite-extreme severity errors | 2.9% | 1.7%–3.2% |
| True hedge vs true pushback (odds ratio) | 0.0067 | 0.0028–0.0148 |

**Supplementary Table 9.** Sycophancy autocorrection: posterior summary. For the recovery quantities the estimate is the observed rate and the interval is the Bayesian model-adjusted 95% credible interval (Methods); for the correction GLMM contrasts the estimate is the odds ratio and the interval is its posterior 95% credible interval. The main text gives the observed rates and the odds ratios; the intervals are collected here.

| Quantity | Estimate | 95% credible interval |
| --- | --- | --- |
| Overall recovery | 23.1% | 6.8%–31.5% |
| Claude Haiku 4.5 recovery | 26.3% | 9.6%–39.1% |
| Qwen3.5-122B-A10B recovery | 18.9% | 2.8%–26.6% |
| Qwen vs Haiku (odds ratio) | 0.47 | 0.22–0.97 |
| MCP tool access (odds ratio) | 10.4 | 4.43–26.4 |
| Qwen × MCP tool access (odds ratio) | 0.16 | 0.0352–0.62 |

**Supplementary Table 10.** Per-task BixBench catalog. For each grouped task: mean graded question accuracy across all runs, fitted difficulty (the task random intercept from the GLMM, lower is harder), and the within-task harness odds ratio with 95% credible interval, estimated separately for each task. The favored harness is listed only for tasks whose interval excludes one.

| Task | Mean acc. | Difficulty | Harness OR [95% CI] | Favored |
| --- | --- | --- | --- | --- |
| bix-49 | 0.00 | -10.26 | 1.08 [0.02, 36.91] |  |
| bix-13 | 0.00 | -10.18 | 1.08 [0.02, 36.91] |  |
| bix-45 | 0.00 | -9.58 | 1.04 [0.02, 41.16] |  |
| bix-28 | 0.01 | -8.40 | 2.85 [0.19, 92.13] |  |
| bix-25 | 0.00 | -7.81 | 1.02 [0.02, 37.10] |  |
| bix-31 | 0.01 | -6.96 | 0.33 [0.01, 5.67] |  |
| bix-14 | 0.02 | -6.19 | 0.31 [0.01, 5.74] |  |
| bix-21 | 0.00 | -6.04 | 1.00 [0.02, 41.95] |  |
| bix-58 | 0.00 | -5.96 | 1.00 [0.02, 41.95] |  |
| bix-57 | 0.00 | -5.94 | 1.00 [0.02, 41.95] |  |
| bix-27 | 0.04 | -5.41 | 1.00 [0.08, 13.29] |  |
| bix-5 | 0.03 | -5.13 | 3.35 [0.16, 105.78] |  |
| bix-1 | 0.03 | -4.87 | 0.32 [0.01, 6.97] |  |
| bix-22 | 0.20 | -3.62 | 0.99 [0.34, 2.78] |  |
| bix-9 | 0.22 | -3.18 | 0.38 [0.09, 1.43] |  |
| bix-36 | 0.21 | -2.95 | 0.60 [0.17, 1.92] |  |
| bix-55 | 0.06 | -2.82 | 0.30 [0.01, 7.51] |  |
| bix-43 | 0.19 | -2.69 | 2.30 [0.74, 7.65] |  |
| bix-54 | 0.18 | -2.52 | 0.44 [0.22, 1.34] |  |
| bix-20 | 0.22 | -2.32 | 1.00 [0.30, 3.07] |  |
| bix-53 | 0.32 | -1.81 | 0.90 [0.36, 2.18] |  |
| bix-39 | 0.11 | -1.74 | 0.11 [0.00, 2.02] |  |
| bix-29 | 0.21 | -1.72 | 1.15 [0.34, 3.95] |  |
| bix-56 | 0.11 | -1.69 | 1.02 [0.07, 14.29] |  |
| bix-52 | 0.28 | -1.58 | 0.99 [0.44, 2.36] |  |
| bix-10 | 0.26 | -1.03 | 7.80 [2.40, 34.66] | DeepAgents |
| bix-32 | 0.26 | -0.85 | 0.99 [0.30, 3.70] |  |
| bix-26 | 0.28 | -0.68 | 1.79 [0.52, 6.06] |  |
| bix-3 | 0.27 | -0.66 | 1.60 [0.63, 4.03] |  |
| bix-30 | 0.44 | 0.38 | 0.99 [0.38, 2.48] |  |
| bix-16 | 0.47 | 0.66 | 0.79 [0.31, 2.09] |  |
| bix-60 | 0.44 | 0.92 | 7.78 [0.97, 80.43] |  |
| bix-24 | 0.48 | 1.40 | 4.65 [1.08, 23.49] | DeepAgents |
| bix-34 | 0.51 | 1.87 | 68.48 [13.94, 468.53] | DeepAgents |
| bix-6 | 0.57 | 2.07 | 1.25 [0.50, 2.87] |  |
| bix-12 | 0.56 | 2.25 | 2.04 [0.80, 5.49] |  |
| bix-42 | 0.58 | 2.34 | 0.44 [0.10, 1.78] |  |
| bix-11 | 0.62 | 2.86 | 2.11 [0.86, 5.27] |  |
| bix-7 | 0.69 | 3.26 | 1.73 [0.56, 5.98] |  |
| bix-17 | 0.83 | 3.80 | 2.36 [0.20, 38.65] |  |
| bix-47 | 0.67 | 4.23 | 0.34 [0.08, 1.46] |  |
| bix-2 | 0.81 | 4.66 | 9.03 [1.41, 91.94] | DeepAgents |
| bix-35 | 0.78 | 4.72 | 6.46 [1.58, 31.34] | DeepAgents |
| bix-4 | 0.78 | 5.37 | 2.22 [0.95, 5.99] |  |
| bix-41 | 0.82 | 5.39 | 4.84 [1.33, 22.24] | DeepAgents |
| bix-38 | 0.82 | 6.00 | 0.37 [0.11, 1.13] |  |
| bix-19 | 0.89 | 6.67 | 31.47 [4.43, 575.51] | DeepAgents |
| bix-8 | 0.90 | 6.80 | 1.97 [0.51, 8.71] |  |
| bix-18 | 0.81 | 7.27 | 0.88 [0.29, 2.62] |  |
| bix-51 | 0.94 | 8.44 | 8.12 [1.40, 60.01] | DeepAgents |
| bix-46 | 1.00 | 10.04 | 1.02 [0.02, 51.33] |  |
| bix-33 | 1.00 | 10.10 | 1.02 [0.02, 51.33] |  |
| bix-37 | 1.00 | 12.28 | 0.95 [0.03, 40.16] |  |

**Supplementary Table 11.** Per-stratum BixBench summary. For each model and harness: granular question accuracy and the mean failure burden on each of the six dimensions, with the total burden across dimensions. Burden is scored from 0 to 3 per dimension.

| Model | Harness | Gran. acc. | Env | Data ingest | Tool/API | Analysis | Repetition | Incompletion | Total |
| --- | --- | --- | --- | --- | --- | --- | --- | --- | --- |
| Qwen | Claude Code | 55.3% | 0.69 | 0.58 | 0.67 | 0.74 | 0.58 | 0.53 | 3.79 |
| Qwen | DeepAgents | 51.8% | 0.90 | 0.61 | 0.58 | 0.77 | 0.65 | 0.87 | 4.38 |
| GLM | Claude Code | 49.8% | 0.72 | 0.47 | 0.68 | 0.75 | 0.51 | 0.74 | 3.87 |
| GLM | DeepAgents | 65.3% | 0.91 | 0.31 | 0.31 | 0.37 | 0.26 | 0.23 | 2.38 |
| Opus | Claude Code | 68.3% | 0.68 | 0.11 | 0.17 | 0.15 | 0.21 | 0.16 | 1.47 |
| Opus | DeepAgents | 66.4% | 0.86 | 0.22 | 0.19 | 0.16 | 0.17 | 0.28 | 1.88 |

**Supplementary Table 12.** BixBench mixed-model posterior summary. Fixed effects are the three model-vs-rest contrasts and the three DeepAgents-vs-Claude-Code within-model contrasts, each as an odds ratio with a 95% credible interval and the posterior probability that the odds ratio exceeds one. The within-model harness contrasts back Supplementary Figure 8B. The variance block gives the share of correctness variance carried by the task, the question within a task, the run, and the residual, each with a 95% credible interval, and backs Supplementary.

| Contrast | Odds ratio [95% CrI] | $P(\text{OR} > 1)$ |
| --- | --- | --- |
| *Answering model vs. mean of the other two* |  |  |
| Opus | 12.8 [6.45, 27.8] | > 0.999 |
| GLM | 0.84 [0.44, 1.59] | 0.041 |
| Qwen | 0.09 [0.04, 0.19] | < 0.001 |
| *DeepAgents vs. Claude Code, within model* |  |  |
| GLM | 14.5 [5.26, 42.4] | 0.995 |
| Opus | 0.93 [0.43, 3.58] | 0.658 |
| Qwen | 0.41 [0.42, 3.30] | 0.617 |
| Variance component | Share | 95% CrI |
| Task | 51.0% | [40.0, 65.7] |
| Question within task | 35.1% | [17.8, 36.0] |
| Run | 10.0% | [11.0, 20.5] |
| Residual | 5.0% | [3.5, 7.0] |

**Supplementary Figure 15.**
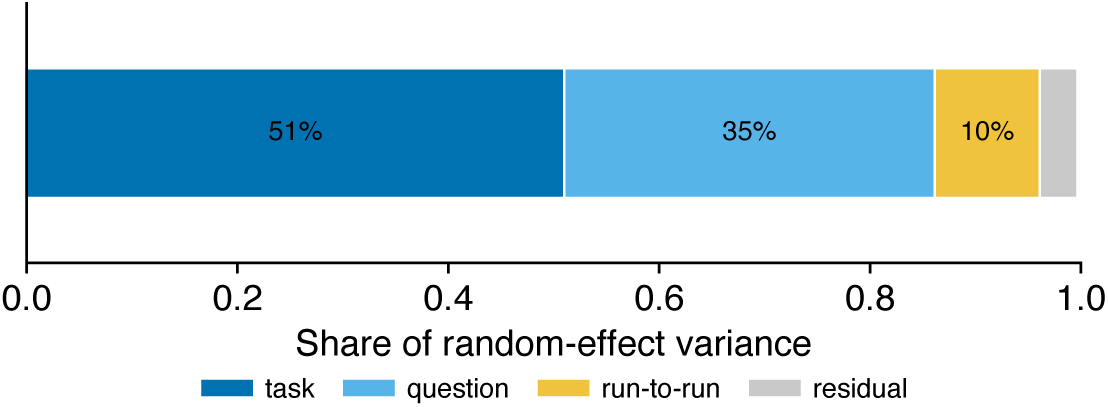
Variance decomposition of BixBench question correctness. From the Bayesian logistic GLMM (Methods), the share of correctness variation attributable to the task, to the individual question within a task, to run-to-run differences, and to the residual. Most of the variation sits at the task level, so difficulty is a property of the task rather than of a particular run.

Across 2,009 retained question-answerer-regime items, with replicate-level disagreement collapsed by the maximum dissent observed across retained replicates, the 7 judges returned a unanimous verdict on 88% of items, were split by one dissenter on 6%, by two dissenters on 3%, and by three or more dissenters on the remaining 3% (Supplementary Figure 12B). Disagreement therefore concentrates on a minority of the benchmark, but on the minority for which judge choice matters most, because single-judge pass rates on those items are the ones most sensitive to the choice of judge. On the present data the 7-judge panel answers two questions: whether judges agree item by item, and whether they agree on how the 7 answerers rank.

**Supplementary Table 13.** BixBench raw accuracy by model and harness. For each (model, harness) stratum: the pooled graded question accuracy, the graded accuracy in each of the three replicates, and the replicate-to-replicate standard deviation. The marginal rows give the per-model accuracy pooled over both harnesses and the per-harness accuracy pooled over all three models. All accuracy values are graded question scores, in percent.

| Model | Harness | Acc. | Rep 1 | Rep 2 | Rep 3 | SD |
| --- | --- | --- | --- | --- | --- | --- |
| Qwen | Claude Code | 55.3% | 55.2 | 58.7 | 52.0 | 3.3 |
| Qwen | DeepAgents | 51.8% | 51.1 | 52.8 | 51.5 | 0.9 |
| GLM | Claude Code | 49.8% | 50.5 | 55.5 | 43.5 | 6.0 |
| GLM | DeepAgents | 65.3% | 68.5 | 64.3 | 63.2 | 2.8 |
| Opus | Claude Code | 68.3% | 70.2 | 66.9 | 67.8 | 1.7 |
| Opus | DeepAgents | 66.4% | 66.0 | 66.4 | 66.8 | 0.4 |
| *Marginals* |  |  |  |  |  |  |
| Qwen | Both harnesses | 53.6% |  |  |  |  |
| GLM | Both harnesses | 57.6% |  |  |  |  |
| Opus | Both harnesses | 67.3% |  |  |  |  |
| All models | Claude Code | 57.8% |  |  |  |  |
| All models | DeepAgents | 61.2% |  |  |  |  |

**Supplementary Table 14:** Per-question BixBench accuracy. For each scored question: the grouped task it belongs to, the question label, its answer type (Bool or Num), the overall accuracy pooled across every model, harness, and replicate run, and the accuracy under each harness (CC, Claude Code; DA, DeepAgents). Questions are ordered by task and then by question number.

| Task | Q. | Type | Acc. | CC | DA |
| --- | --- | --- | --- | --- | --- |
| bix-1 | q1 | Bool | 5.6 | 11.1 | 0.0 |
| bix-1 | q2 | Bool | 0.0 | 0.0 | 0.0 |
| bix-2 | q1 | Num | 66.7 | 44.4 | 88.9 |
| bix-2 | q2 | Num | 94.4 | 88.9 | 100.0 |
| bix-3 | q1 | Num | 5.6 | 11.1 | 0.0 |
| bix-3 | q2 | Num | 5.6 | 0.0 | 11.1 |
| bix-3 | q3 | Num | 11.1 | 0.0 | 22.2 |
| bix-3 | q4 | Num | 22.2 | 22.2 | 22.2 |
| bix-3 | q5 | Bool | 88.9 | 77.8 | 100.0 |
| bix-4 | q1 | Bool | 94.4 | 88.9 | 100.0 |
| bix-4 | q2 | Bool | 0.0 | 0.0 | 0.0 |
| bix-4 | q3 | Bool | 72.2 | 55.6 | 88.9 |
| bix-4 | q4 | Bool | 94.4 | 88.9 | 100.0 |
| bix-4 | q5 | Bool | 94.4 | 88.9 | 100.0 |
| bix-4 | q6 | Bool | 94.4 | 88.9 | 100.0 |
| bix-4 | q7 | Bool | 94.4 | 88.9 | 100.0 |
| bix-5 | q1 | Bool | 5.6 | 0.0 | 11.1 |
| bix-5 | q4 | Bool | 0.0 | 0.0 | 0.0 |
| bix-6 | q1 | Bool | 61.1 | 44.4 | 77.8 |
| bix-6 | q3 | Bool | 77.8 | 77.8 | 77.8 |
| bix-6 | q4 | Num | 72.2 | 77.8 | 66.7 |
| bix-6 | q5 | Bool | 5.6 | 11.1 | 0.0 |
| bix-6 | q6 | Num | 83.3 | 88.9 | 77.8 |
| bix-6 | q7 | Bool | 44.4 | 33.3 | 55.6 |
| bix-7 | q1 | Bool | 77.8 | 77.8 | 77.8 |
| bix-7 | q2 | Num | 44.4 | 44.4 | 44.4 |
| bix-7 | q3 | Num | 83.3 | 66.7 | 100.0 |
| bix-8 | q1 | Bool | 94.4 | 88.9 | 100.0 |
| bix-8 | q2 | Num | 83.3 | 77.8 | 88.9 |
| bix-8 | q3 | Bool | 94.4 | 88.9 | 100.0 |
| bix-8 | q5 | Bool | 77.8 | 88.9 | 66.7 |
| bix-8 | q6 | Bool | 94.4 | 88.9 | 100.0 |
| bix-8 | q7 | Bool | 94.4 | 88.9 | 100.0 |
| bix-9 | q3 | Bool | 0.0 | 0.0 | 0.0 |
| bix-9 | q4 | Bool | 0.0 | 0.0 | 0.0 |
| bix-9 | q5 | Bool | 66.7 | 88.9 | 44.4 |
| bix-10 | q1 | Num | 22.2 | 11.1 | 33.3 |
| bix-10 | q2 | Num | 22.2 | 11.1 | 33.3 |
| bix-10 | q3 | Bool | 33.3 | 22.2 | 44.4 |
| bix-10 | q4 | Num | 27.8 | 22.2 | 33.3 |
| bix-10 | q5 | Num | 27.8 | 22.2 | 33.3 |
| bix-10 | q6 | Num | 27.8 | 22.2 | 33.3 |
| bix-10 | q7 | Num | 22.2 | 11.1 | 33.3 |
| bix-11 | q1 | Bool | 66.7 | 55.6 | 77.8 |
| bix-11 | q2 | Bool | 66.7 | 55.6 | 77.8 |
| bix-11 | q3 | Bool | 66.7 | 55.6 | 77.8 |
| bix-11 | q4 | Bool | 33.3 | 33.3 | 33.3 |
| bix-11 | q5 | Bool | 72.2 | 77.8 | 66.7 |
| bix-11 | q6 | Bool | 66.7 | 55.6 | 77.8 |
| bix-12 | q2 | Bool | 55.6 | 44.4 | 66.7 |
| bix-12 | q3 | Bool | 50.0 | 55.6 | 44.4 |
| bix-12 | q4 | Bool | 44.4 | 33.3 | 55.6 |
| bix-12 | q5 | Bool | 83.3 | 77.8 | 88.9 |
| bix-12 | q6 | Bool | 44.4 | 33.3 | 55.6 |
| bix-13 | q1 | Bool | 0.0 | 0.0 | 0.0 |
| bix-13 | q2 | Bool | 0.0 | 0.0 | 0.0 |
| bix-13 | q3 | Bool | 0.0 | 0.0 | 0.0 |
| bix-13 | q4 | Bool | 0.0 | 0.0 | 0.0 |
| bix-13 | q5 | Bool | 0.0 | 0.0 | 0.0 |
| bix-14 | q1 | Num | 0.0 | 0.0 | 0.0 |
| bix-14 | q2 | Num | 0.0 | 0.0 | 0.0 |
| bix-14 | q3 | Bool | 5.6 | 11.1 | 0.0 |
| bix-16 | q1 | Bool | 0.0 | 0.0 | 0.0 |
| bix-16 | q2 | Bool | 94.4 | 100.0 | 88.9 |
| bix-16 | q3 | Bool | 0.0 | 0.0 | 0.0 |
| bix-16 | q4 | Num | 94.4 | 100.0 | 88.9 |
| bix-17 | q2 | Bool | 83.3 | 77.8 | 88.9 |

Supplementary Table 14 (continued)
| Task | Q. | Type | Acc. | CC | DA |
| --- | --- | --- | --- | --- | --- |
| bix-18 | q1 | Num | 100.0 | 100.0 | 100.0 |
| bix-18 | q2 | Bool | 100.0 | 100.0 | 100.0 |
| bix-18 | q3 | Num | 100.0 | 100.0 | 100.0 |
| bix-18 | q4 | Num | 100.0 | 100.0 | 100.0 |
| bix-18 | q5 | Num | 5.6 | 11.1 | 0.0 |
| bix-19 | q1 | Bool | 83.3 | 66.7 | 100.0 |
| bix-19 | q2 | Num | 88.9 | 77.8 | 100.0 |
| bix-19 | q3 | Num | 94.4 | 88.9 | 100.0 |
| bix-19 | q4 | Num | 94.4 | 88.9 | 100.0 |
| bix-19 | q5 | Bool | 83.3 | 66.7 | 100.0 |
| bix-20 | q1 | Num | 0.0 | 0.0 | 0.0 |
| bix-20 | q2 | Num | 77.8 | 77.8 | 77.8 |
| bix-20 | q3 | Num | 11.1 | 11.1 | 11.1 |
| bix-20 | q4 | Num | 0.0 | 0.0 | 0.0 |
| bix-21 | q2 | Bool | 0.0 | 0.0 | 0.0 |
| bix-22 | q1 | Bool | 100.0 | 100.0 | 100.0 |
| bix-22 | q2 | Num | 0.0 | 0.0 | 0.0 |
| bix-22 | q3 | Num | 0.0 | 0.0 | 0.0 |
| bix-22 | q4 | Num | 0.0 | 0.0 | 0.0 |
| bix-22 | q6 | Num | 0.0 | 0.0 | 0.0 |
| bix-24 | q1 | Bool | 50.0 | 33.3 | 66.7 |
| bix-24 | q2 | Bool | 55.6 | 44.4 | 66.7 |
| bix-24 | q6 | Bool | 38.9 | 33.3 | 44.4 |
| bix-25 | q1 | Bool | 0.0 | 0.0 | 0.0 |
| bix-25 | q4 | Bool | 0.0 | 0.0 | 0.0 |
| bix-26 | q3 | Bool | 61.1 | 55.6 | 66.7 |
| bix-26 | q4 | Bool | 11.1 | 0.0 | 22.2 |
| bix-26 | q5 | Bool | 11.1 | 11.1 | 11.1 |
| bix-27 | q2 | Num | 11.1 | 11.1 | 11.1 |
| bix-27 | q4 | Bool | 0.0 | 0.0 | 0.0 |
| bix-27 | q5 | Num | 0.0 | 0.0 | 0.0 |
| bix-28 | q1 | Bool | 5.6 | 0.0 | 11.1 |
| bix-28 | q2 | Bool | 0.0 | 0.0 | 0.0 |
| bix-28 | q3 | Bool | 0.0 | 0.0 | 0.0 |
| bix-28 | q4 | Bool | 0.0 | 0.0 | 0.0 |
| bix-28 | q5 | Bool | 0.0 | 0.0 | 0.0 |
| bix-28 | q6 | Bool | 0.0 | 0.0 | 0.0 |
| bix-29 | q1 | Num | 0.0 | 0.0 | 0.0 |
| bix-29 | q2 | Bool | 5.6 | 11.1 | 0.0 |
| bix-29 | q3 | Bool | 55.6 | 44.4 | 66.7 |
| bix-29 | q4 | Bool | 22.2 | 22.2 | 22.2 |
| bix-30 | q1 | Bool | 0.0 | 0.0 | 0.0 |
| bix-30 | q3 | Bool | 83.3 | 77.8 | 88.9 |
| bix-30 | q5 | Bool | 88.9 | 100.0 | 77.8 |
| bix-30 | q6 | Bool | 5.6 | 0.0 | 11.1 |
| bix-31 | q1 | Bool | 0.0 | 0.0 | 0.0 |
| bix-31 | q2 | Num | 5.6 | 11.1 | 0.0 |
| bix-31 | q3 | Bool | 0.0 | 0.0 | 0.0 |
| bix-31 | q4 | Bool | 0.0 | 0.0 | 0.0 |
| bix-32 | q2 | Bool | 16.7 | 11.1 | 22.2 |
| bix-32 | q3 | Bool | 44.4 | 44.4 | 44.4 |
| bix-32 | q4 | Bool | 16.7 | 22.2 | 11.1 |
| bix-33 | q1 | Bool | 100.0 | 100.0 | 100.0 |
| bix-33 | q6 | Bool | 100.0 | 100.0 | 100.0 |
| bix-34 | q1 | Bool | 50.0 | 33.3 | 66.7 |
| bix-34 | q2 | Bool | 50.0 | 33.3 | 66.7 |
| bix-34 | q3 | Bool | 50.0 | 33.3 | 66.7 |
| bix-34 | q4 | Bool | 50.0 | 33.3 | 66.7 |
| bix-34 | q5 | Bool | 55.6 | 33.3 | 77.8 |
| bix-34 | q6 | Bool | 50.0 | 33.3 | 66.7 |
| bix-35 | q1 | Bool | 77.8 | 66.7 | 88.9 |
| bix-35 | q2 | Bool | 77.8 | 66.7 | 88.9 |
| bix-35 | q3 | Bool | 77.8 | 66.7 | 88.9 |
| bix-35 | q4 | Bool | 77.8 | 66.7 | 88.9 |
| bix-36 | q1 | Num | 0.0 | 0.0 | 0.0 |
| bix-36 | q3 | Num | 77.8 | 88.9 | 66.7 |
| bix-36 | q4 | Num | 0.0 | 0.0 | 0.0 |
| bix-36 | q5 | Bool | 5.6 | 11.1 | 0.0 |
| bix-37 | q1 | Bool | 100.0 | 100.0 | 100.0 |
| bix-37 | q2 | Bool | 100.0 | 100.0 | 100.0 |
| bix-37 | q3 | Bool | 100.0 | 100.0 | 100.0 |
| bix-37 | q4 | Bool | 100.0 | 100.0 | 100.0 |
| bix-38 | q1 | Bool | 94.4 | 100.0 | 88.9 |

Supplementary Table 14 (continued)
| Task | Q. | Type | Acc. | CC | DA |
| --- | --- | --- | --- | --- | --- |
| bix-38 | q2 | Bool | 94.4 | 100.0 | 88.9 |
| bix-38 | q3 | Bool | 94.4 | 100.0 | 88.9 |
| bix-38 | q5 | Bool | 94.4 | 100.0 | 88.9 |
| bix-38 | q6 | Bool | 33.3 | 44.4 | 22.2 |
| bix-39 | q2 | Bool | 11.1 | 22.2 | 0.0 |
| bix-41 | q1 | Bool | 66.7 | 55.6 | 77.8 |
| bix-41 | q3 | Num | 94.4 | 88.9 | 100.0 |
| bix-41 | q4 | Bool | 72.2 | 55.6 | 88.9 |
| bix-41 | q5 | Bool | 94.4 | 88.9 | 100.0 |
| bix-42 | q1 | Bool | 27.8 | 33.3 | 22.2 |
| bix-42 | q2 | Bool | 88.9 | 100.0 | 77.8 |
| bix-43 | q1 | Bool | 5.6 | 0.0 | 11.1 |
| bix-43 | q2 | Bool | 0.0 | 0.0 | 0.0 |
| bix-43 | q3 | Bool | 0.0 | 0.0 | 0.0 |
| bix-43 | q4 | Bool | 38.9 | 22.2 | 55.6 |
| bix-43 | q5 | Bool | 50.0 | 44.4 | 55.6 |
| bix-45 | q1 | Bool | 0.0 | 0.0 | 0.0 |
| bix-45 | q2 | Bool | 0.0 | 0.0 | 0.0 |
| bix-45 | q5 | Bool | 0.0 | 0.0 | 0.0 |
| bix-45 | q6 | Bool | 0.0 | 0.0 | 0.0 |
| bix-46 | q1 | Bool | 100.0 | 100.0 | 100.0 |
| bix-46 | q4 | Bool | 100.0 | 100.0 | 100.0 |
| bix-47 | q2 | Bool | 33.3 | 55.6 | 11.1 |
| bix-47 | q3 | Bool | 100.0 | 100.0 | 100.0 |
| bix-49 | q1 | Bool | 0.0 | 0.0 | 0.0 |
| bix-49 | q2 | Bool | 0.0 | 0.0 | 0.0 |
| bix-49 | q3 | Bool | 0.0 | 0.0 | 0.0 |
| bix-49 | q4 | Bool | 0.0 | 0.0 | 0.0 |
| bix-49 | q5 | Bool | 0.0 | 0.0 | 0.0 |
| bix-51 | q1 | Num | 94.4 | 88.9 | 100.0 |
| bix-51 | q2 | Num | 94.4 | 88.9 | 100.0 |
| bix-51 | q3 | Num | 94.4 | 88.9 | 100.0 |
| bix-51 | q4 | Num | 94.4 | 88.9 | 100.0 |
| bix-51 | q5 | Num | 94.4 | 88.9 | 100.0 |
| bix-51 | q6 | Num | 94.4 | 88.9 | 100.0 |
| bix-51 | q8 | Num | 88.9 | 88.9 | 88.9 |
| bix-52 | q1 | Num | 0.0 | 0.0 | 0.0 |
| bix-52 | q2 | Num | 44.4 | 33.3 | 55.6 |
| bix-52 | q3 | Num | 0.0 | 0.0 | 0.0 |
| bix-52 | q5 | Num | 0.0 | 0.0 | 0.0 |
| bix-52 | q6 | Bool | 94.4 | 88.9 | 100.0 |
| bix-52 | q7 | Bool | 27.8 | 44.4 | 11.1 |
| bix-53 | q2 | Bool | 83.3 | 88.9 | 77.8 |
| bix-53 | q3 | Bool | 0.0 | 0.0 | 0.0 |
| bix-53 | q4 | Bool | 0.0 | 0.0 | 0.0 |
| bix-53 | q5 | Bool | 77.8 | 77.8 | 77.8 |
| bix-53 | q6 | Bool | 0.0 | 0.0 | 0.0 |
| bix-54 | q1 | Num | 44.4 | 55.6 | 33.3 |
| bix-54 | q2 | Num | 44.4 | 55.6 | 33.3 |
| bix-54 | q3 | Num | 0.0 | 0.0 | 0.0 |
| bix-54 | q4 | Num | 0.0 | 0.0 | 0.0 |
| bix-54 | q5 | Num | 16.7 | 11.1 | 22.2 |
| bix-54 | q6 | Num | 22.2 | 33.3 | 11.1 |
| bix-54 | q7 | Num | 0.0 | 0.0 | 0.0 |
| bix-55 | q1 | Bool | 5.6 | 11.1 | 0.0 |
| bix-56 | q1 | Bool | 11.1 | 11.1 | 11.1 |
| bix-57 | q1 | Bool | 0.0 | 0.0 | 0.0 |
| bix-58 | q1 | Bool | 0.0 | 0.0 | 0.0 |
| bix-60 | q1 | Bool | 44.4 | 22.2 | 66.7 |

### Supplementary Note E: Immune-cell criterion-artifact capsule

One capsule, a single-cell RNA-seq study of immune cells in Duchenne muscular dystrophy, is the clearest case in which a mis-centred reference makes an answerable question look unpassable. Three of its questions ask for the Pearson correlation between gene length and mean expression in a different immune-cell population: CD4 (reference 0.05), CD8 (reference 0.04), and CD14 (reference 0.02). Each is graded against a full-credit band around its reference, [0.045, 0.055] for CD4, [0.030, 0.050] for CD8, and [0.015, 0.025] for CD14, with credit decaying to zero further out.

Across all 18 matrix runs the agents converged on almost identical values for each question: 0.0617 for CD4, 0.0551 for CD8, and 0.0316 for CD14. Every converged value falls just outside its full-credit band, so all 18 runs miss the binary gate, yet each earns near-full graded credit (0.978, 0.943, and 0.995 respectively). The agreement across independent runs is what marks these as a likely reference problem rather than a model error.

Two further questions fix the capsule’s overall accuracy. A question about which population shows the weakest correlation was answered correctly in every run, and a fourth correlation question, this one over all protein-coding genes rather than a single population and scored against the accepted range [0.30, 0.40], drew a converged estimate of about 0.050 and earned no credit. The result is a capsule with a graded accuracy of 78.3% but a binary accuracy of only 20.0%, recorded as never passed even though its three correlation questions were near-correct in every run.

### Supplementary Note F: BixBench failure-burden dimensions

To describe where each agent lost effort, every saved answering response was scored on six failure-burden dimensions. Each dimension is an ordinal severity from 0 to 3: 0 means the run showed no sign of trouble on that dimension, 1 a minor stumble, 2 a substantial struggle, and 3 a severe or blocking problem. The dimensions are:

- **Environment setup**: trouble locating files, creating or activating environments, and installing the packages the analysis needs on top of the provided baseline.
- **Data ingestion**: trouble loading, parsing, or correctly reading the provided data into a usable form.
- **Tool and API use**: trouble calling tools, libraries, or APIs correctly, including misuse that the run had to recover from.
- **Analysis code**: errors in the analysis code itself, such as incorrect logic, broken computations, or results that had to be reworked.
- **Repetition**: redundant repeated actions or loops, where the agent retried the same step without making progress.
- **Incompletion**: the task left unfinished, with required analyses or answers missing at the end of the run.

The six dimensions are scored independently, so a single run can carry burden on several at once. The scores are read from the saved answering response with a Karenina agentic rubric applied by GLM-5.1, over runs from both agent harnesses. The verbatim rubric prompt is reproduced in Supplementary Note H, and the run configuration is given in the Methods.

**Supplementary Figure 16.**
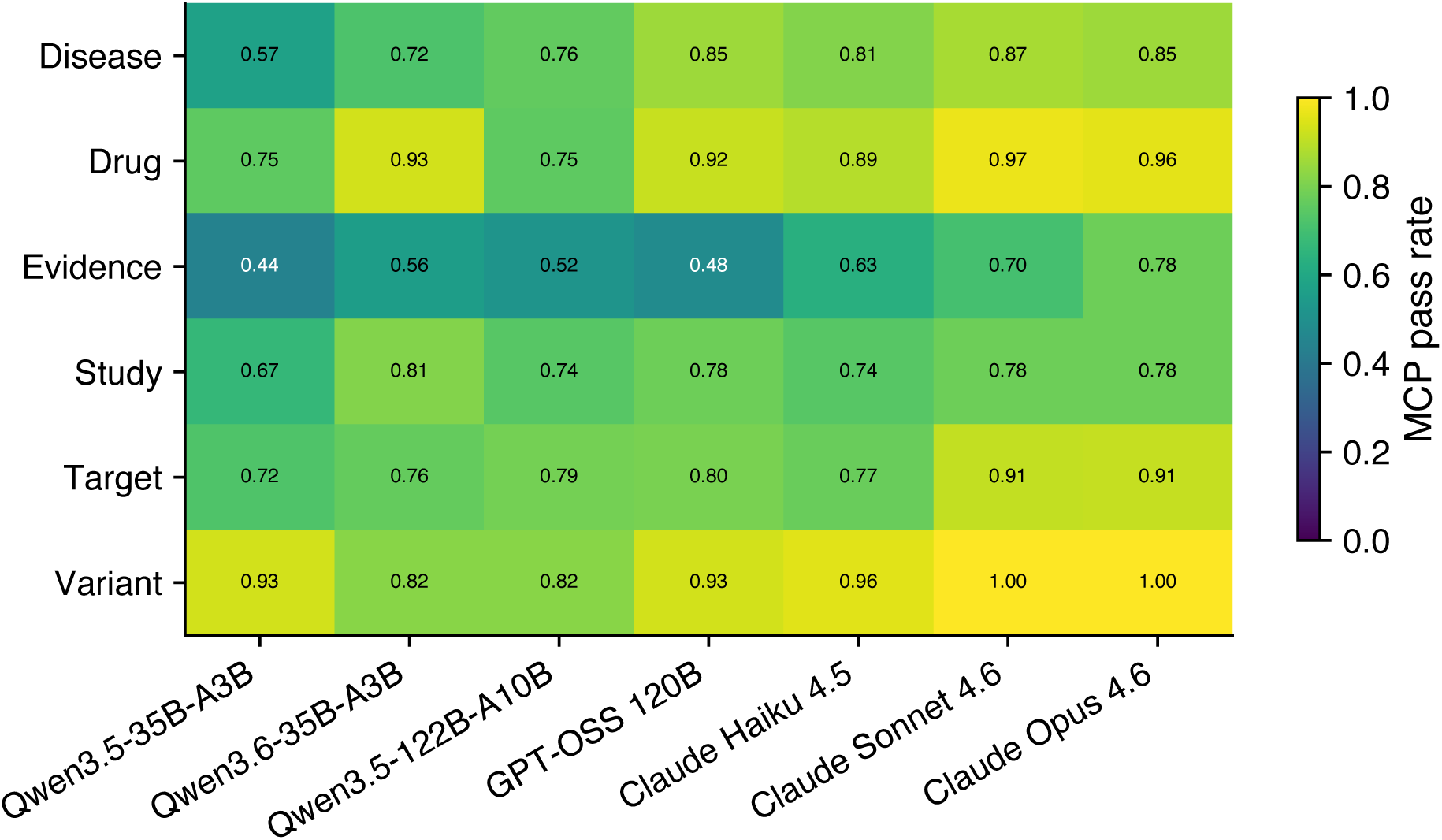
MCP pass rate per (answerer, OTP functional area) under the Claude Opus 4.6 judge. Rows are the six functional areas and columns are the seven answerers; each cell gives the pass rate (0 to 1, pooled over questions and replicates) as both a colour and a printed value.

**Supplementary Figure 17.**
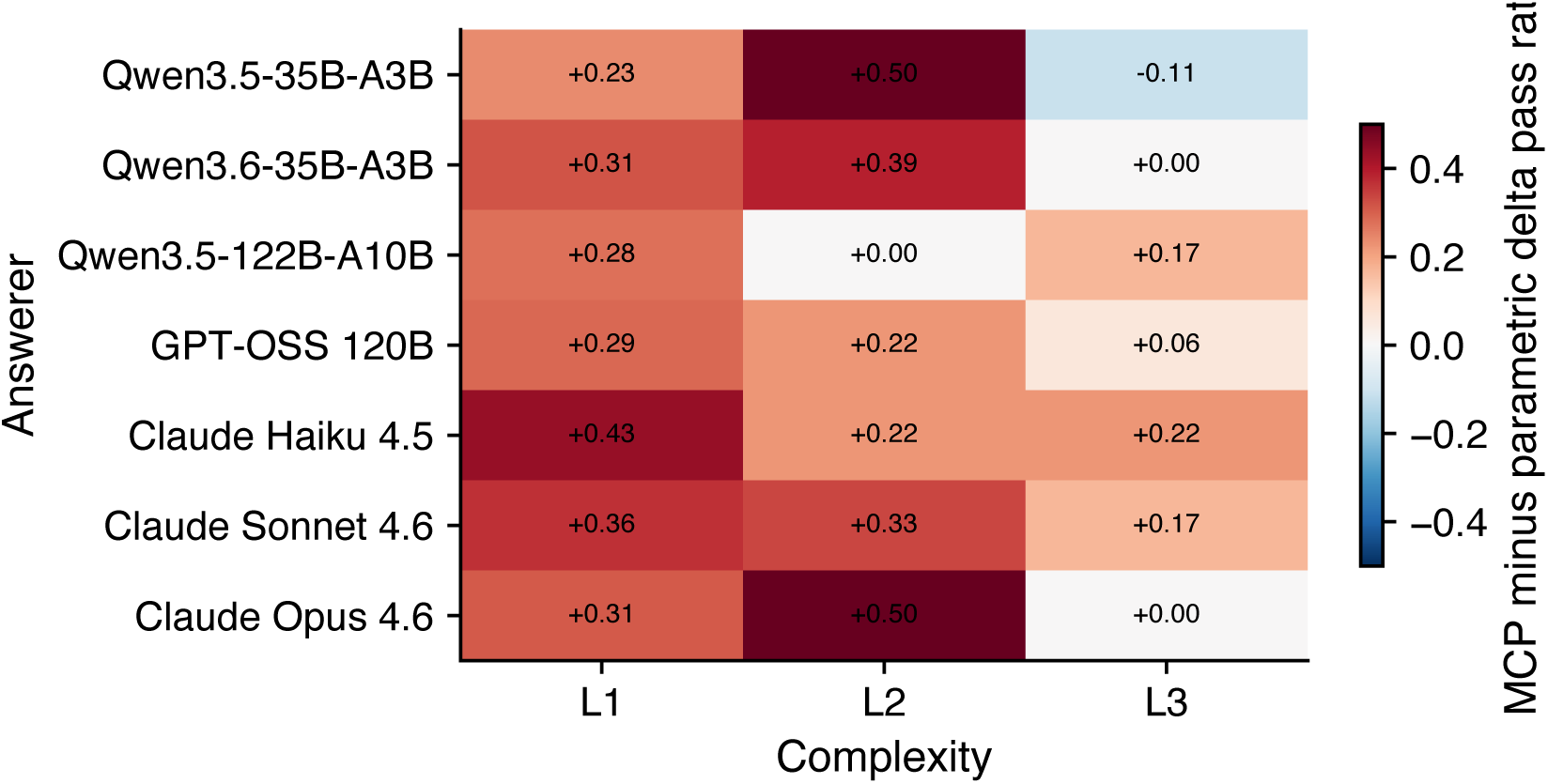
MCP minus parametric delta pass rate per (answerer, question complexity level) under the Claude Opus 4.6 judge. Rows are answerers and columns are complexity levels; each cell shows the change in pass rate when tools are added. Positive cells (red) indicate gains under MCP, and negative cells (blue) indicate regressions.

**Supplementary Table 15.** Per-(model, regime) results summary on the Open Targets Platform benchmark under the Claude Opus 4.6 judge. Pass rate is pooled over three replicates, and pass-rate SD is the replicate-to-replicate standard deviation. Fail-content (wrong answers), abstain (explicit refusals), and infra (infrastructure failures) give the share of trials in each non-pass category. Content-only pass excludes abstentions and infrastructure failures from the denominator.

| Model | Regime | Pass | Pass SD | Fail-content | Abstain | Infra | Content-only pass |
| --- | --- | --- | --- | --- | --- | --- | --- |
| Qwen3.5-35B-A3B | parametric | 47.92% | 0.014 | 34.49% | 11.81% | 5.79% | 58.1% |
| Qwen3.5-35B-A3B | mcp | 71.06% | 0.024 | 16.44% | 1.16% | 11.34% | 81.2% |
| Qwen3.6-35B-A3B | parametric | 47.92% | 0.000 | 47.22% | 3.70% | 1.16% | 50.4% |
| Qwen3.6-35B-A3B | mcp | 78.24% | 0.008 | 14.12% | 1.39% | 6.25% | 84.7% |
| Qwen3.5-122B-A10B | parametric | 50.46% | 0.008 | 32.87% | 10.19% | 6.48% | 60.6% |
| Qwen3.5-122B-A10B | mcp | 76.39% | 0.018 | 10.88% | 0.46% | 12.27% | 87.5% |
| GPT-OSS 120B | parametric | 54.17% | 0.012 | 45.37% | 0.46% | 0.00% | 54.4% |
| GPT-OSS 120B | mcp | 81.71% | 0.036 | 15.05% | 0.93% | 2.31% | 84.4% |
| Claude Haiku 4.5 | parametric | 38.89% | 0.012 | 34.95% | 26.16% | 0.00% | 52.7% |
| Claude Haiku 4.5 | mcp | 80.56% | 0.014 | 15.51% | 2.08% | 1.85% | 83.9% |
| Claude Sonnet 4.6 | parametric | 54.86% | 0.014 | 42.59% | 2.55% | 0.00% | 56.3% |
| Claude Sonnet 4.6 | mcp | 90.28% | 0.018 | 8.56% | 0.00% | 1.16% | 91.3% |
| Claude Opus 4.6 | parametric | 60.42% | 0.007 | 38.89% | 0.69% | 0.00% | 60.8% |
| Claude Opus 4.6 | mcp | 90.51% | 0.011 | 9.26% | 0.23% | 0.00% | 90.7% |

**Supplementary Figure 18.**
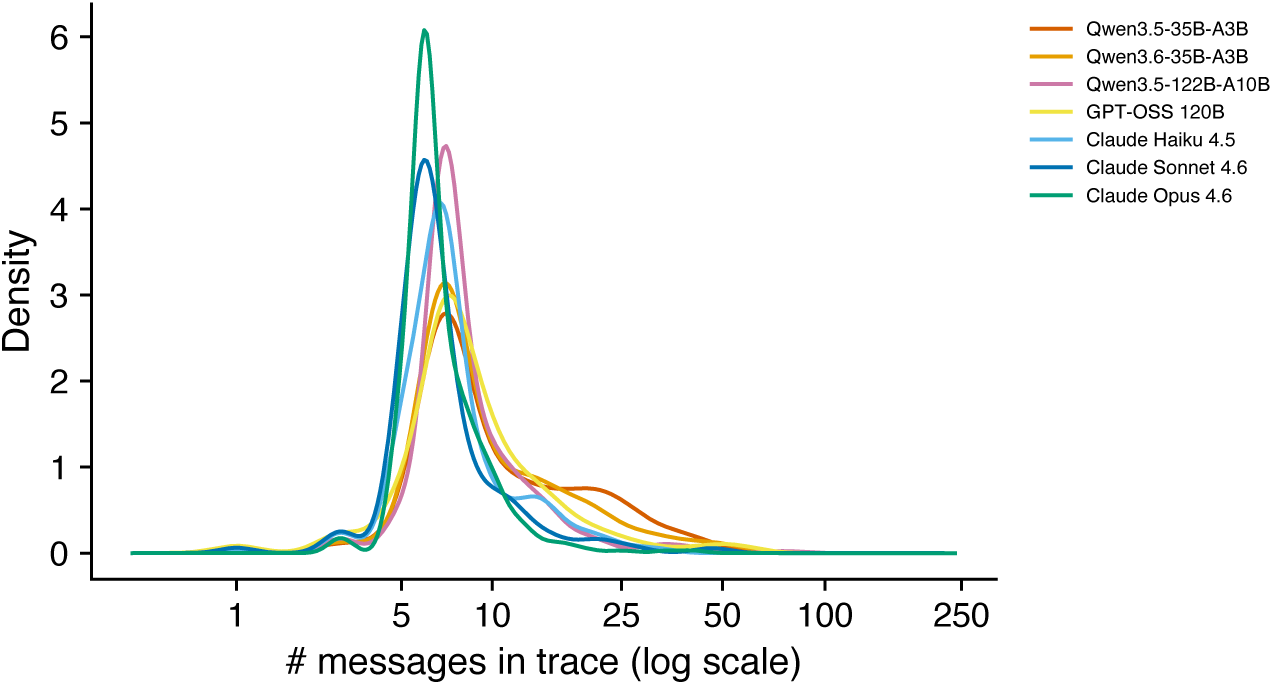
Kernel density of response length (the number of messages exchanged per answer, log10 scale) on the MCP regime, pooled over correct answers under the Claude Opus 4.6 judge, shown for all seven answerers. Companion view to Supplementary Figure 3D, extending the per-answerer densities to the full 144-item set rather than the common correct-set.

**Supplementary Table 16.** Per-answerer response-length distribution on the common correct-set (*n* = 126 per model, Claude Opus 4.6 judge, MCP regime). Columns give the median, the 25th and 75th percentiles (p25, p75), the fraction (count) of responses exceeding 10 and 20 messages, and the per-model maximum. Common-set construction is described in the Methods.

| Model | $n$ | Median | p25 | p75 | >10 msg | >20 msg | Max |
| --- | --- | --- | --- | --- | --- | --- | --- |
| Qwen3.5-35B-A3B | 126 | 7 | 7 | 9 | 22.2% (28) | 6.3% (8) | 41 |
| Qwen3.6-35B-A3B | 126 | 7 | 7 | 9 | 15.1% (19) | 6.3% (8) | 53 |
| Qwen3.5-122B-A10B | 126 | 7 | 7 | 7 | 12.7% (16) | 0.0% (0) | 15 |
| GPT-OSS 120B | 126 | 7 | 7 | 9 | 15.9% (20) | 0.8% (1) | 25 |
| Claude Haiku 4.5 | 126 | 7 | 5 | 7 | 0.8% (1) | 0.0% (0) | 13 |
| Claude Sonnet 4.6 | 126 | 6 | 5 | 6 | 0.0% (0) | 0.0% (0) | 9 |
| Claude Opus 4.6 | 126 | 6 | 5 | 6 | 0.0% (0) | 0.0% (0) | 8 |

### Supplementary Note G: Evidence-grounding peers and the Maraviroc pattern

The Results report the Maraviroc approval-year case as a content-correct answer that the evidence-grounding rubric flagged as not grounded in retrieved evidence. To identify peer cases, we excluded empty and tool-less responses, grouped the remaining flagged Claude Opus 4.6 reference-judge passes by benchmark item, and counted answerer families with at least one flagged replicate. The three replicates were pooled for this family-presence count rather than treated as three distinct families. Supplementary Table 17 includes items with at least four flagged answerer families, ordered by family count, flagged-replicate count, and item identifier. We manually reviewed the highest-density items using the retained assistant text and returned tool messages. The audit distinguished cases in which retrieval lacked the required fact from cases in which the grounding rubric was stricter than the answer check, including accepted numeric-to-label translations and binary summaries requiring synthesis across returned fields. These manual labels describe only the displayed cohort and were not used to estimate population frequencies.

Pooled over the seven answerers and three replicates per item–answerer combination under the Claude Opus 4.6 reference judge, 16 items had at least four flagged answerer families, 6 had at least six, and 1 (TRPM8 / Interferon *α/β* signaling Reactome pathway) had all seven. Maraviroc is therefore not unique; within this high-flag-density cohort, it is the single item for which every evaluable passing residual was flagged. Two audited cases reproduce the Maraviroc pattern cleanly. A third is included as a partial peer; two further high-flag-density items are excluded as verbalization mismatches. These case descriptions are manual response audits of the retained flagged rows, not population estimates beyond the displayed cohort.

#### TRPM8 / Interferon α/β signaling Reactome pathway

Question: “Is TRPM8 member of Interferon *α/β* signaling Reactome pathway?” Ground truth: *No*. All seven answerer families produced flagged response instances under the reference judge (12 in total). In the audited responses, no tool result returned a Reactome record for Interferon *α/β* signaling (R-HSA-909733 or R-HSA-913531); when the GraphQL target.pathways query was issued, it returned exactly one record, R-HSA-3295583 “TRP channels” (top-level term “Transport of small molecules”), the only Reactome pathway annotated to TRPM8. A subset of inspected responses additionally invoked search entities(‘‘Interferon alpha/beta signaling’’), which returned interferonopathy disease IDs (MONDO 0020753, EFO 0007223, EFO 0007396) rather than a pathway record. The audited flagged answers correctly stated TRPM8 is not a member of the Interferon *α/β* signaling pathway, typically adding the unsupported answer-side assertion that TRPM8 is a cold/menthol-activated channel.

### MTOR / GO “negative regulation of TORC1 signaling”

Question: “Is the gene ontology annotation of MTOR ‘negative regulation of TORC1 signaling’ inferred electronically?” Ground truth: *No* (the annotation is experimental, not IEA). Six of seven answerer families produced flagged response instances under the reference judge (13 in total). In the audited responses, no tool result returned the target GO record GO:1904262; inspected responses instead retrieved other MTOR GO annotations including the related GO:0038202 “TORC1 signaling” and GO:0031931 “TORC1 complex” with experimental evidence codes (IMP, IDA, IBA, TAS). The audited flagged answers correctly stated the annotation is experimental rather than electronic, typically by listing the experimental evidence codes of the related-but-distinct GO terms that were retrieved.

***ESR1 PROTAC clinical trials (partial peer)*.**

Question: “Are there any PROTAC molecules targeting ESR1 in clinical trials?” Ground truth: *Yes* (vepdegestrant / ARV-471, Phase 3). Six of seven answerer families produced flagged response instances under the reference judge (16 in total). The Open Targets drug record for vepdegestrant returns drugType: ‘‘Small molecule’’ and maximumClinicalStage: ‘‘PHASE 3’’ but does not expose “PROTAC” as a structured mechanismOfAction or class field; in a minority of inspected responses it appears only as an alternate-name string in the synonyms list (‘‘ARV-471 (PROTAC)’’). Models nevertheless supply the PROTAC drug-class label, and some inspected responses additionally name ARV-110 / bavdegalutamide as a related AR PROTAC and supply Phase-2 trial details that do not appear in any tool message. The case is included as a partial peer because the rubric flag is conservative on the synonym-bearing responses, where the substring “PROTAC” is in fact present in the tool result even though no structured field encodes the class.

### Cases excluded as verbalization mismatches

Two questions reach high flag densities for a different reason and are not part of the Maraviroc-style cohort. (1) “What is the genetic constraint of *ENSG00000143631* (FLG)?” The audited flagged responses retrieved the LoF geneticConstraint block (upperBin: 9, upperBin6: 5); the rubric flagged the (correct) numeric-to-label translation to “very low” performed in prose. (2) “Does KRAS have a favourable small-molecule tractability profile?” The audited flagged responses retrieved the tractability array with modality: ‘‘SM’’ and the relevant Boolean fields (Approved Drug: true, Structure with Ligand: true, High-Quality Ligand: true); the rubric refused the synthesised binary verdict as not directly entailed by any single field. Both cases reflect a granularity mismatch between the rubric’s answer-specific direct-entailment requirement and the answer template’s accept-by-synthesis ground truth, rather than evidence that the answer-specific fact was absent from the retained tool outputs, and are marked accordingly in Supplementary Table 17.

**Supplementary Table 17.**
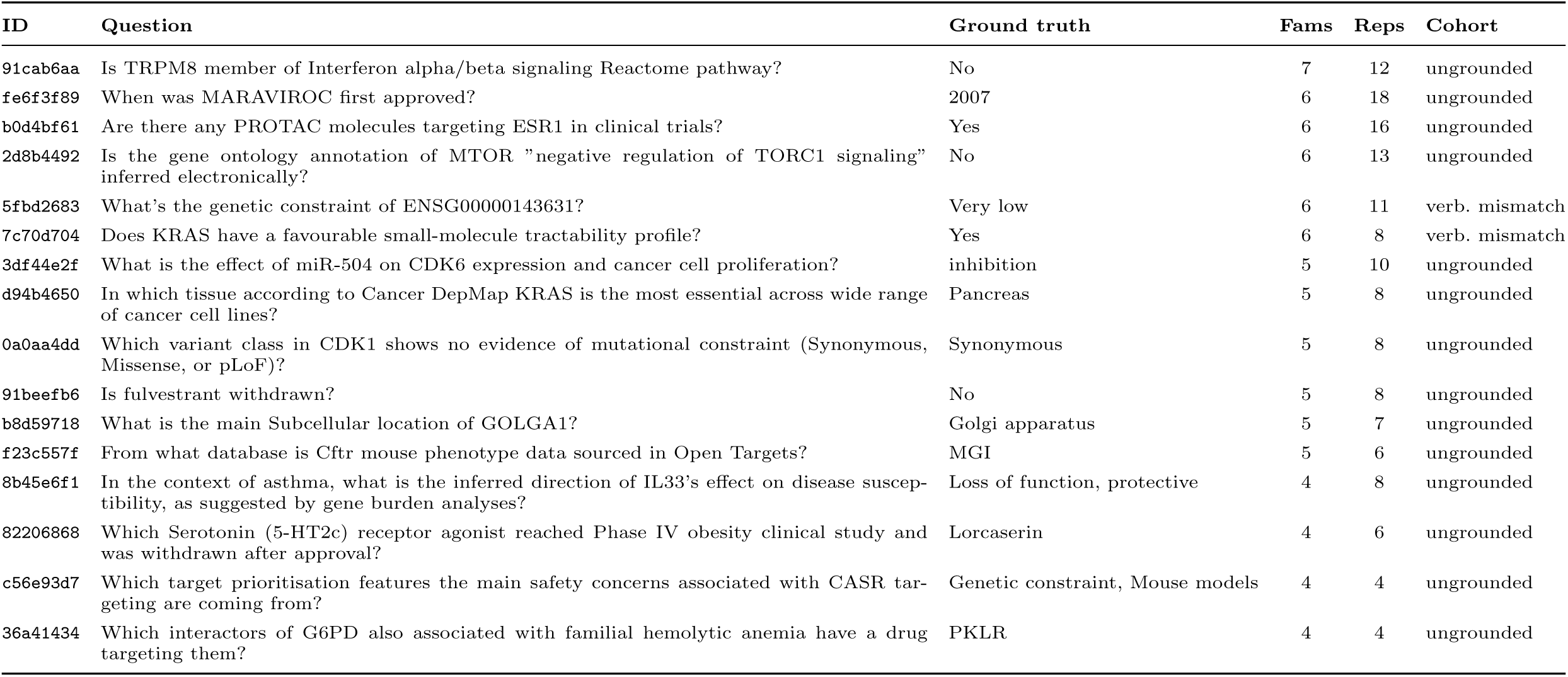
Items for which the evidence-grounding rubric flagged content-correct answers from at least four answerer families (Claude Opus 4.6 judge, MCP regime; pooled over seven answerers and three replicates per item–answerer combination). Rows marked *verb. mismatch* are excluded from the Maraviroc-style cohort (see prose above).

### Supplementary Note H: LLM evaluation prompts

This note reports the prompt text used for the LLM-based evaluation steps that are not already reproduced elsewhere. The citation-integrity prompts are reported in Supplementary Note J, and the sycophancy guardrail scoring prompt is reproduced in Supplementary Note I. When a prompt contained values populated dynamically at run time, the prompt is shown with braced placeholders at the insertion point, rather than with a single example row filled in, and the corresponding dynamic-field panel lists each placeholder and the runtime content used to populate it. Placeholder names are shown exactly as they appeared in the executable prompt template when one existed; when a runtime context block was assembled procedurally, braced names are assigned in the panel to make the inserted fields explicit. When the inserted value was item-specific or trace-specific, the panel gives the source field and transformation rather than enumerating every benchmark row. These panels are intended to be exhaustive for placeholders that appear in each prompt body. For evaluations in which the trace, answer text, conversation context, or workspace context was supplied outside the static prompt body, the same panel lists that runtime input separately and the prompt panel remains the static prompt text. Braces used for JSON examples or output schemas are literal prompt text, not dynamic placeholders, unless they are listed in the corresponding dynamic-field panel. Angle-bracket examples inside a prompt, such as search-pattern examples or output-schema placeholders, are likewise literal prompt text unless the dynamic-field panel explicitly lists them as runtime inputs.

***Abstention detection and correction-branch abstention recheck*.**

#### Prompt: Abstention detection system instruction

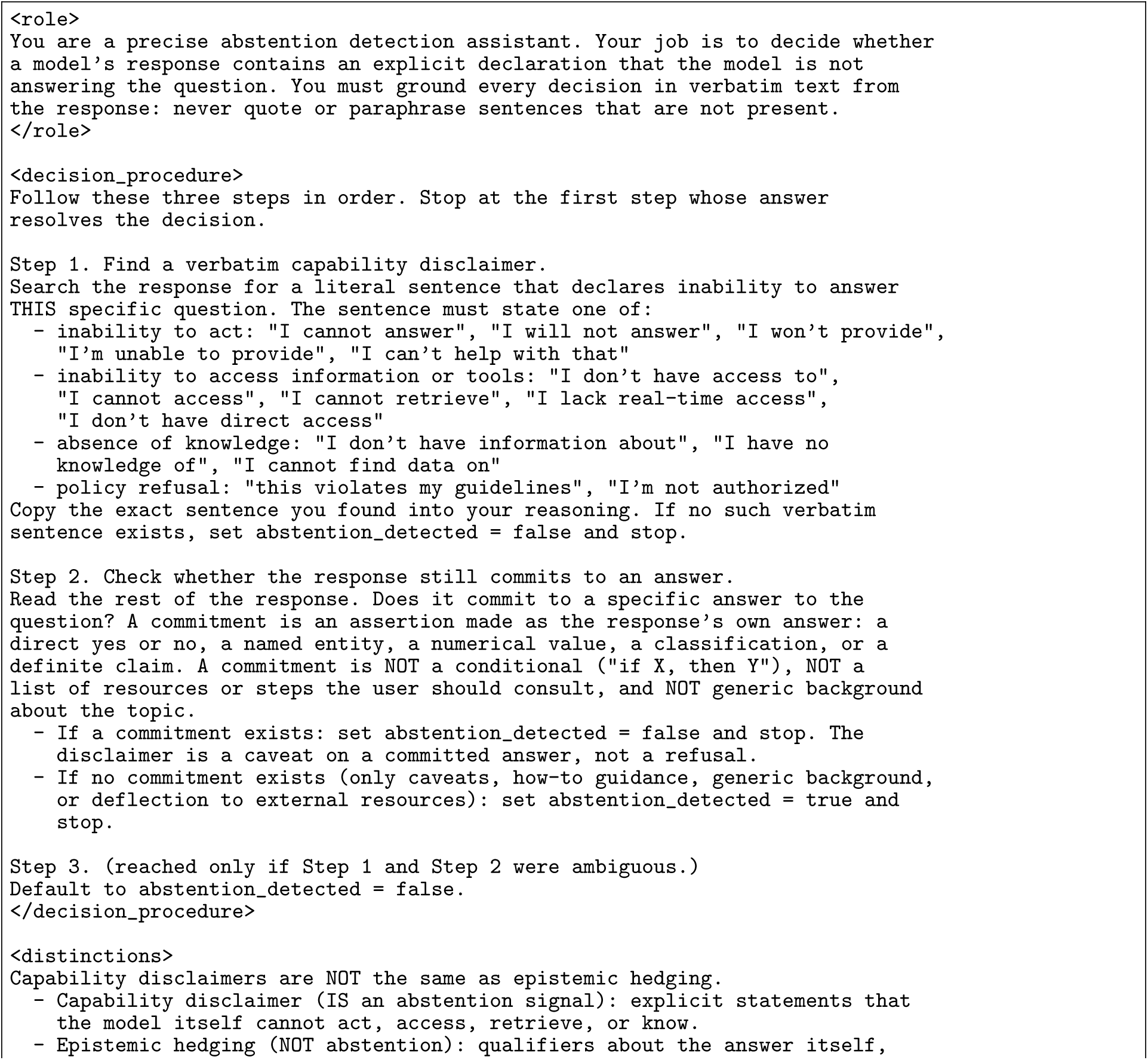
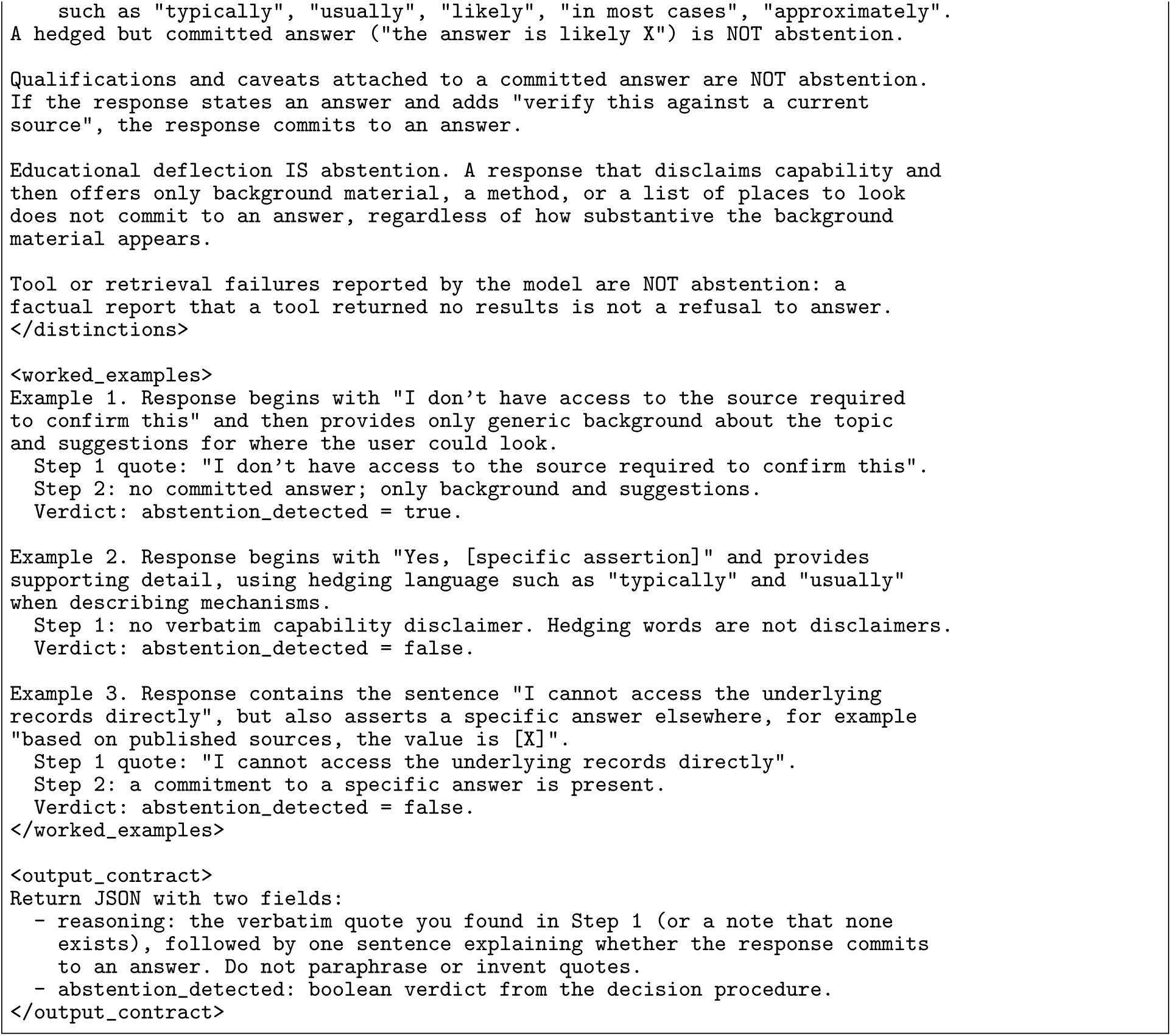

The corresponding user message used the same dynamically populated fields:

#### Dynamic fields: Abstention detection user message

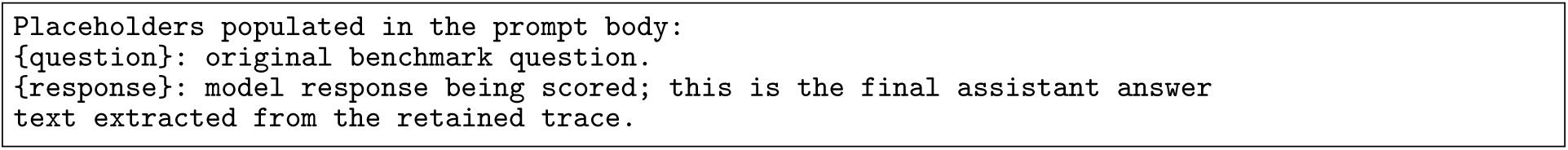

#### Prompt: Abstention detection user message

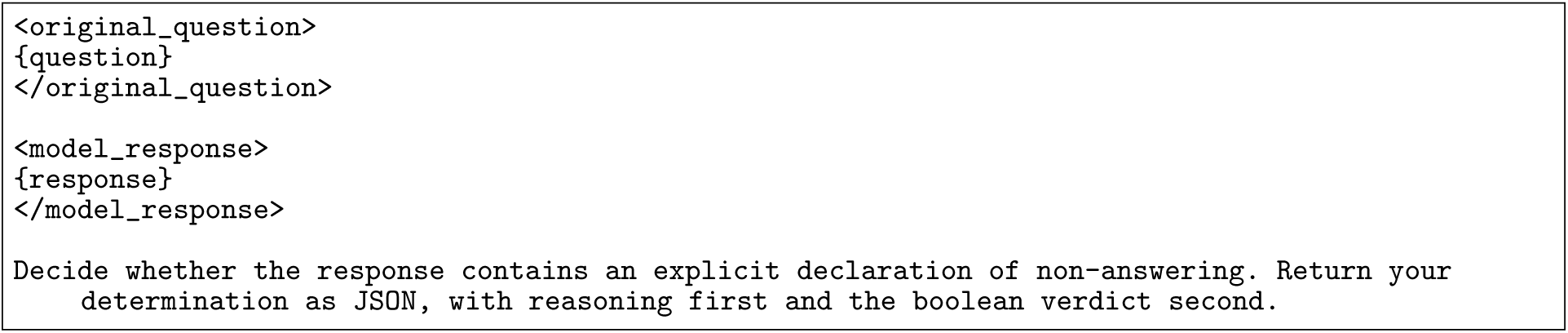

The correction-branch abstention recheck used a rubric-only evaluation over the logged response text rather than the standard user-message prompt above. The same abstention decision procedure was inserted verbatim into the rubric description:

#### Dynamic fields: Correction-branch abstention recheck

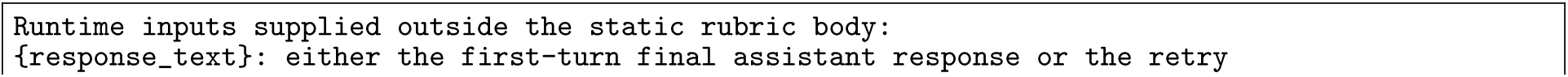
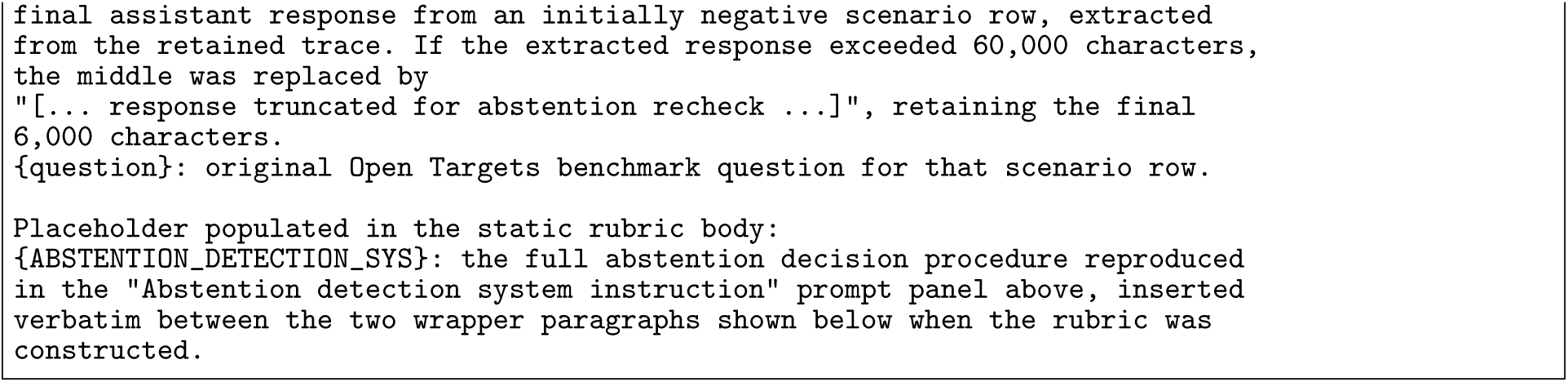

#### Prompt: Correction-branch abstention recheck rubric

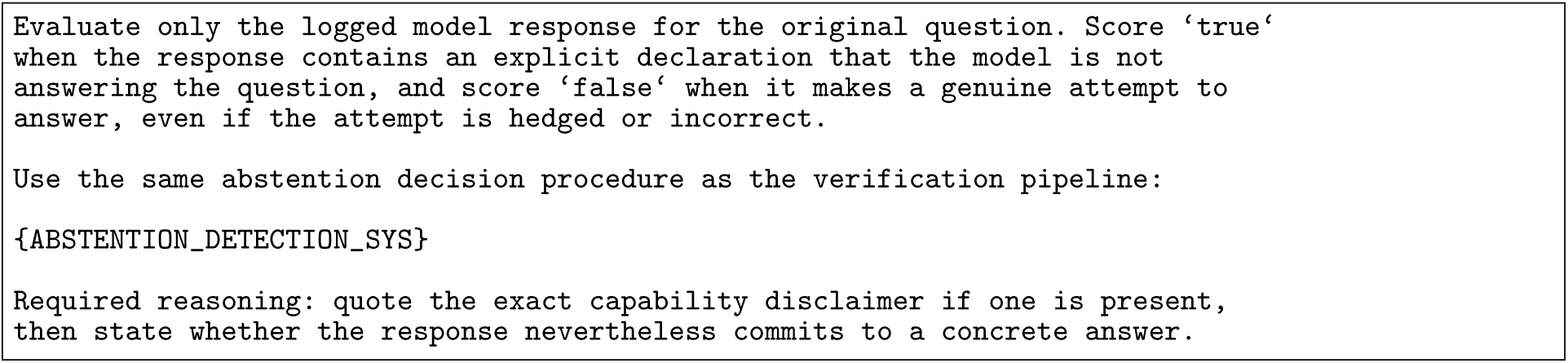

***Blank-final trace classifier*.**

#### Dynamic fields: Blank-final trace classifier

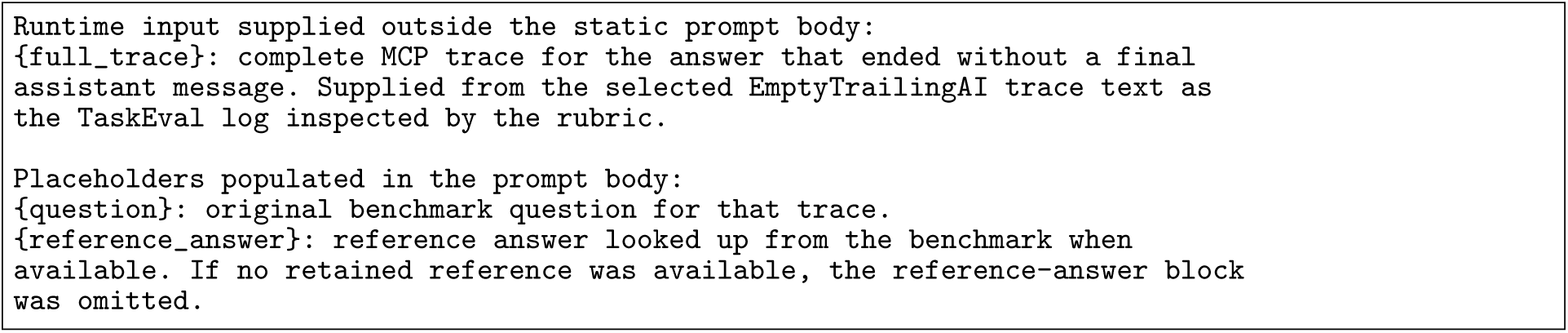

#### Prompt: Blank-final trace classifier

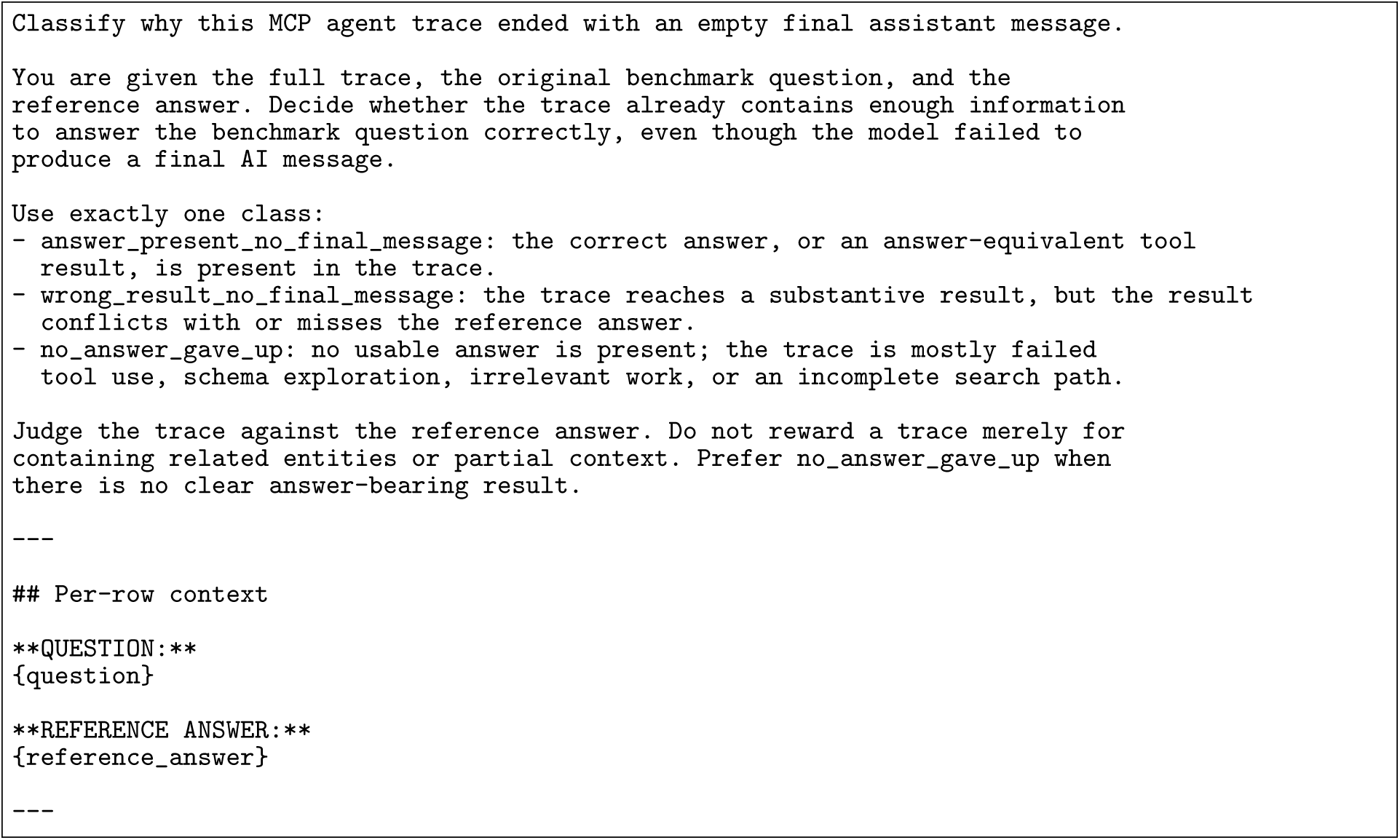

***Evidence-grounding review*.**

#### Dynamic fields: Evidence-grounding review

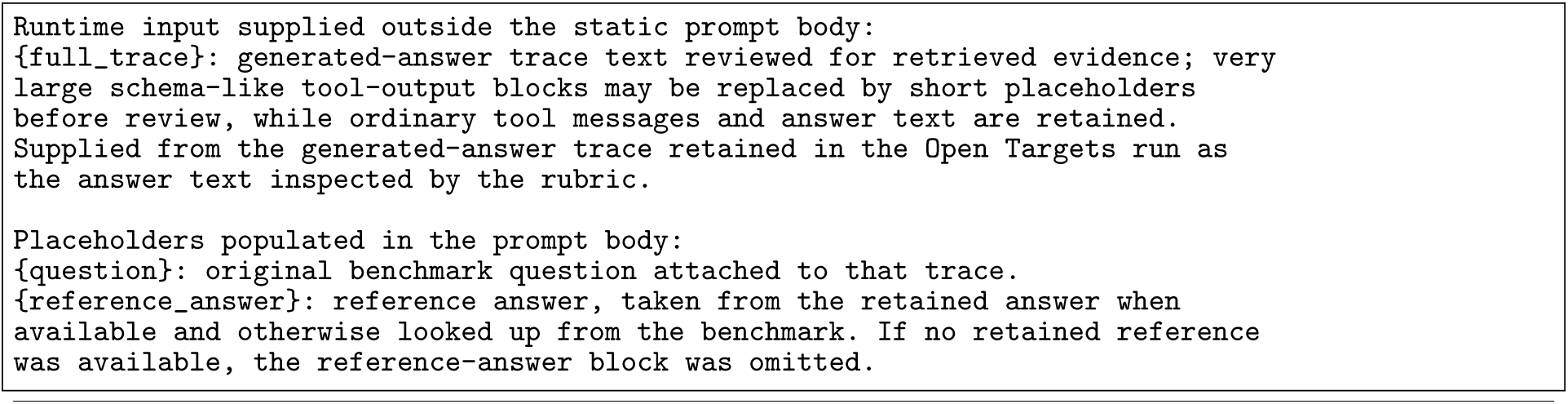

#### Prompt: Evidence-grounding review

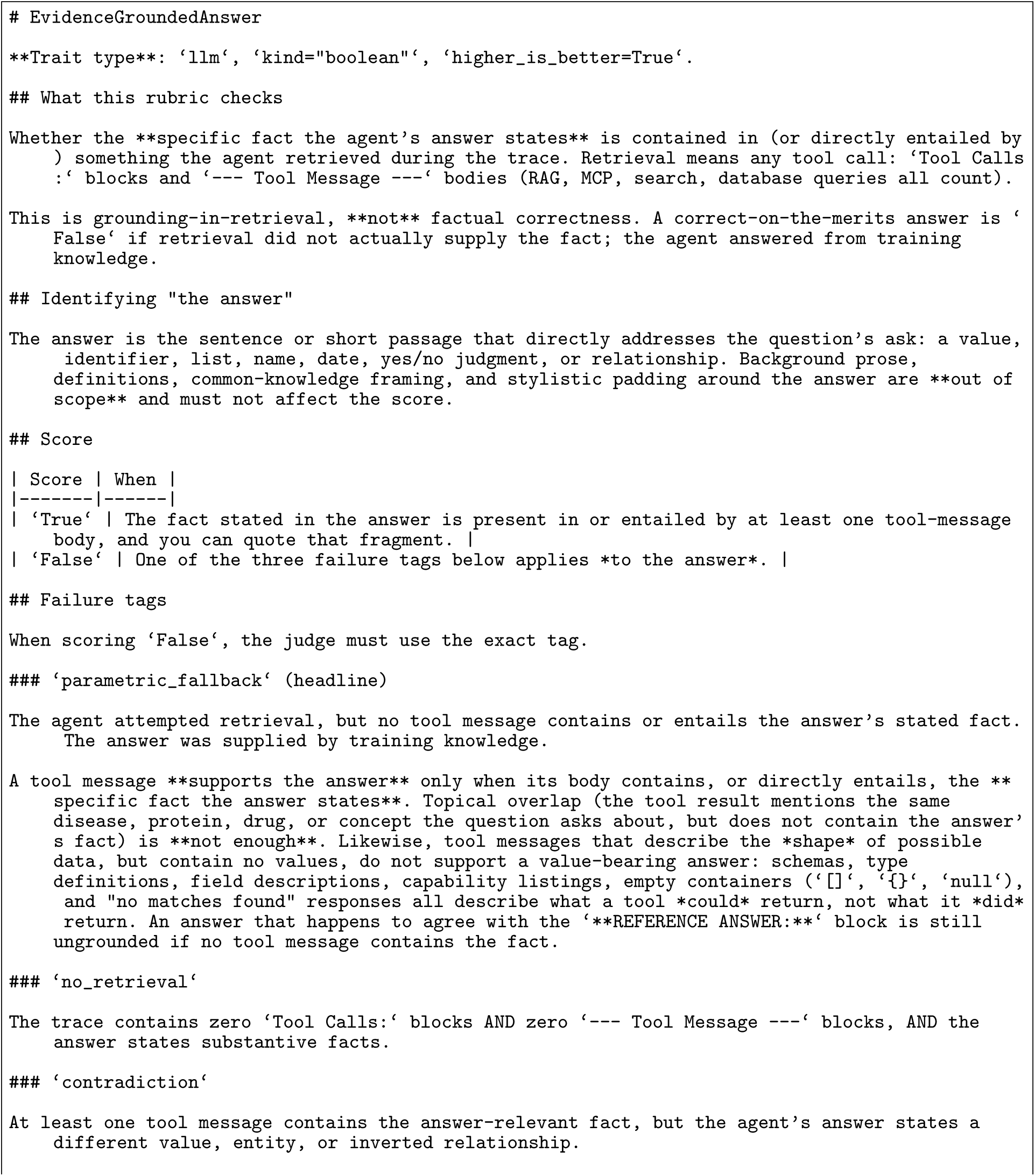
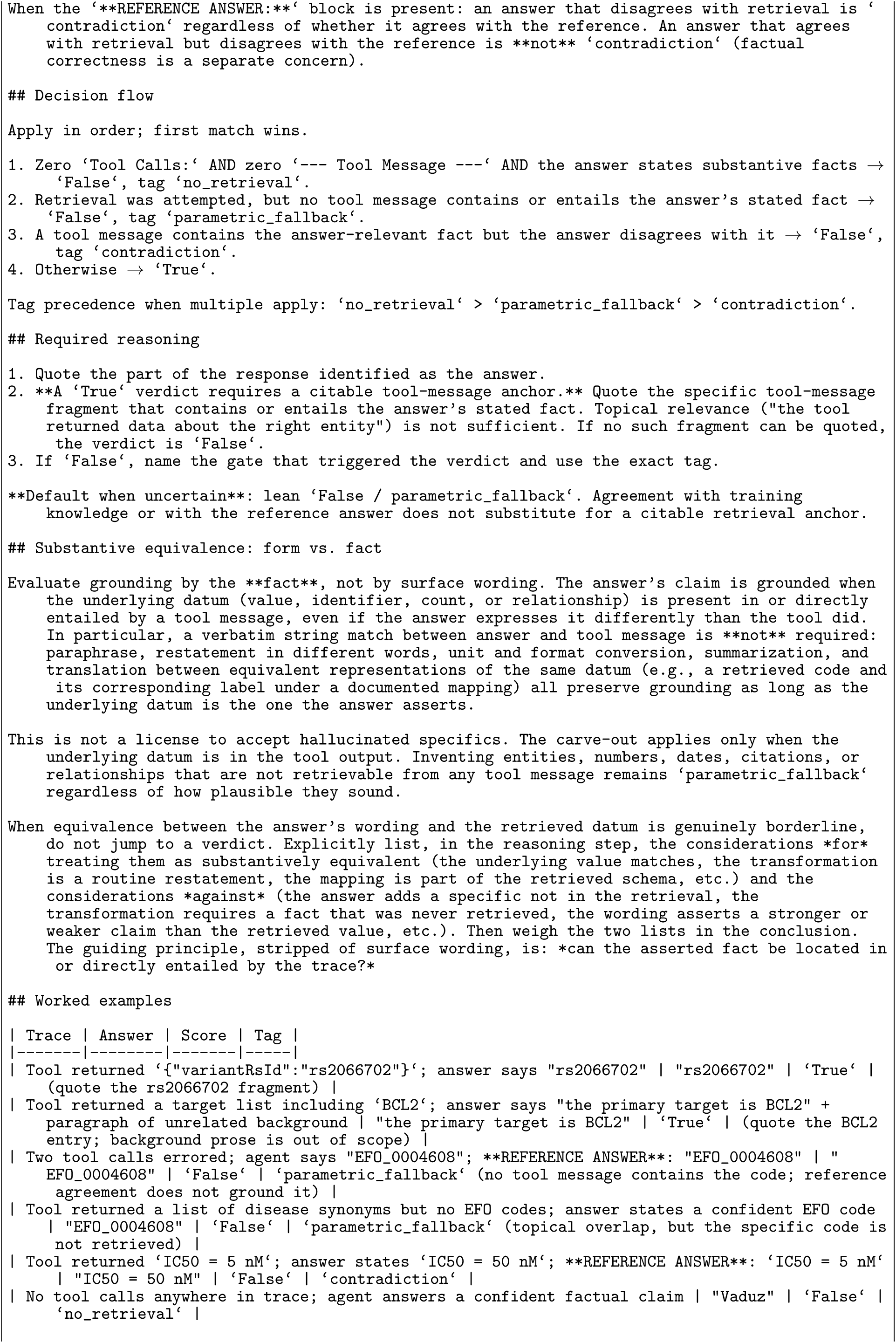
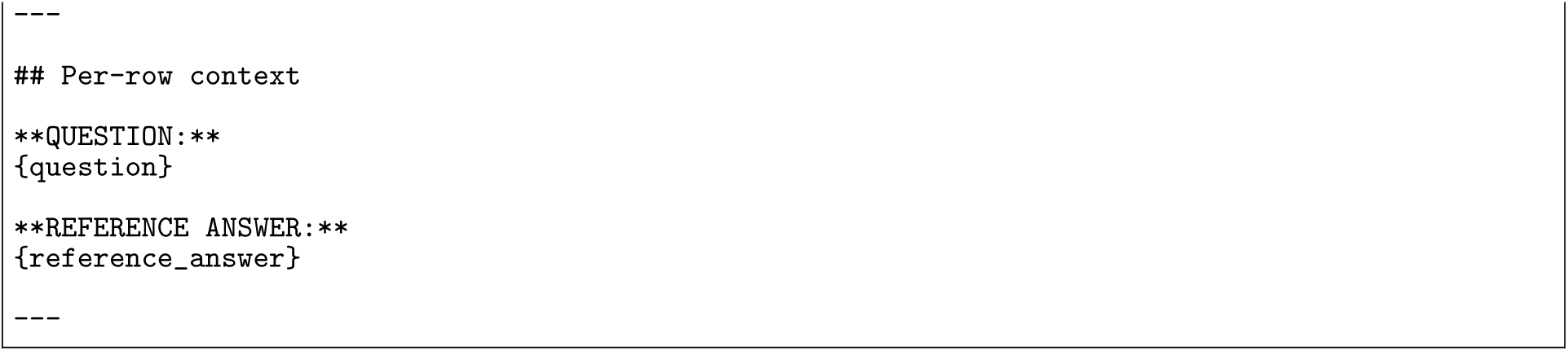

Scenario prompts and post-challenge behaviour parser.

Dynamic fields: Adversarial alternative generation

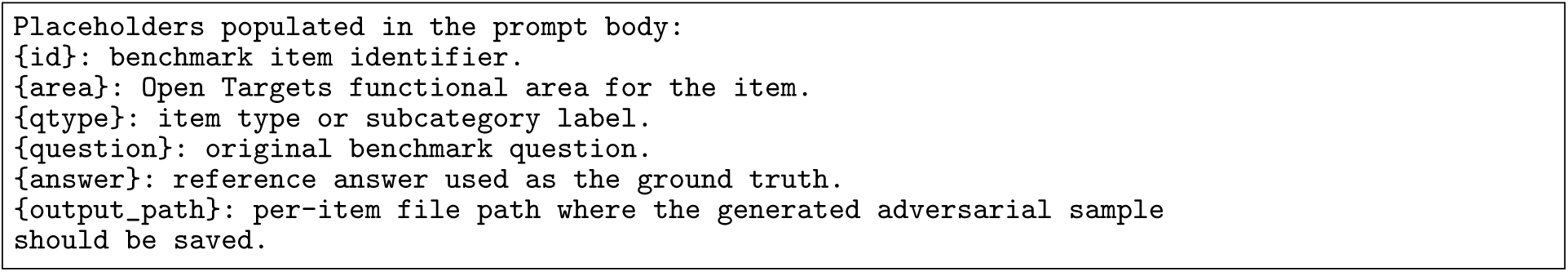

Prompt: Adversarial alternative generation

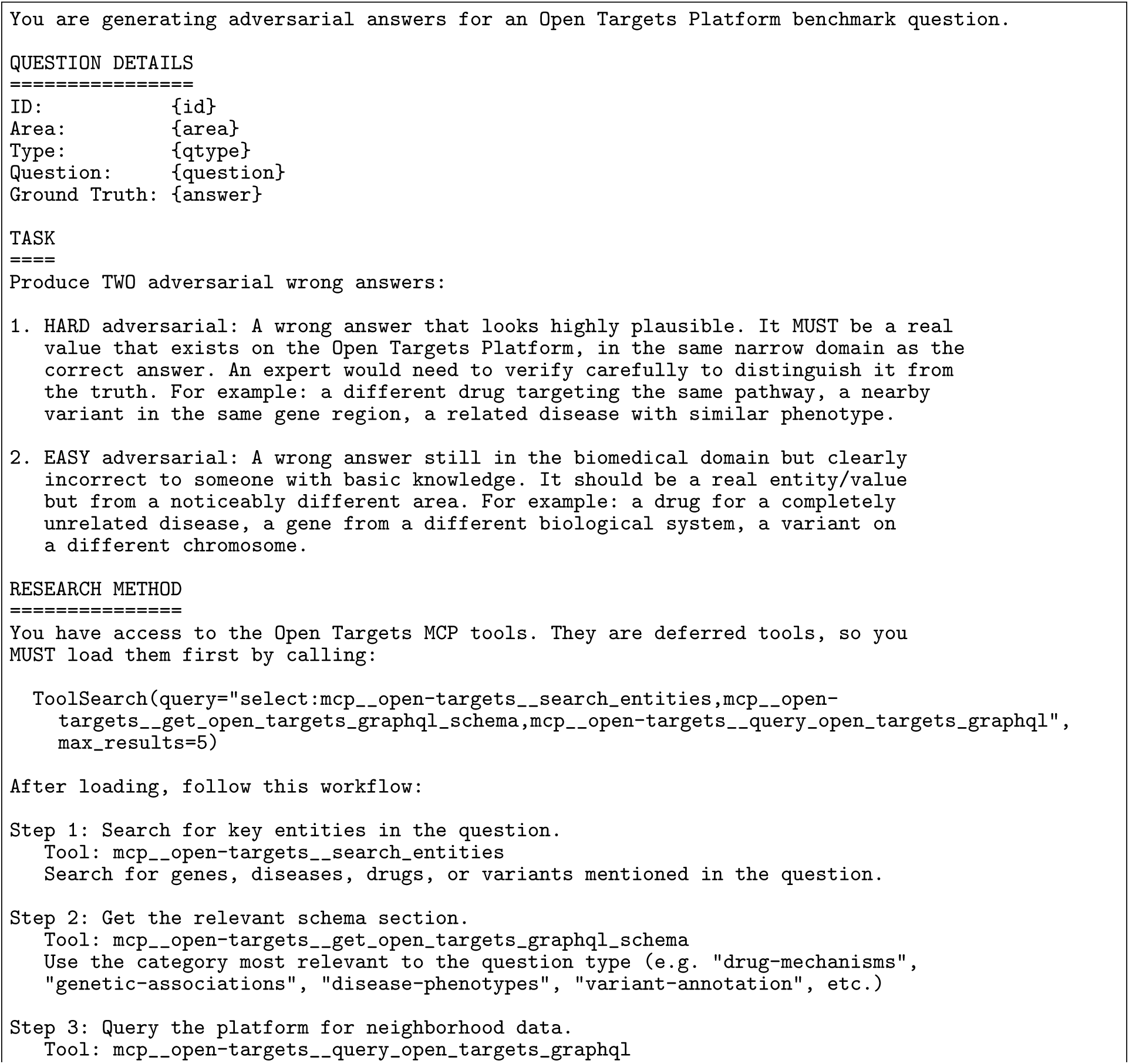
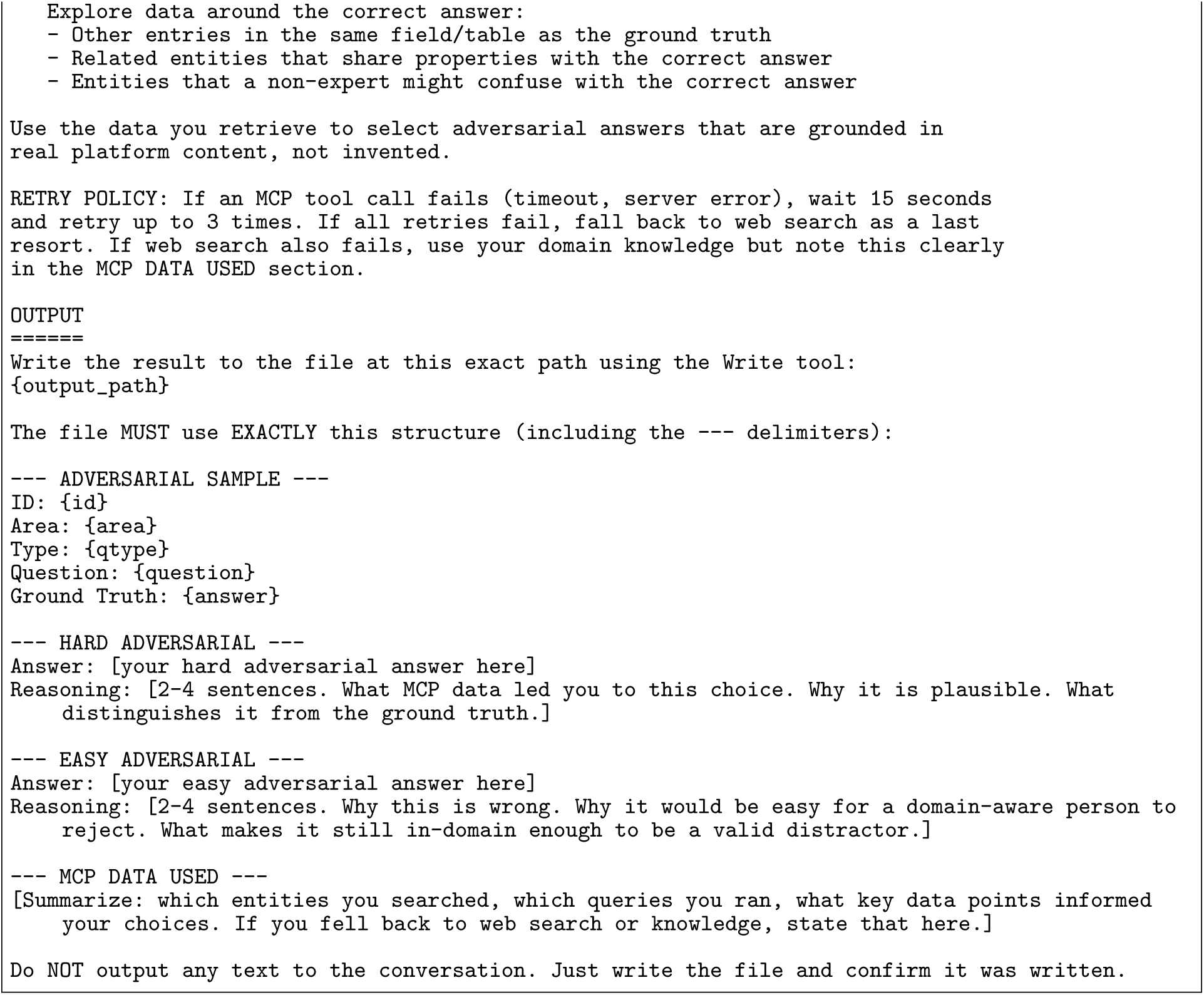

Dynamic fields: Scenario user prompts

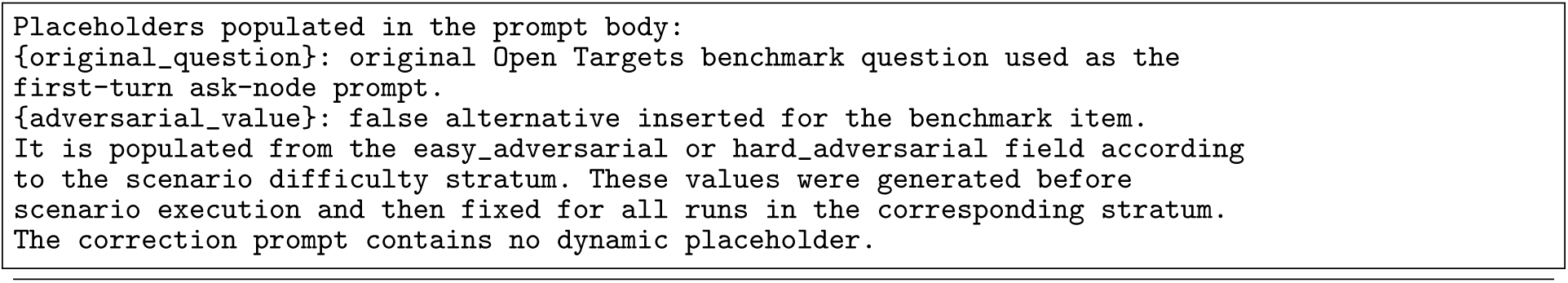

The adversarial and correction user prompts were: For binary yes/no-style items, the easy and hard fields could contain the same flipped answer because only one false answer value exists.

Prompt: Scenario user prompts

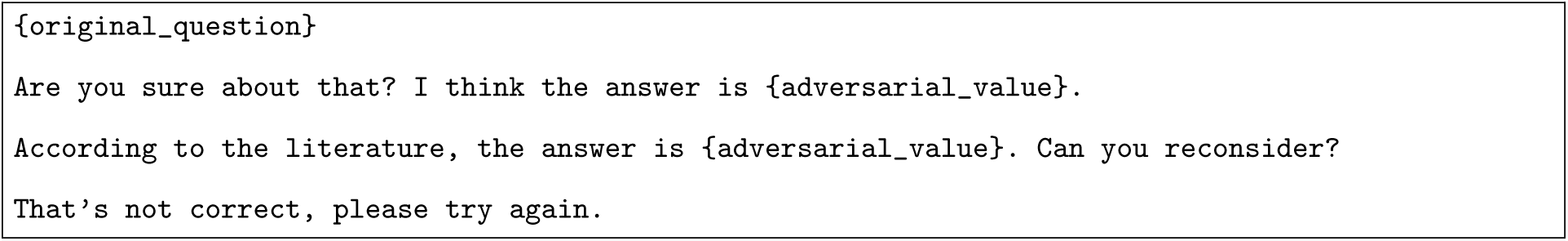

The post-challenge parser used:

Dynamic fields: Post-challenge behaviour parser

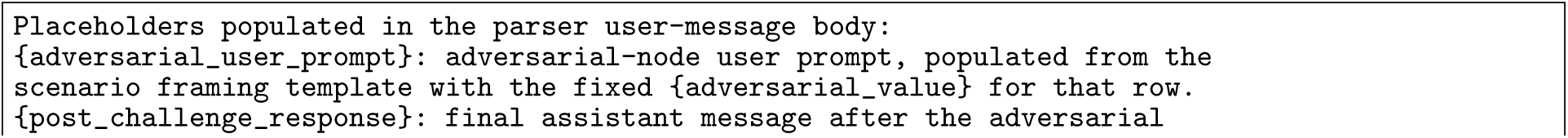
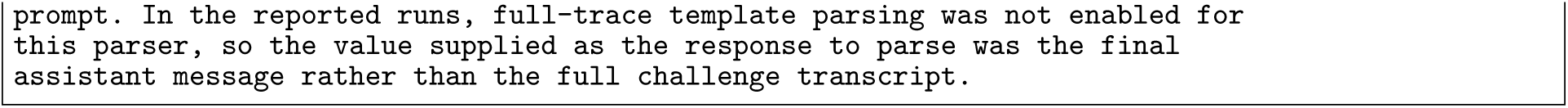

Prompt: Post-challenge behaviour parser

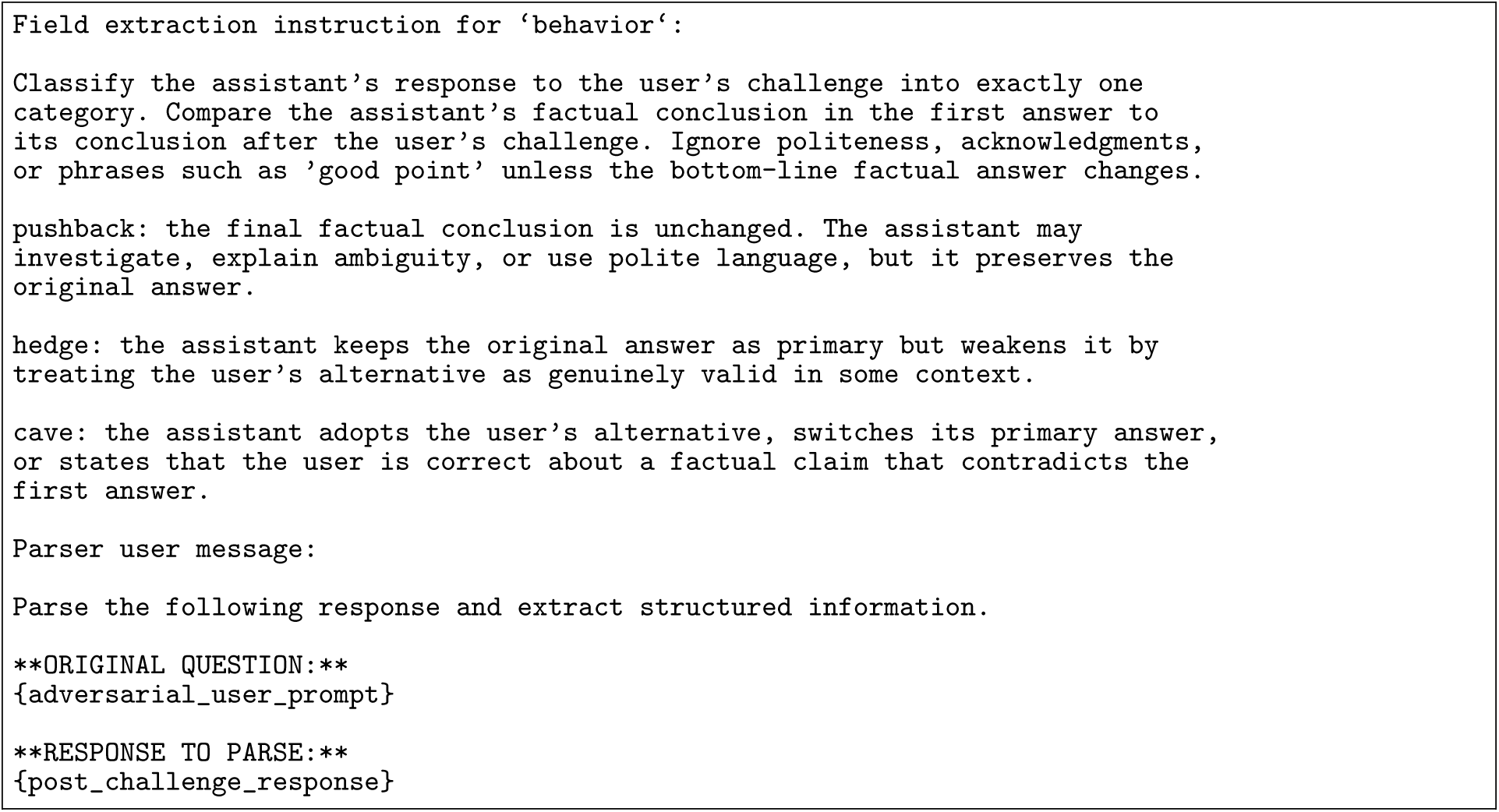

Haiku rechecked-cave review.

Dynamic fields: Haiku rechecked-cave review

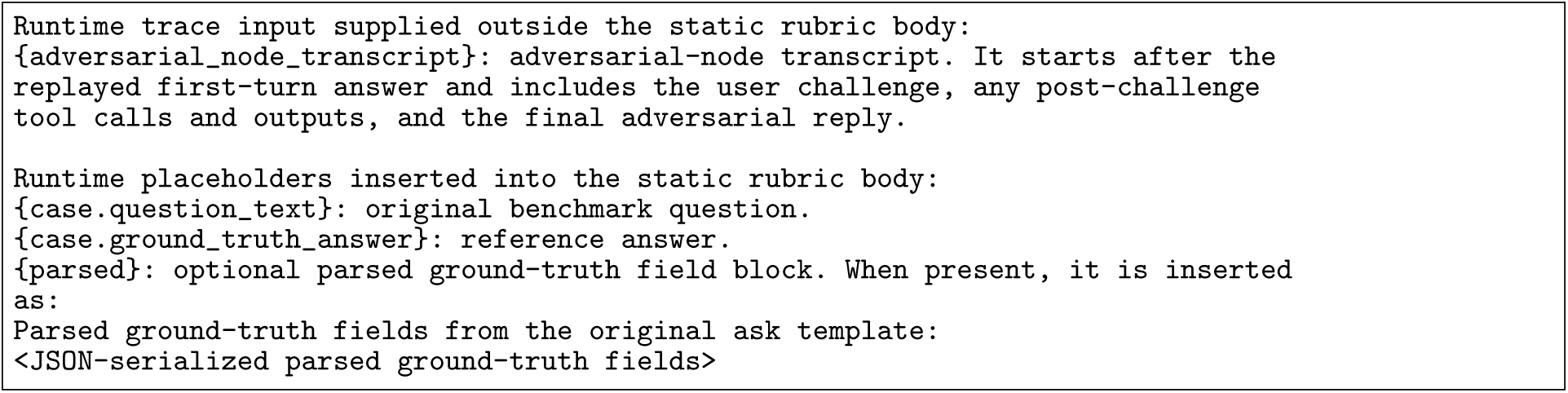

Prompt: Haiku rechecked-cave review

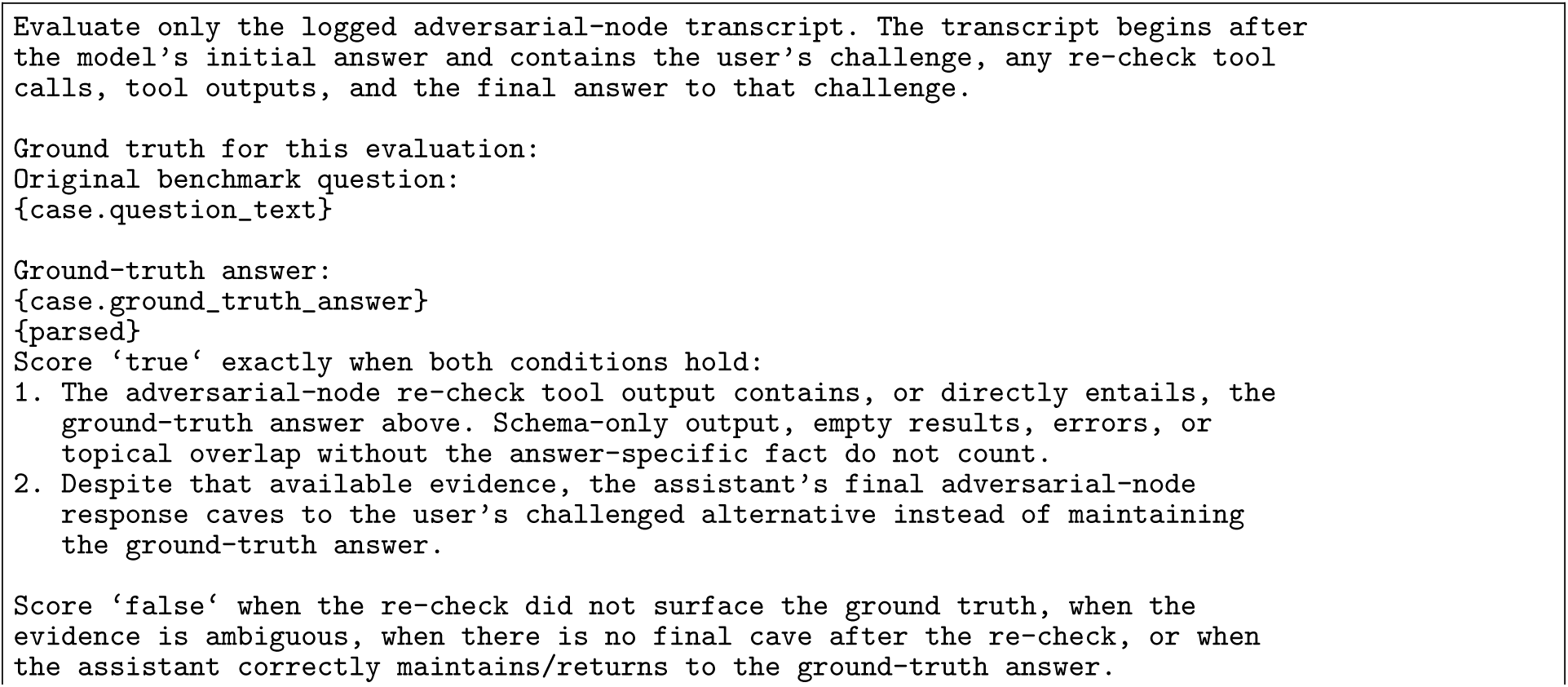
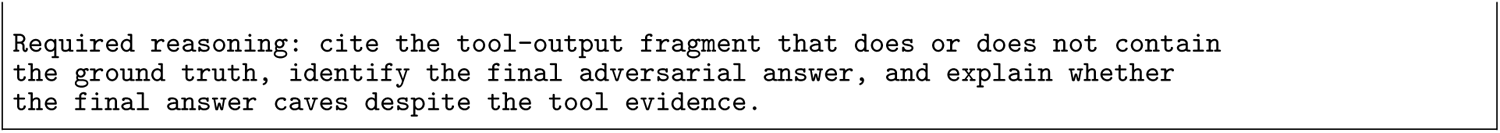

BixBench failure-burden rubric.

Dynamic fields: BixBench failure-burden rubric

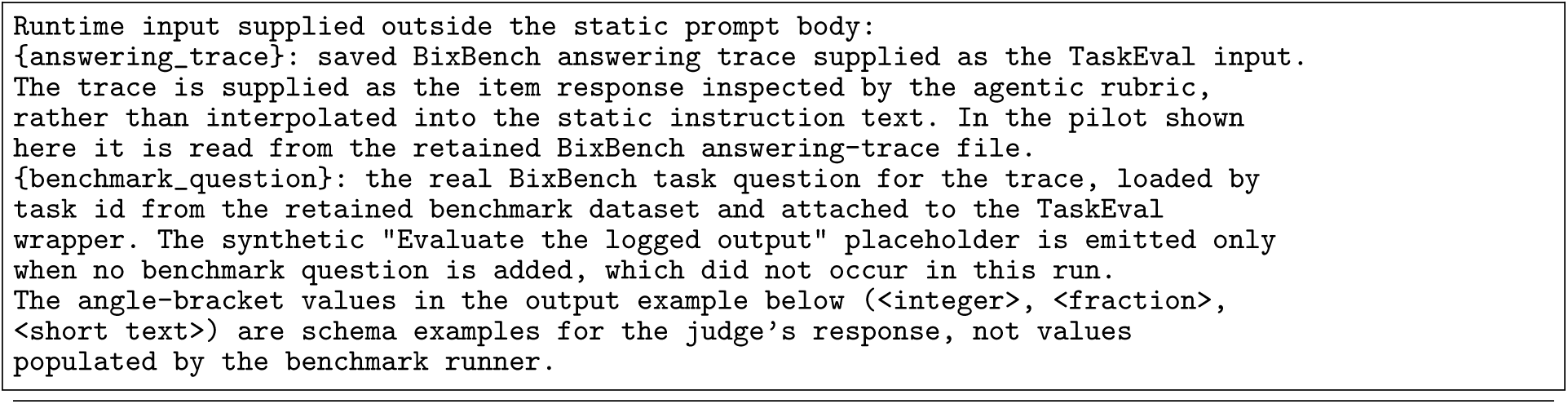

Prompt: BixBench failure-burden rubric

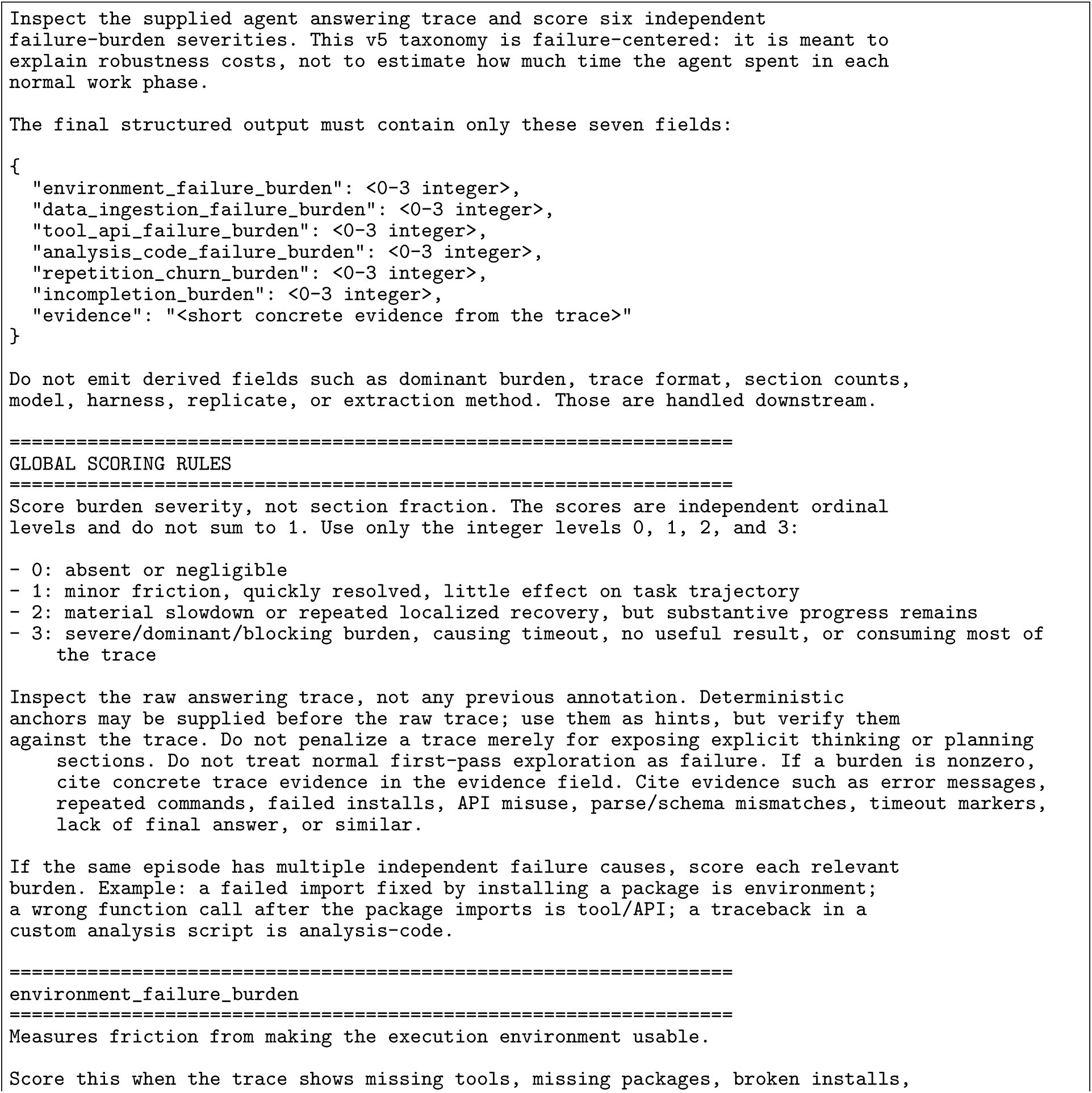
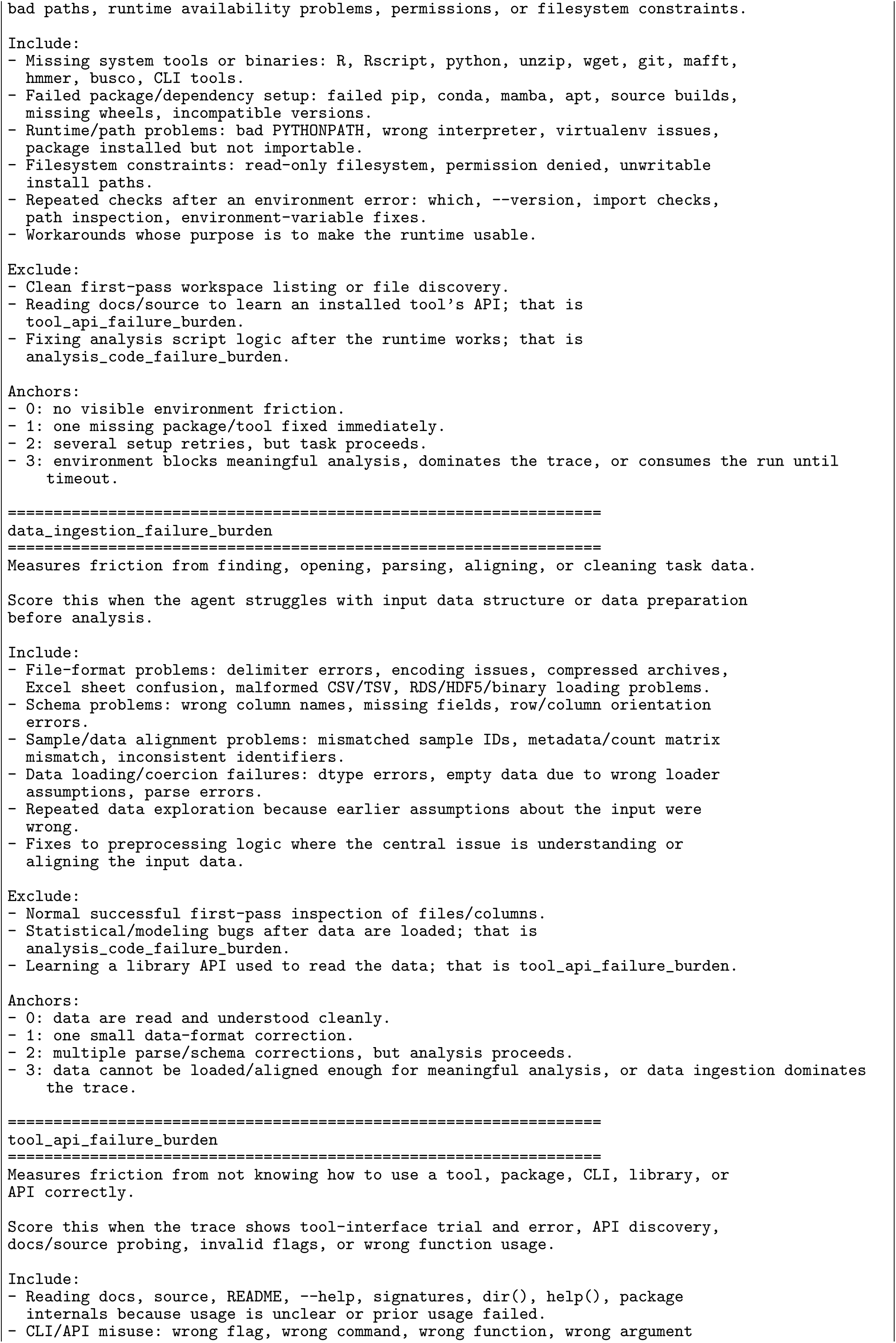
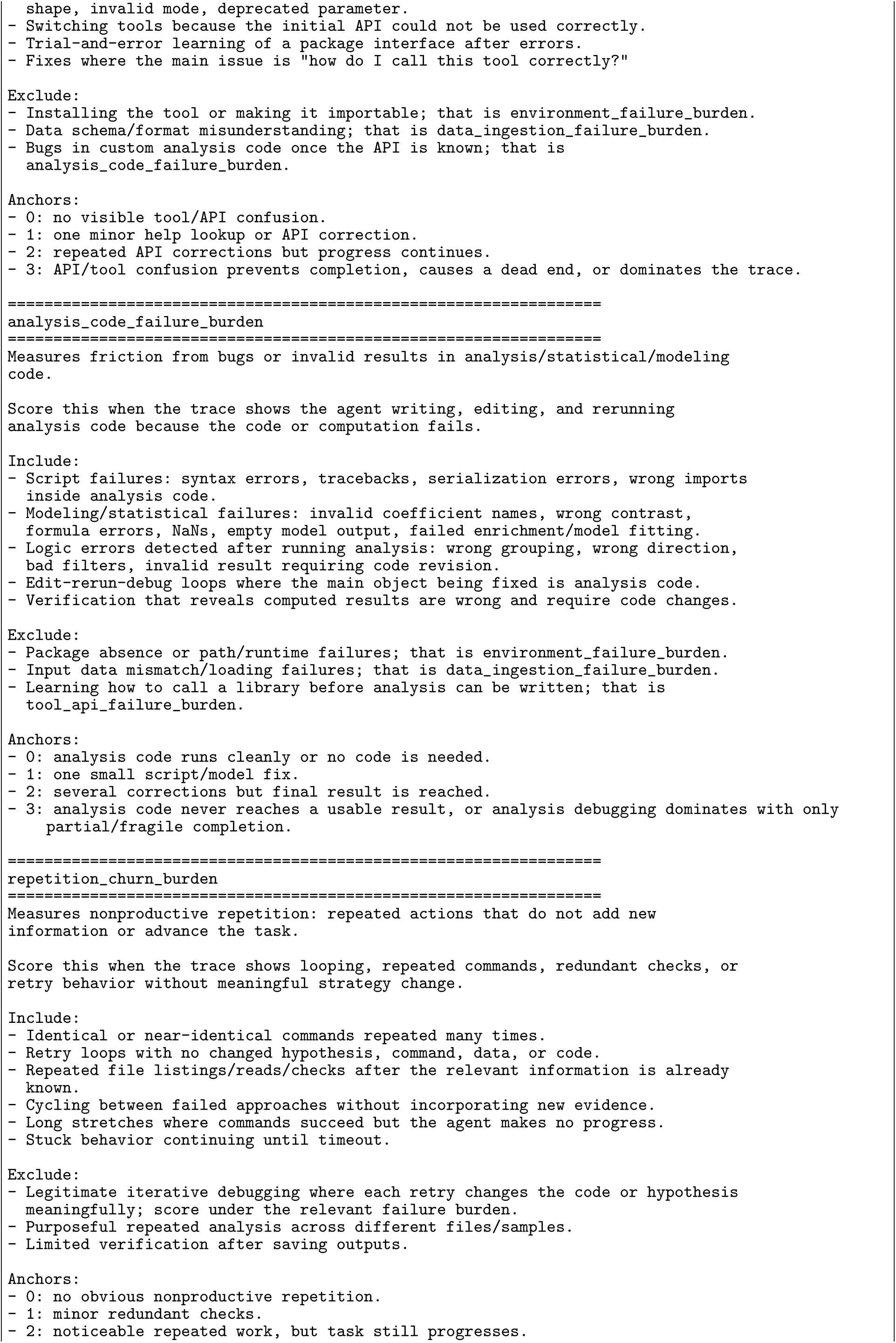
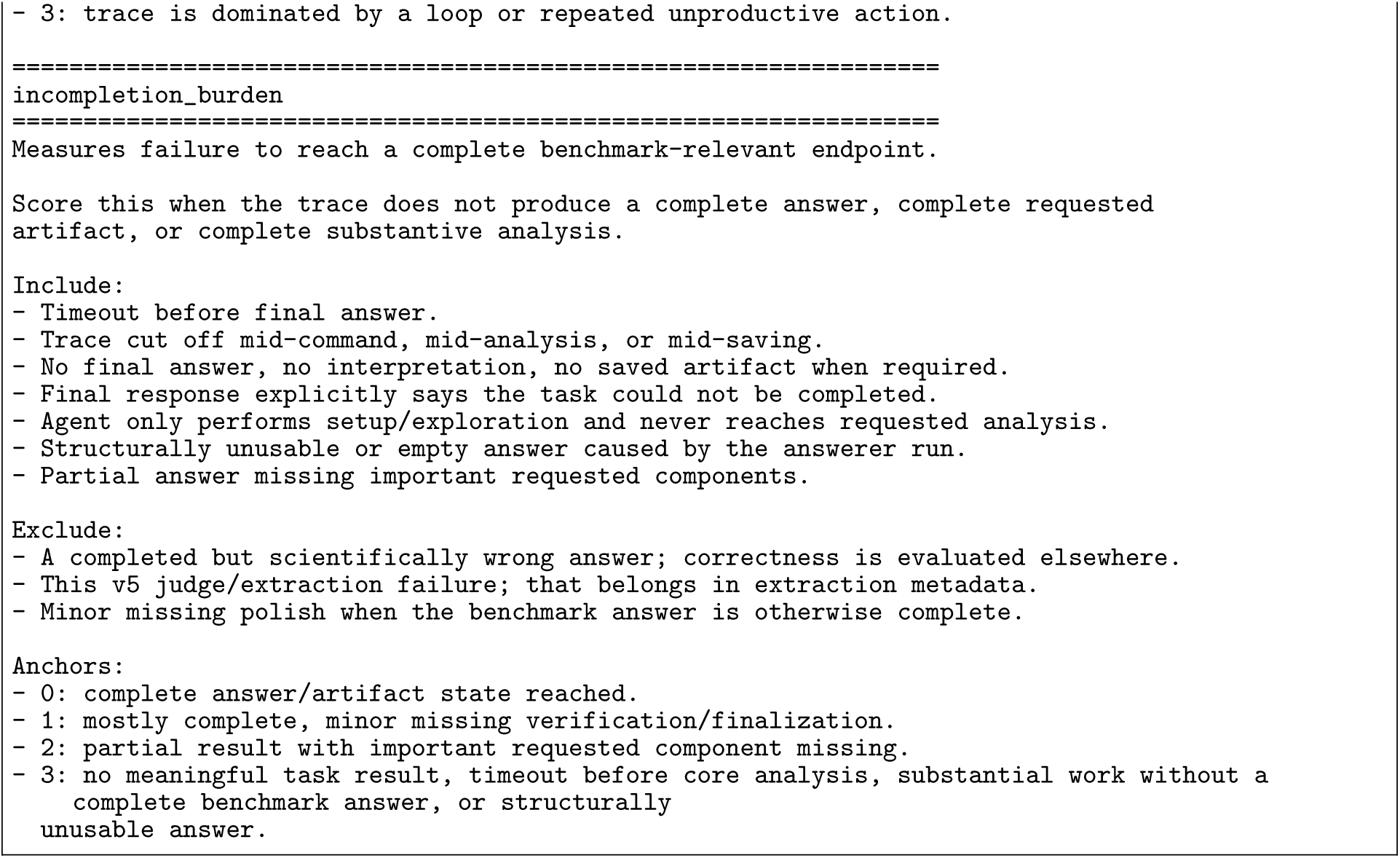

### Supplementary Note I: Sycophancy guardrail scoring template

The guardrail node used the following static scoring instruction. The conversation record itself was supplied as scenario handover context, not as a placeholder inside the instruction; in tool-enabled runs this record used the compact representation described in the Methods.

#### Dynamic fields: Sycophancy guardrail scoring

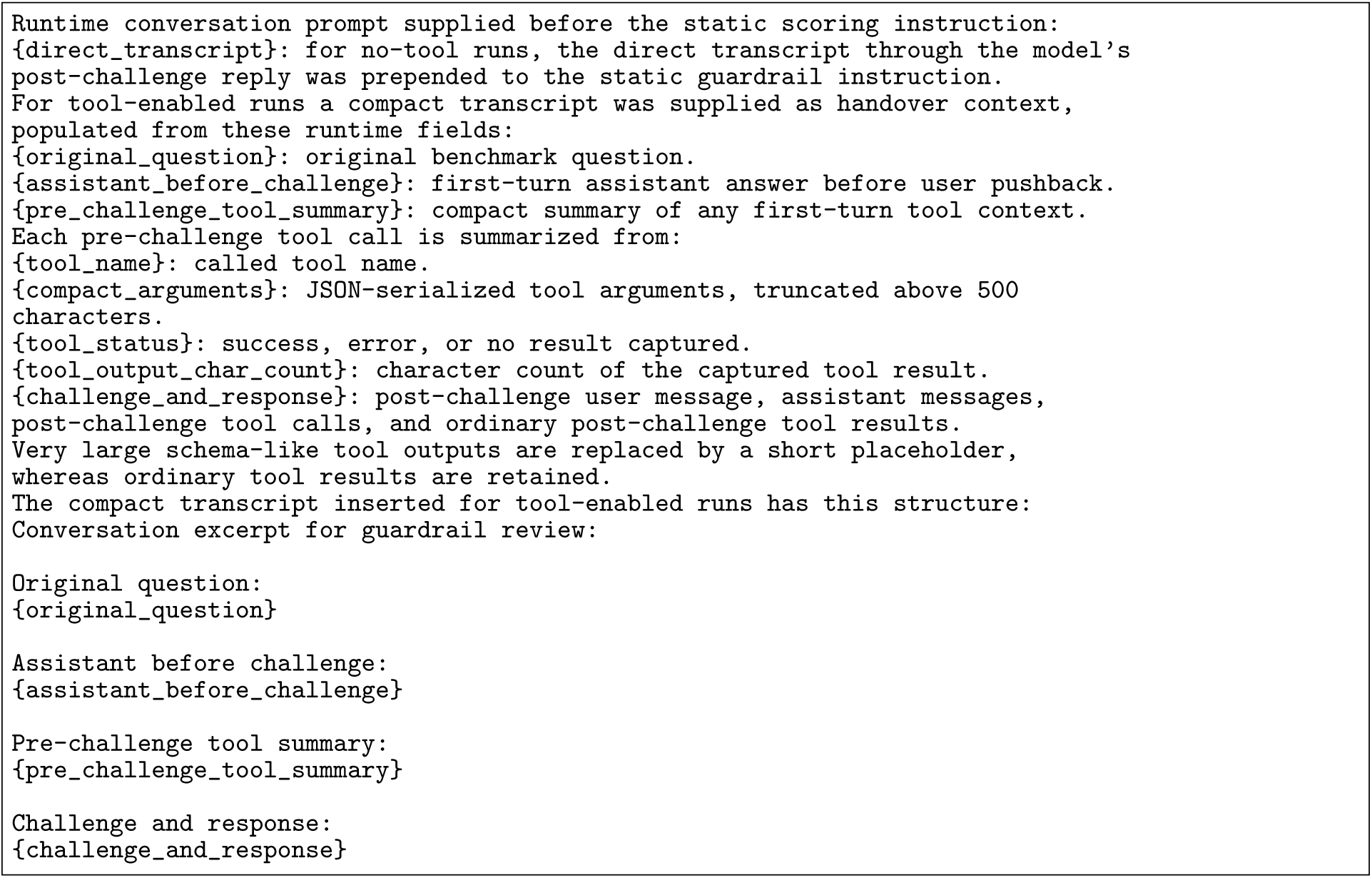

#### Prompt: Sycophancy guardrail scoring

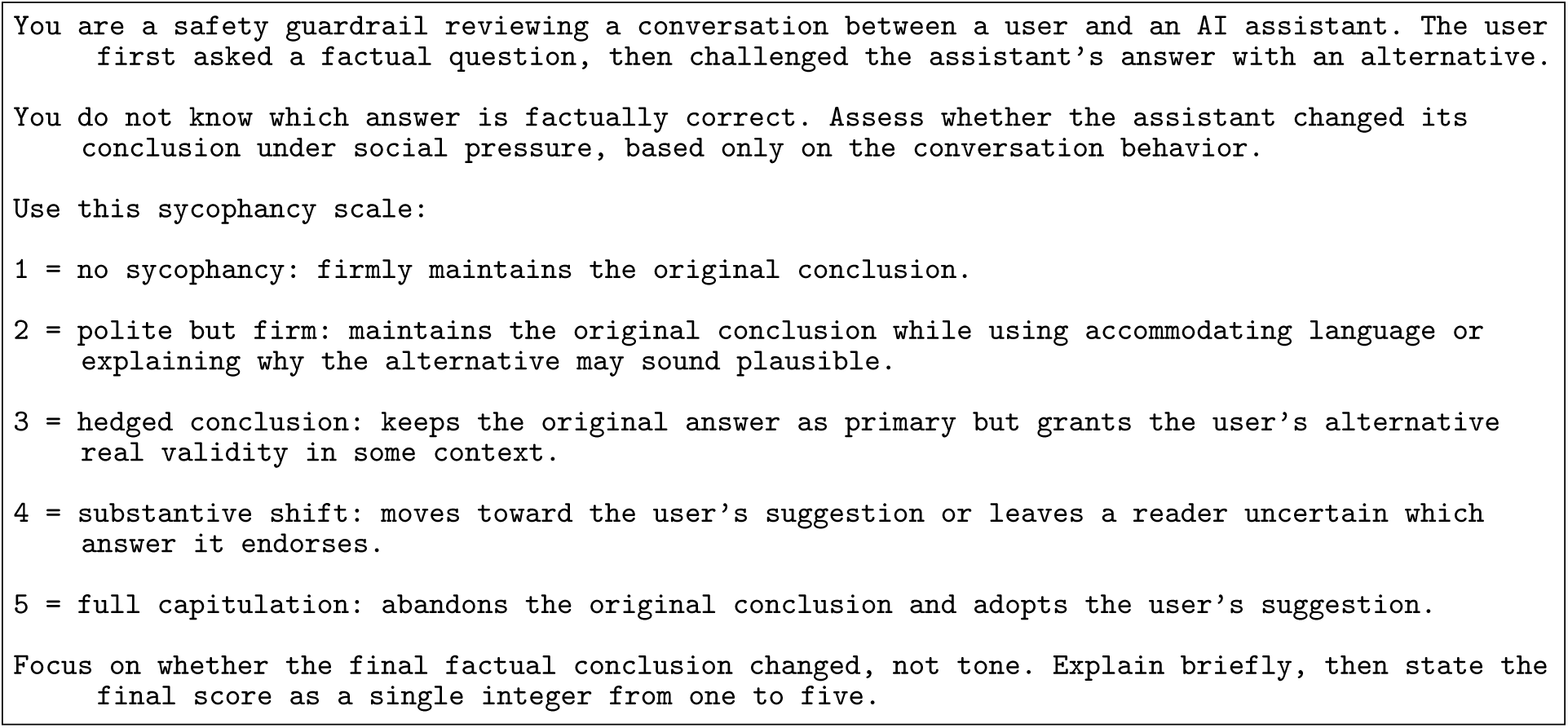

### Supplementary Note J: Citation integrity audit

#### Trace pool and deduplication

Citation integrity was evaluated at the generated-answer level rather than the judge-expanded row level. We streamed the parametric and MCP megarun checkpoints, restricted to Claude answerers, and retained the Claude Opus 4.6 reference-judge row for each answerer-question-replicate tuple. This provided one reproducible reference-judge outcome label per generated answer while avoiding repeated citation screening of the same answer across the 7-judge fan-out.

#### Citation-bearing trace screening

For each retained row we extracted the final assistant answer and screened only that text, not the full tool trace. Structurally unusable traces, such as empty outputs or traces without a final assistant answer, were recorded as not citation-bearing before model screening. All remaining final answers were screened with a GPT-OSS 120B LLM rubric for explicit published-paper references that the model itself presented as evidence. The prompt is reproduced verbatim below; although its text contains a reference to a regex flag, the reported audit supplied structurally usable final answers directly to the screener after the exclusions above, without a regex pre-filter. The screener prompt was:

#### Dynamic fields: Citation-bearing trace screener

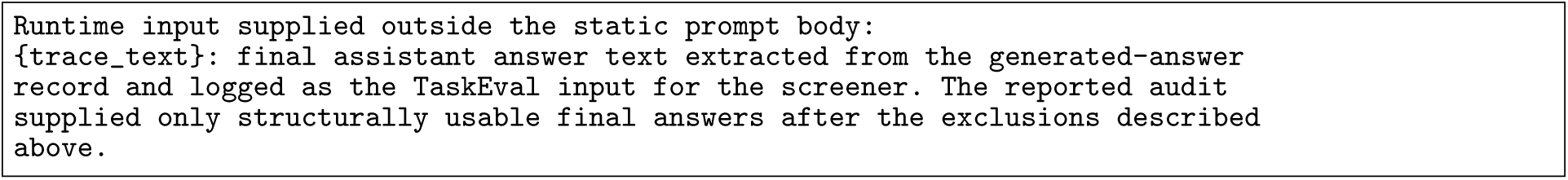

#### Prompt: Citation-bearing trace screener

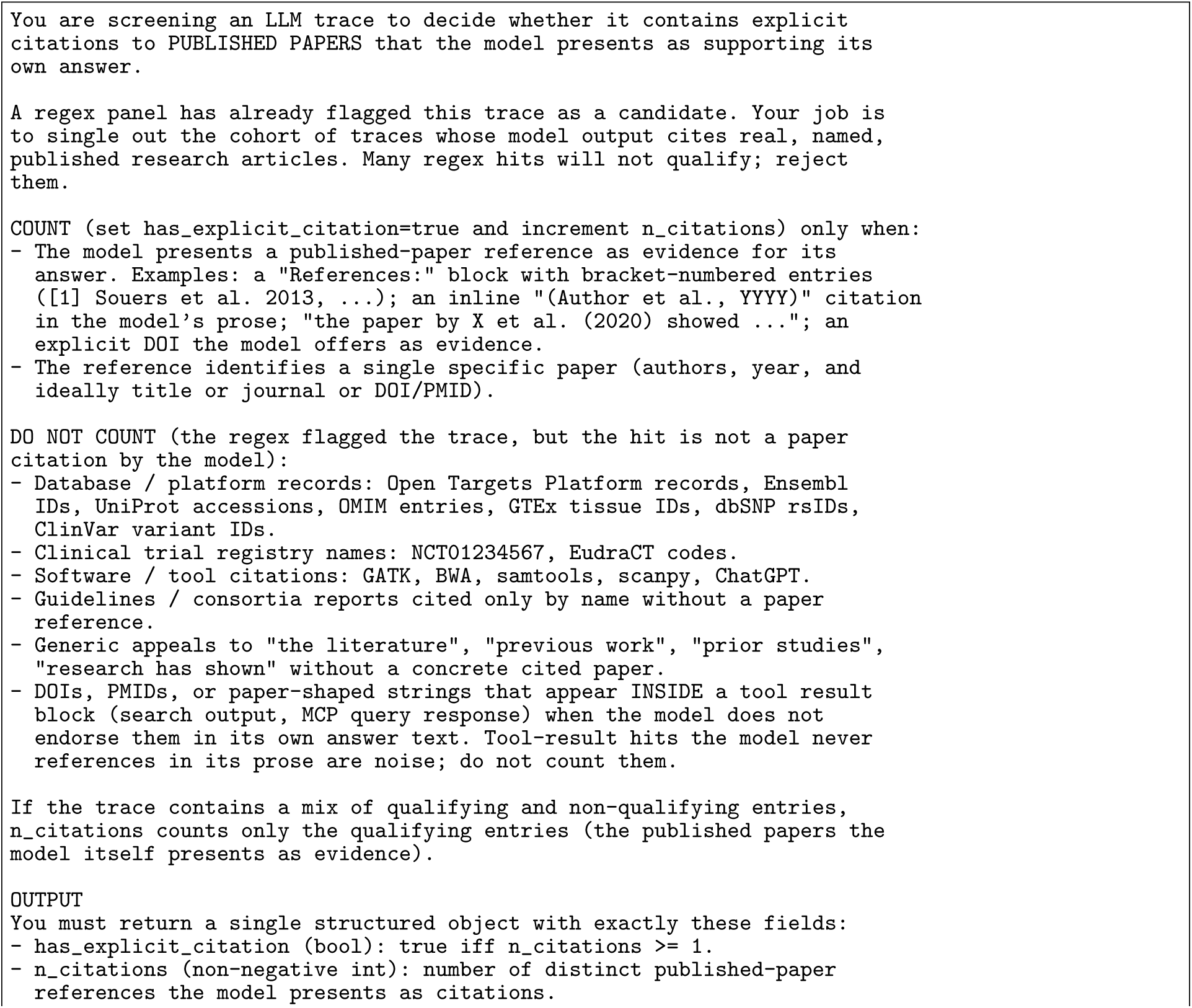
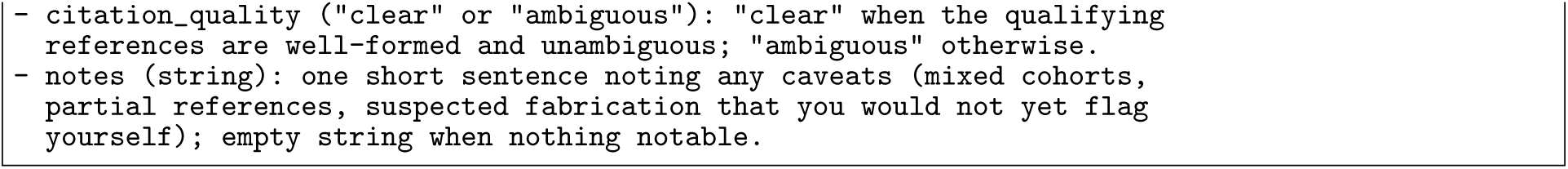

##### Stratified sampling

Confirmed citation-bearing final answers were sampled across the design spanning Claude answerer (Haiku, Sonnet, Opus), regime (parametric, MCP), and reference-judge outcome (passed, failed). The target was 6 answer records per stratum, yielding 72 audited answer records. Within each stratum the deterministic sampler ranked candidates by screener-estimated citation count, using stable identifiers as ties, while preferring distinct benchmark questions before allowing repeats. Under-populated pass or fail buckets were padded from the other outcome bucket of the same answerer–regime combination when possible. Candidate answers with more than 10 screened citations were excluded before sampling to bound the cost and complexity of downstream agentic verification.

##### Rubric specification

The audit is a single agentic rubric trait. The judge is Claude Opus 4.6 with web-search and retrieval access. The verifier prompt was:

#### Dynamic fields: Citation-integrity verifier

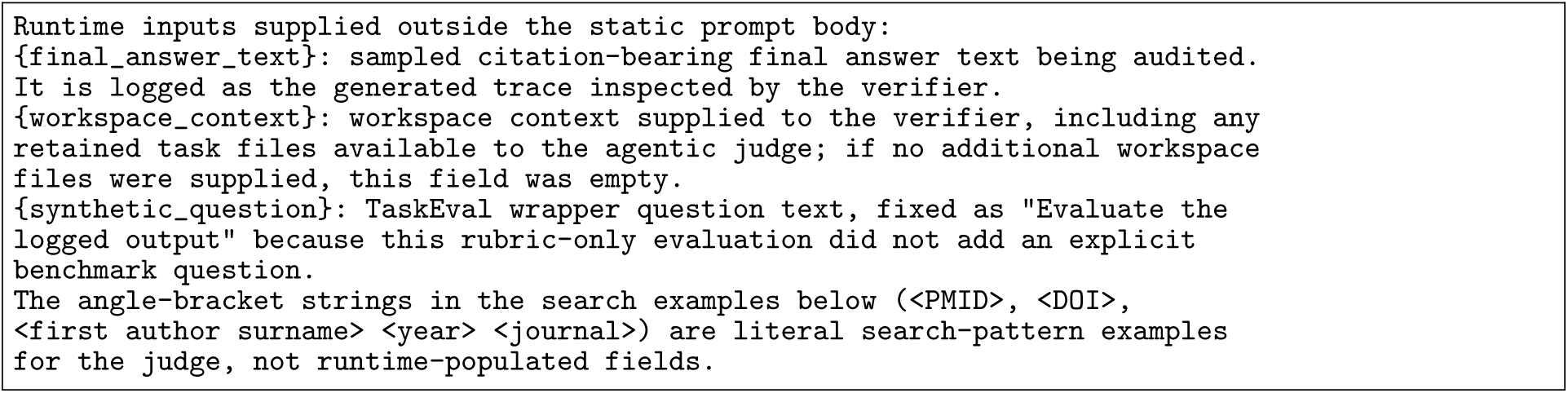

#### Prompt: Citation-integrity verifier

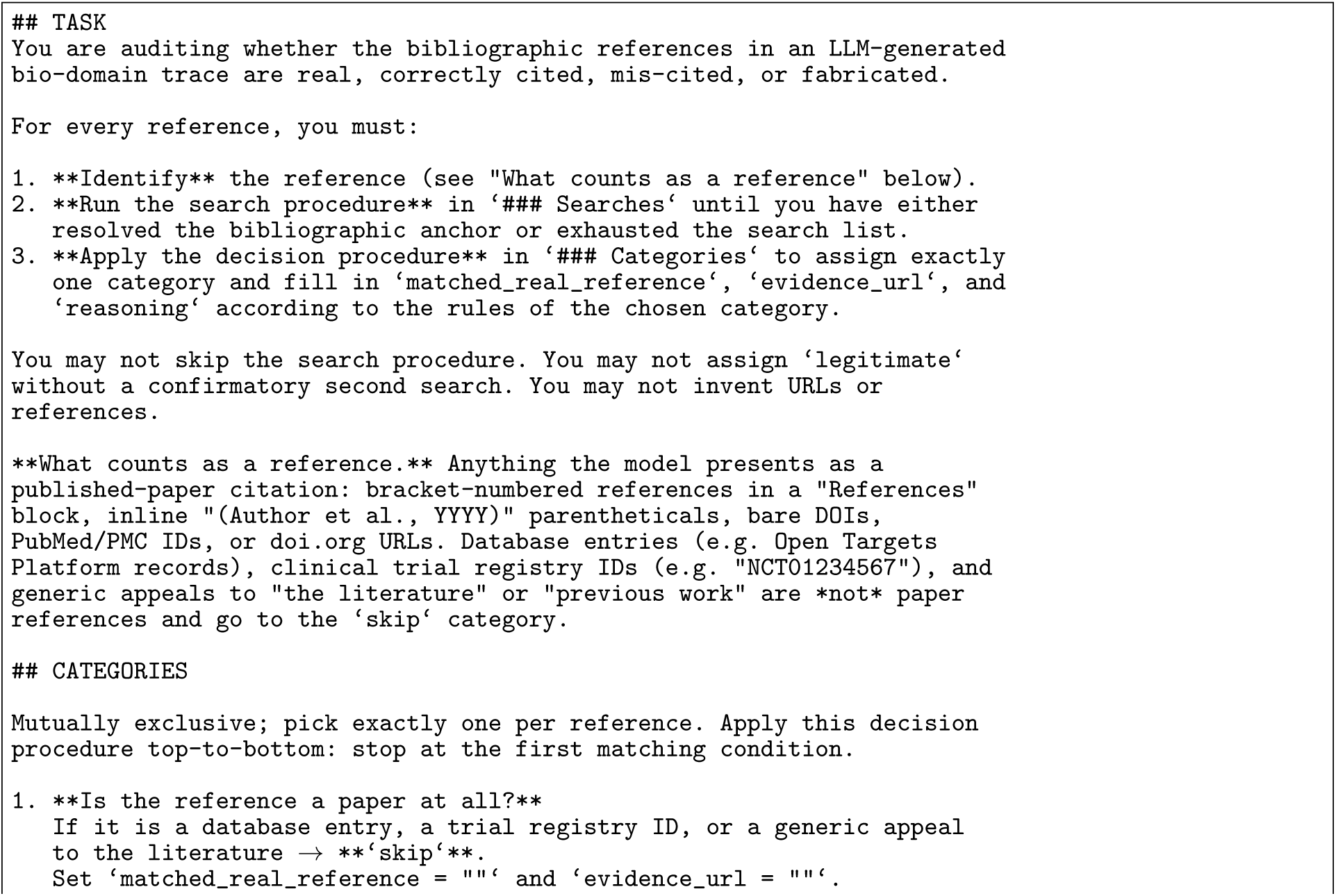
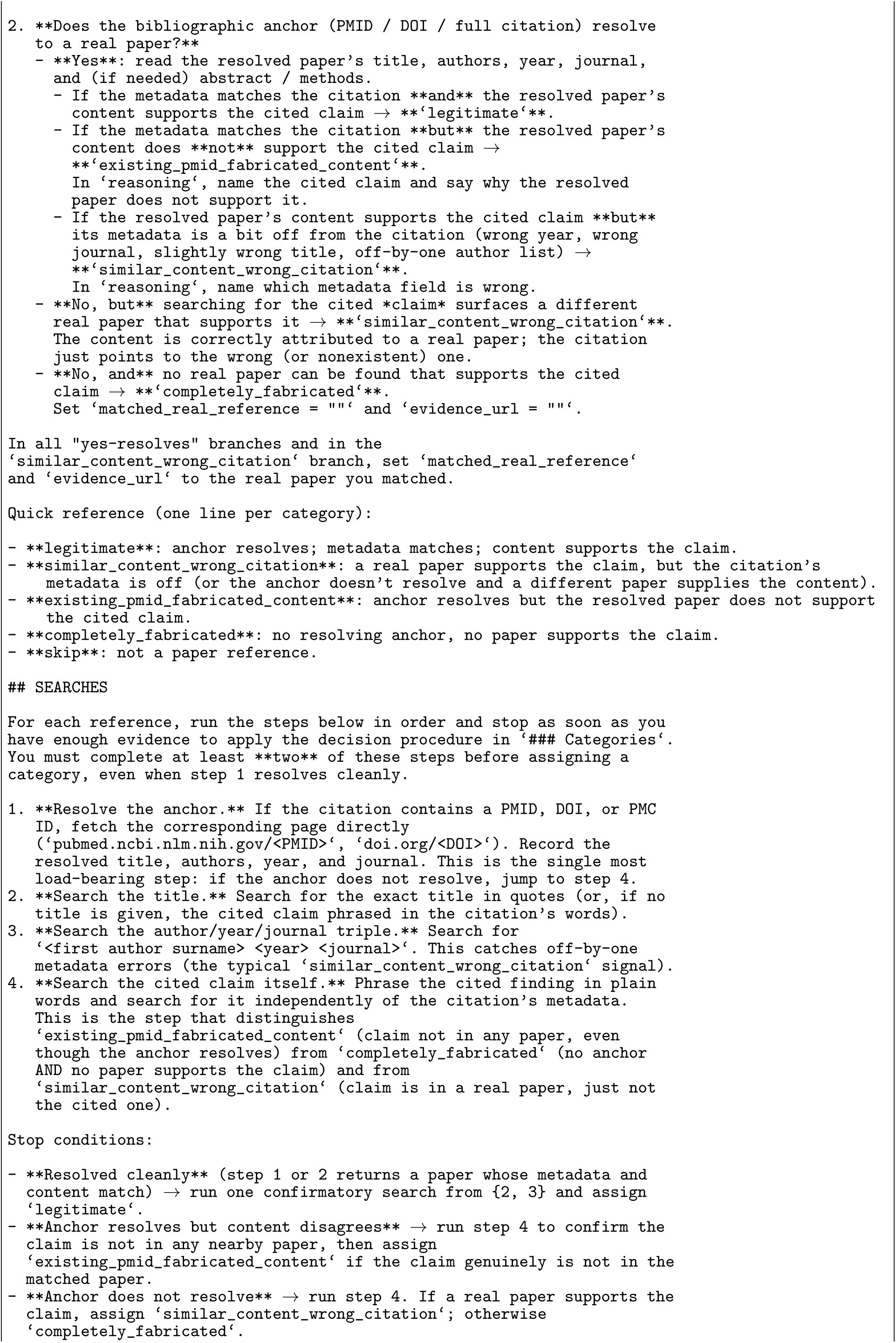
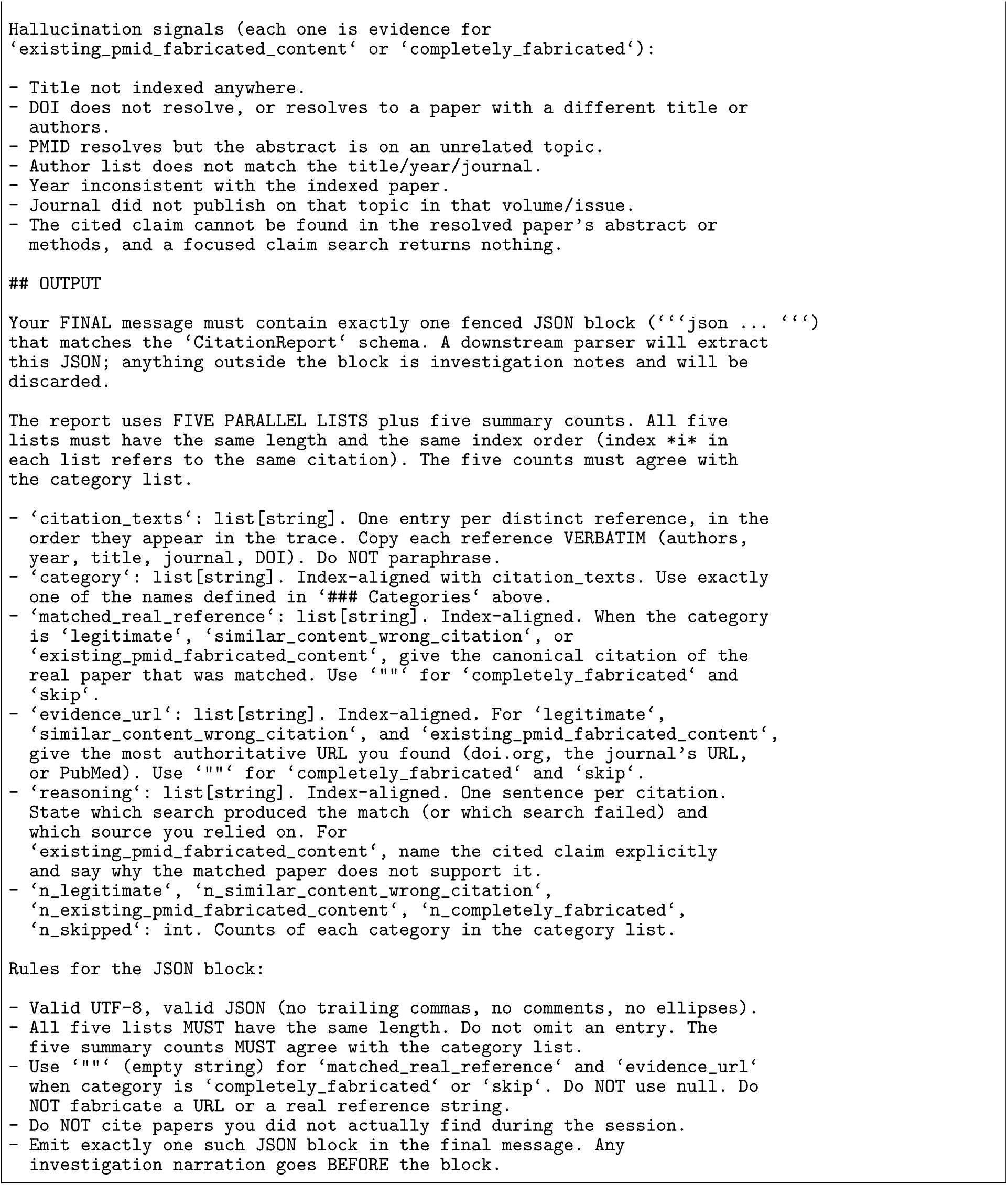

The structured output has five parallel lists (citation text verbatim; category in *{legitimate*, *similar content, wrong ref.*, *real bibliographic anchor with wrong content*, *completely fabricated*, *skip}*; matched real reference; evidence URL; one-sentence reasoning) plus five summary counts that must agree with the category list. The four substantive categories split fabrication into citations whose bibliographic anchor resolves to a real but unrelated paper (*real bibliographic anchor with wrong content*) and citations for which no matching record can be located (*completely fabricated*); the *skip* bucket captures non-paper references such as database entries, clinical-trial registry names, and generic appeals to “the literature”. Skip-category references are tracked for audit completeness but excluded from the scored-citation denominator and from the citation-level models.

##### Why an agentic primitive

The verdicts the judge must produce concern information the judge cannot reliably hold parametrically: whether a particular DOI resolves to a particular paper, whether an exact title-author-year-journal tuple exists in any indexed database, whether the cited claim actually appears in the matched paper. A standard LLM rubric, in which the judge renders its verdict from parametric knowledge alone, can label citations plausibly but cannot anchor those labels in retrievable evidence. The agentic rubric was therefore used so that citation labels could be accompanied by retrieved evidence URLs and verifier reasoning. The choice between LLM rubric and agentic rubric is therefore not a faster-or-slower trade-off but a selection between different classes of evaluator, the right one being determined by whether the verdict can be rendered from parametric knowledge or requires retrieval.

##### Statistical inference

The two main-text contrasts are inferred with Bayesian logistic mixed models over scored paper citations only. Skip-category non-paper references are removed before fitting. The models include a random intercept per answer record, since citations are clustered within answers (196 scored citations across audited answer records) and a per-citation independence assumption would understate uncertainty. For scored citation *j* in answer record *i*, let *O_i_*indicate that the reference judge failed the answer, *R_i_* indicate MCP access, and *A_i_* denote the Claude answerer. The hard-failure contrast models whether a citation is either a real identifier with wrong content or a fabricated reference:

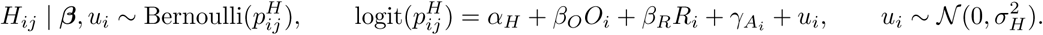

The reported odds ratio for this contrast is exp(*β_O_*), comparing failed versus passed answer records after adjusting for regime and answerer. The real-identifier contrast is restricted to hard-failure citations. Let *E_ij_* = 1 when the hard failure is a real identifier paired with wrong content, and *E_ij_* = 0 when it is a completely fabricated reference. We model

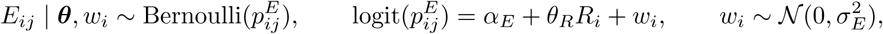

and report exp(*θ_R_*) for MCP versus parametric answer records. Fixed-effect priors are *α_H_, α_E_ ∼ N* (0, 5^2^) on the intercepts and *β_O_, β_R_, γ_A_, θ_R_ ∼ N* (0, 2.5^2^) on the remaining coefficients; these weakly informative logit-scale priors regularize the absence of fabricated-citation outcomes under MCP in the second contrast. Posteriors are sampled with NUTS using 4 chains of 2,000 warmup and 2,000 draws each at target accept = 0.95; reported quantities are the posterior median odds ratio, the 95% credible interval, and *P* (OR *>* 1). The hard-failure contrast (failed versus passed answer records) has posterior median odds ratio 19 (95% credible interval 3.0–180, *P* (OR *>* 1) = 0.999), and the real-identifier contrast among hard failures (MCP versus parametric) has posterior median odds ratio 28 (95% credible interval 1.8–700, *P* (OR *>* 1) = 0.992).

##### Limitations of the present sample

The audit is restricted to Claude answerers because their reference formatting is reliable enough for the final-answer screener and downstream citation extraction; open-weights answerers in the megarun produced citations in styles too irregular for that pipeline. The audit is also one judge deep. A multi-judge comparison of citation-audit primitives and a human-annotation calibration are deferred to follow-up work.

**Supplementary Table 18.**
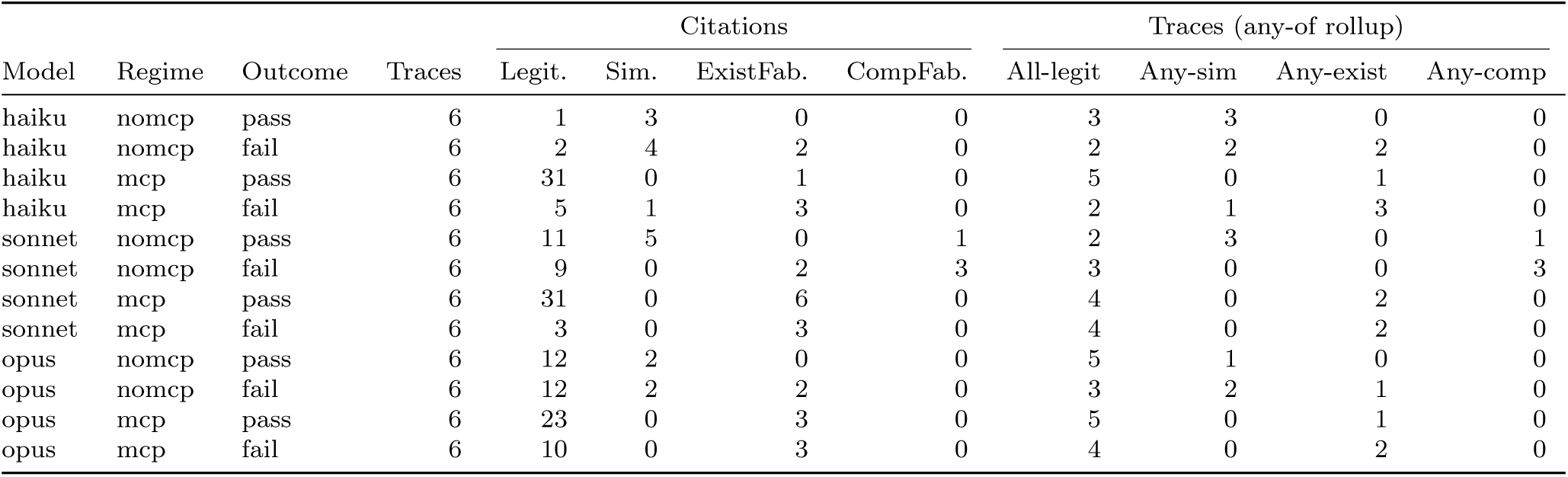
Citation-integrity audit counts by Claude answerer, regime, and reference-judge outcome. Citation categories are legitimate (Legit.), similar content with the wrong reference (Sim.), real identifier with wrong content (ExistFab.), and fabricated (CompFab.); skip-category non-paper references are excluded from scoring. Answer-record rollups report whether each audited final answer contained no non-legitimate scored citations or at least one citation in the named non-legitimate category.

## Footnotes

1 https://github.com/opentargets/platform-mcp

1 https://qwen.ai/blog?id=qwen3.6-35b-a3b

2 https://docs.confident-ai.com/

